# Structural basis for CCR6 modulation by allosteric antagonists

**DOI:** 10.1101/2024.04.05.588354

**Authors:** David Jonathan Wasilko, Brian S. Gerstenberger, Kathleen A. Farley, Wei Li, Jennifer Alley, Mark E. Schnute, Ray J. Unwalla, Jorge Victorino, Kimberly K. Crouse, Ru Ding, Parag V. Sahasrabudhe, Fabien Vincent, Richard K. Frisbie, Alpay Dermenci, Andrew Flick, Chulho Choi, Gary Chinigo, James J. Mousseau, John I. Trujillo, Philippe Nuhant, Prolay Mondal, Vincent Lombardo, Daniel Lamb, Barbara J. Hogan, Gurdeep Singh Minhas, Elena Segala, Christine Oswald, Ian W. Windsor, Seungil Han, Mathieu Rappas, Robert M. Cooke, Matthew F. Calabrese, Gabriel Berstein, Atli Thorarensen, Huixian Wu

## Abstract

The CC chemokine receptor 6 (CCR6) is a potential target for chronic inflammatory diseases such as psoriasis and inflammatory bowel disease. Previously, we reported an active CCR6 structure in complex with its cognate chemokine CCL20, revealing the molecular basis of CCR6 activation mediated by CCL20. Here, we present two inactive CCR6 structures determined by cryo-EM in ternary complexes with different allosteric antagonists, CCR6/SQA1/OXM1 and CCR6/SQA1/OXM2. OXM1 and OXM2 are oxomorpholine (OXM) analogues which are highly selective for CCR6 and disrupt the molecular network critical for receptor activation by binding to an extracellular allosteric pocket within the transmembrane domain. A U-shaped conformation stabilized by intramolecular interactions was revealed by structural and NMR studies of active OXM analogues. SQA1 is a squaramide (SQA) derivative with close-in analogues that were previously reported to be antagonists of CCR6 and other chemokine receptors. Our structures reveal an intracellular pocket occupied by SQA1 that overlaps with the G protein binding site. In addition, SQA1 stabilizes a closed conformation of the intracellular pocket, a hallmark of the inactive state of GPCRs. Minimal communication was found between the two allosteric pockets. Overall, our work provides new evidence of the versatility of GPCR antagonism by small molecules, complementing previous knowledge on CCR6 activation, and sheds light on drug discovery approaches to target CCR6 for autoimmune disorders.

## Introduction

The chemokine receptors (CKRs) are a group of class A G protein-coupled receptors (GPCRs) that respond to a family of cytokines known as chemokines^1^. More than twenty distinct CKRs have been discovered in humans, including 18 conventional CKRs that are typically G_i_-coupled and 5 atypical CKRs which are structurally related to the conventional receptors but are non-signaling^2^. CKRs can also be classified by their primary type of chemokine ligands as CC, CXC, CX_3_C, and C receptors, where C refers to the conserved N-terminal cysteines and X represents the amino acids between the two N-terminal cysteines of the chemokine^1^. Upon activation by chemokines, CKRs mediate a wide range of physiological processes including embryonic development, immune cell proliferation, activation, differentiation, and cell death. Consequently, CKRs play a significant role in the development and homeostasis of the immune system, and are important drug targets for cancer, infectious diseases, inflammation, and autoimmune disorders^3,4^.

The CC chemokine receptor 6 (CCR6)^5^ is a member of the family expressed on a variety of lymphocytes including T cells, particularly the T helper 17 (Th17) cells^6^, B cells, and dendritic cells. CCR6 is activated by its specific ligand CCL20^7^, which stimulates the migration of CCR6^+^ lymphocytes to sites of inflammation. The expression of both CCL20 and CCR6 is upregulated in the epithelial tissues of patients with chronic inflammatory conditions such as psoriasis^8^ and inflammatory bowel disease (IBD)^9^. Therefore, inhibiting CCL20-mediated CCR6 signaling is a promising strategy for anti-inflammatory drug discovery^10^. Indeed, a monoclonal antibody targeting CCL20 has resulted in the effective alleviation of inflammation in preclinical models^11^. In addition, CCR6 knockout mice are protected from induced psoriasis-like inflammation^12^. Accordingly, a small molecule antagonist of CCR6 could be useful as an oral treatment for autoimmune disorders.

Previously, we reported an active CCR6 structure in complex with CCL20 and G protein^13^. This structure revealed a shallow CCL20 pocket formed mainly by extracellular loops and the N-terminus of CCR6, leaving the orthosteric pocket within the seven-transmembrane (7TM) domain largely unoccupied. The CCL20-CCR6 recognition mode as well as the large extracellular pocket of this protein receptor pose significant challenges to the development of small molecule antagonists targeting the orthosteric site. Alternatively, modulating CKRs via allosteric ligands, notable for its versatility and diversity of mechanisms-of-action (MoA), has been an attractive and promising approach in CKR drug discovery^14–18^.

In this work, we present two inactive CCR6 structures in ternary complexes with different allosteric antagonists from squaramide (SQA) and oxomorpholine (OXM) series. The structures were determined using single-particle cryogenic electron microscopy (cryo-EM) facilitated by an engineered fiducial marker. The SQA and OXM analogues occupy two different pockets, binding of which can occur simultaneously on CCR6. Different mechanisms to allosterically antagonize CCR6 are revealed for SQA and OXM analogues, highlighting the diversity of GPCR allosteric modulation by small molecules.

## Results and Discussion

### Additive stabilization of CCR6 by SQA and OXM

SQA1 (**Fig. 1a**) is a SQA derivative that was initially discovered as a CXCR2 inhibitor and later found to also inhibit CCR6 activity^19^. Conversely, the OXM analogues OXM1 and OXM2 (**Fig. 1a**) were discovered in a ligand-based virtual screen using SQA1 as the seed and subsequent structure-activity relationship (SAR) studies. All three small molecules demonstrate robust inhibition of CCL20-mediated chemotaxis in primary human T cells expressing endogenous CCR6 (**Fig. 1b**). Significant CCR6 stabilization was observed in a thermal shift assay (**Fig. 1c**) upon incubation with the small molecules, confirming direct target engagement for the SQA and OXM molecules. Surprisingly, the stabilization of CCR6 by analogues from the two series is additive, as a significantly larger thermal shift was observed when analogues from both series were added to the receptor. Such additive stabilization was not dependent on the order of ligand addition between SQA1 and OXM analogues but was not observed when CCR6 was co-incubated with the two different OXM molecules. These results suggest that despite the discovery of the OXM series via a SQA-based virtual screen, the two series bind to different pockets on CCR6 and can bind independently and simultaneously to the receptor.

**Figure 1.**
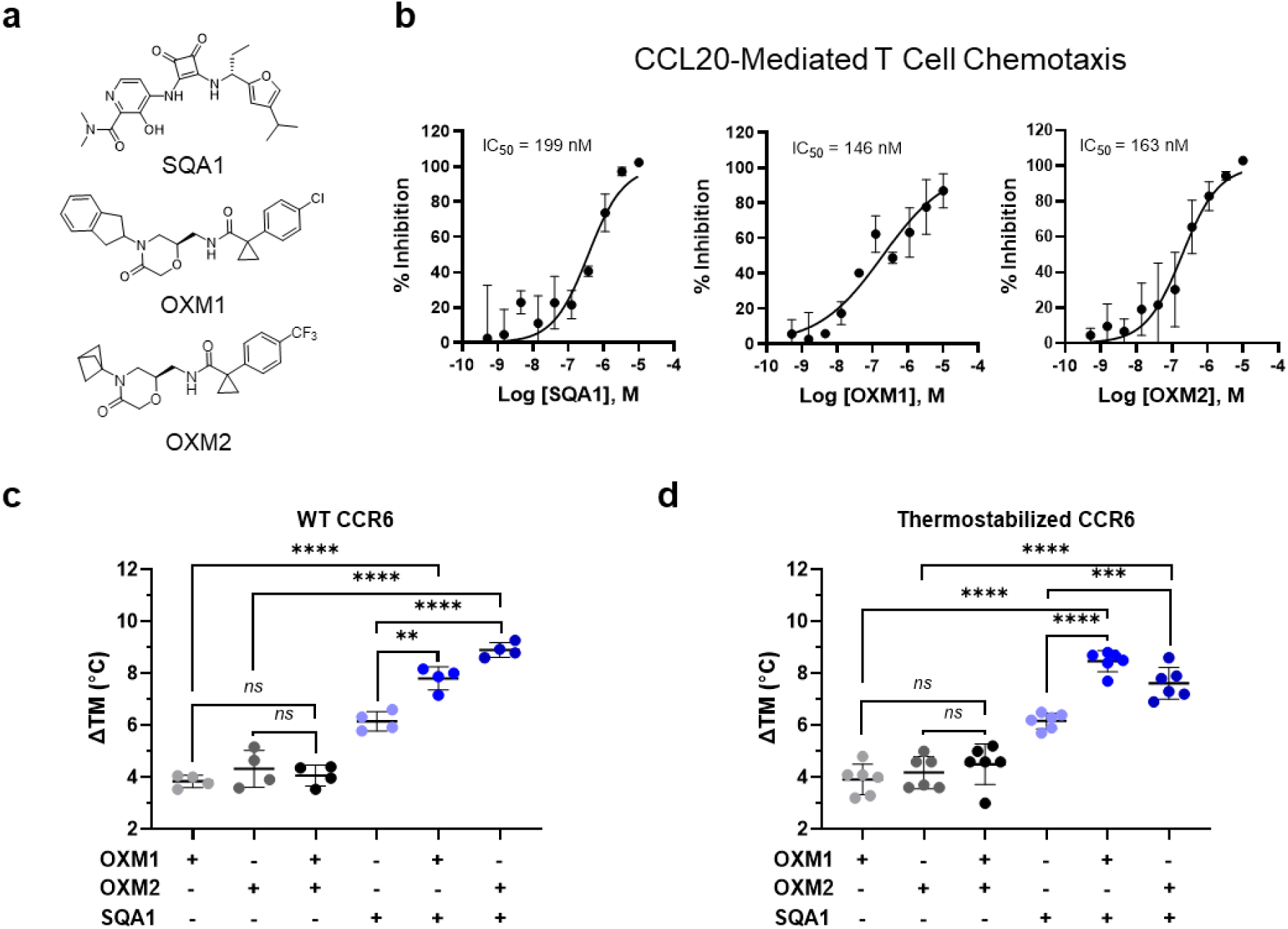
SQA and OXM analogues inhibit CCL20-mediated activation of CCR6 and display additive binding stabilization to the receptor. **a**, Chemical structures of SQA1, OXM1 and OXM2. **b**, Inhibition of CCL20-mediated CCR6^+^ human T-cell chemotaxis demonstrated by SQA1 (IC_50_ = 199 nM or pIC_50_ = 6.7 ± 0.8, mean ± s.d., *n* = 6), OXM1 (IC_50_ = 146 nM, or pIC_50_ = 6.8 ± 0.2, mean ± s.d., *n* = 4), and OXM2 (IC_50_ = 163 nM, pIC_50_ = 6.78 ± 0.11, mean ± s.d., *n* = 2). **c-d**, Thermal shifts of **c**, WT (*n* = 4) and **d**, thermostabilized (*n* = 6) CCR6_Nα7.1 from Apo to liganded conditions as indicated. Statistical analysis was done by unpaired *t* test (*ns*, not significant; ** *p* < 0.01; *** *p* < 0.001; **** *p* < 0.0001). Data for graphs **b-d** are available as source data.

### Construct engineering for cryo-EM studies

To reveal how the SQA and OXM analogues bind to CCR6, we conducted cryo-EM studies using an engineered CCR6 construct Nα7.1 which contains seven thermostabilizing alanine-substitutions^20,21^ in the transmembrane domain and a point mutation L134^3.41^W (superscripts indicate Ballesteros–Weinstein numbering for GPCRs^22^) to increase receptor expression^23^. Binding of the SQA and OXM analogues to the thermostabilized CCR6 construct was confirmed with comparable potency to the wild-type (WT) receptor (**Fig. S1**). Additive stabilization between the SQA and OXM analogues was also observed with the stabilized construct (**Fig. 1d**). For structural studies, twenty-seven residues at both N and C-termini of CCR6 were truncated to improve protein homogeneity. Furthermore, the residues between L241 and K251 of intracellular loop 3 (ICL3) were replaced by a thermostabilized apocytochrome b562 (BRIL)^24^. Two amino acid linkers were grafted from the A_2A_ adenosine receptor^25^ to stabilize continuous alpha-helices as the junctions between TM5 (CCR6) and helix 1 (BRIL), as well as TM6 (CCR6) and helix 4 (BRIL). The engineered CCR6-BRIL chimeric construct (**Fig. S2**) was co-purified with analogues from both SQA and OXM series. An anti-BRIL Fab^26^ with an anti-Fab nanobody (Nb)^27^ was employed to complex with the BRIL in the chimeric CCR6 construct. The BRIL/Fab/Nb module was introduced as a fiducial marker in the single-particle cryo-EM analysis to facilitate particle alignment^28^. This approach led to the successful determination of two inactive CCR6 structures bound by ligands from the SQA and OXM series.

The complex structures of CCR6/SQA1/OXM1 and CCR6/SQA1/OXM2 were determined at overall resolutions of 2.63 Å and 3.02 Å, respectively (**Figs. S3-4, Table S1**). Cryo-EM reconstructions of both complexes reveal robust density, allowing the unambiguous fitting of BRIL, Fab, Nb, and the 7TM of CCR6 (**Figs. 2a, b**). Most extracellular loops (ECLs) and intracellular loops (ICLs) are unresolved or ambiguous in the reconstruction except for the bottom of ECL2, which adopts a β-hairpin structure like that reported in the active CCR6 and other CKR structures^13^. The disordered loops in the final reconstructions are not unexpected based on the known intrinsic flexibility of these regions in GPCRs. In both structures, the densities of the SQA and OXM analogues are robustly resolved, with each series binding to a distinct but well-defined pocket within the 7TM bundle, allowing confident modeling of ligand binding for subsequent structural analysis (**Figs. 2, S3-4**).

**Figure 2.**
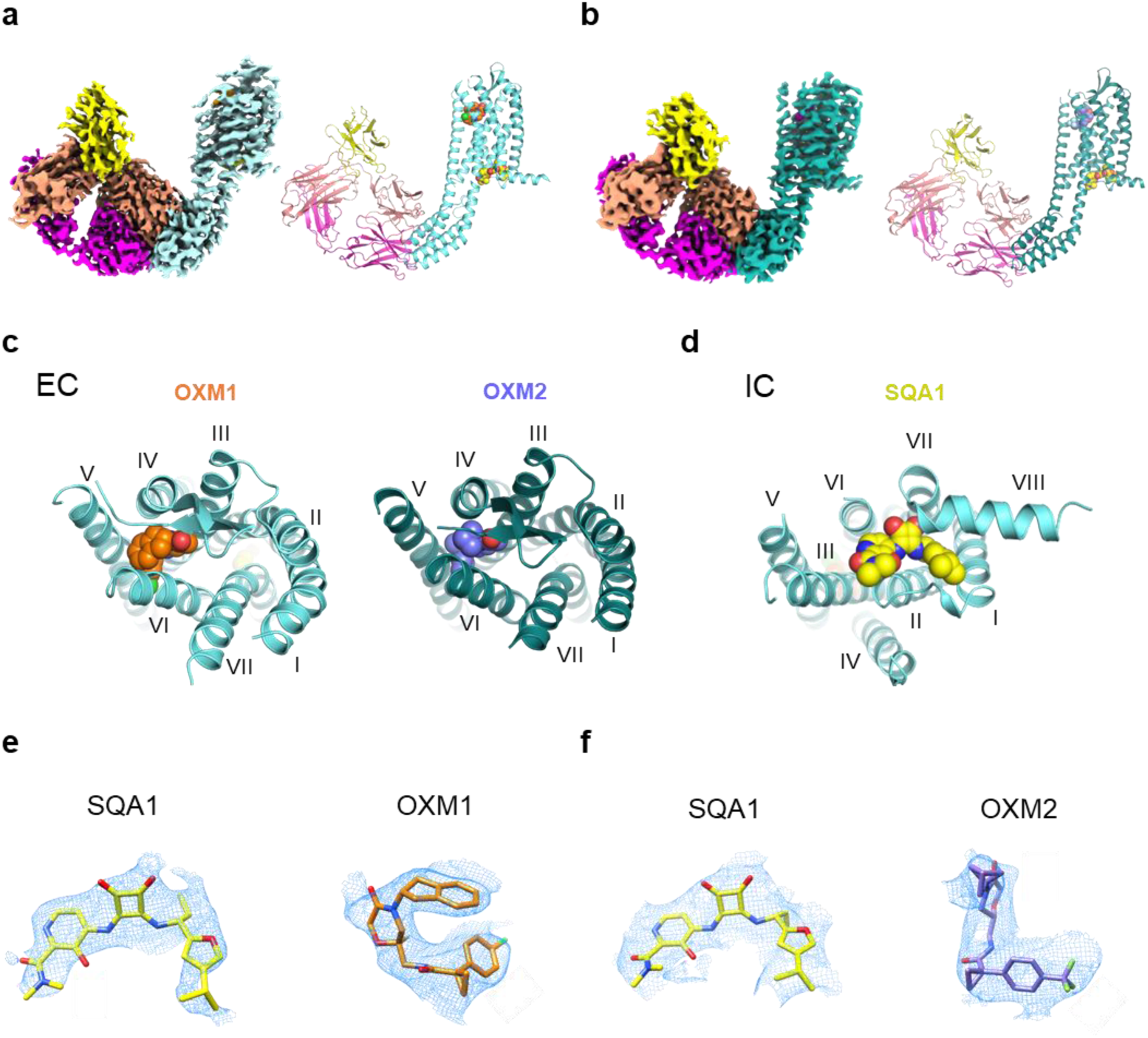
Overall structures of CCR6 in complex with OXM and SQA analogues. **a-b**, Cryo-EM map (left) and model (right) of CCR6 complex with anti-BRIL Fab and anti-Fab Nb bound to **a**, OXM1 (orange carbon spheres) and SQA1 (yellow carbon spheres), or **b**, OXM2 (slate carbon spheres) and SQA1 (yellow carbon spheres). Protein is colored by subunit as follows: Fab heavy chain, magenta; Fab light chain, salmon; Nb, yellow; CCR6, aquamarine in **a** and deep teal in **b**. **c**, Extracellular (EC) view of CCR6 bound to OXM analogues, OXM1 (left) and OXM2 (right). **d**, Intracellular (IC) view of CCR6 bound to the SQA analogue, SQA1. Both **c** and **d** use the same color code as **a** and **b**. **e**, SQA1 and OXM1 fit into the cryo-EM density map. **f**, EM density for SQA1 and OXM2 fit into the cryo-EM density map.

### The OXM analogues bind to an extracellular allosteric pocket

U-shaped density is resolved in an extracellular pocket for both reconstructions and can be fitted by the two different OXM analogues, OXM1 and OXM2 (**Figs. 2a, b**). Binding of the OXM analogues to CCR6 is mainly mediated by hydrophobic interactions with side chains from TMs 3, 4, 5, 6, and ECL2 (**Figs. 2c, 3a-d**). In particular, the *p*-chlorophenyl group of OXM1 and the *p*-trifluoromethylphenyl moiety of OXM2 are sandwiched between TMs 5 and 6 with their *para*-substitutions pointing towards the hydrophobic membrane, while making hydrophobic interactions with F129^3.36^, L219^5.43^, F223^5.47^, I268^6.49^, M272^6.53^, and L275^6.56^. On the other end of the bent molecules are hydrophobic substitutions on the amide nitrogen of the oxomorpholine core, i.e., an indane moiety in OXM1 and a bicyclo[1.1.1]pentane in OXM2. These two groups bind in a hydrophobic pocket capped by the C-terminal tip of ECL2 that connects to TM5. Due to the compact size of this subpocket, the larger indane group of OXM1 bends down towards the *p*-chlorophenyl group, forming an intramolecular T-shaped π stacking interaction which likely further stabilizes the observed U-shaped conformation of OXM1. Other than the hydrophobic interactions, several H-bonds are mediated by the two amide groups of the OXM analogues with polar side chains such as N130^3.37^, K122^3.29^, T182^4.60^, and N271^6.52^. Remarkably, a majority of the OXM binding pocket side chains are CCR6-specific (**Fig. S5**). Consequently, OXM1 was found to be highly selective for CCR6 and is not active against other CKRs (**Table S2**).

**Figure 3.**
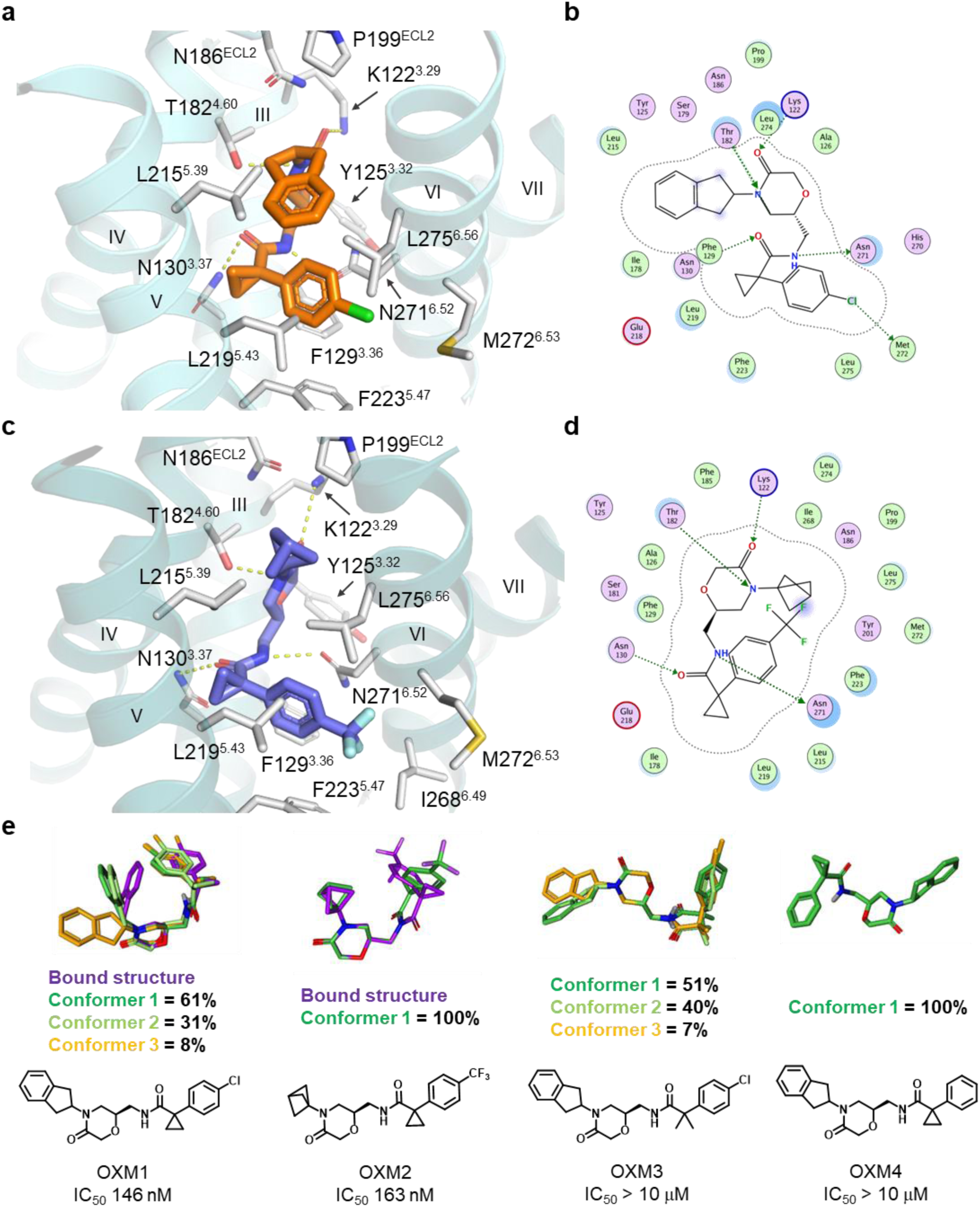
Structures of OXM analogues bound to CCR6 and comparison to solution conformations. **a**, Binding pocket of OXM1 (orange carbon sticks). **b**, Protein-ligand interaction diagram for OXM1 prepared with Molecular Operating Environment (MOE)^36^. **c**, Binding pocket of OXM2 (slate carbon sticks). **d**, Protein-ligand interaction diagram for OXM2 prepared with MOE^36^. In **a** and **c**, CCR6 protein is shown in aquamarine and deep teal ribbons, respectively. Side chains interacting with the small molecules are shown as white carbon sticks. Hydrogen bonds are highlighted by yellow dashed lines. In **b** and **d**, OXM pocket residues are presented as follows: polar residues in pink, hydrophobic residues in green, acidic residues with a red contour ring, basic residues with a blue contour ring. Green dotted arrows indicate hydrogen and halogen bonds mediated by side chains that contribute to ligand binding. Ligand atoms exposed to environment are shaded in blue according to degrees of exposure, scaled by size. Light-blue halos around residues indicate the degree of interaction with ligand, scaled by size. The dotted contour around the ligand reflects steric room for methyl substitution. **e**, Chemical structures of different OXM analogues and their solution conformations were determined by NMR using residual dipolar couplings (RDC)^30^. Three most populated conformations are illustrated by dark green, light green, and orange carbon sticks, respectively. The bound conformations of OXM1 and OXM2 are shown as purple carbon sticks in the overlays for comparison. The IC_50_ value of each OXM analogue determined in the CCL20-mediated human CCR6^+^ T-cell chemotaxis assay is shown under the corresponding chemical structure.

### Conformational restraints required for OXM activity

The observed bent pose of OXM1 in the ligand-bound structure prompted us to study the solution conformation of this molecule using Nuclear Magnetic Resonance (NMR) experiments to understand the energy landscape of this conformation. Rotating-frame nuclear Overhauser Enhancement signals (ROEs) were collected using previously reported methods^29^. Robust ROEs were obtained between the indane C(sp^3^)-*H* (H-18) and the aromatic hydrogens from the *p*-chlorophenyl ring (Hs-3,4,6,7) (**Fig. S6**), indicating intramolecular interactions between the indane and the *p*-chlorophenyl moieties are also maintained in solution. Additionally, we performed proton and carbon assignments for OXM1 (**Figs. S7-13**, **Table S3**) in a compressed dimethyl sulfoxide (DMSO) gel^30,31^ which were used to calculate the residual dipolar coupling (RDC) values of the proton-carbon and proton-nitrogen J-couplings. The RDC data was fit to a conformational ensemble that consisted of low energy structures generated by molecular dynamics simulations in a solvent-dependent manner (see methods). This exercise output several U-shaped conformations of OXM1 as the best fits. Ninety-two percent of the low energy conformers greatly resemble the bound pose observed in the cryo-EM structure (**Fig. 3e**), suggesting a low energy conformation for the bound state.

We rationalize the conformational restraints conferred by the cyclopropyl group as well as the T-shaped π stacking interactions between the indane and the *p*-chlorophenyl groups are critical factors in stabilizing the observed U-shaped conformation of OXM1 in both bound and free forms. To test this hypothesis, we synthesized and analyzed two additional OXM analogues, OXM3 which bears a *gem*-dimethyl replacement at the cyclopropyl position, and OXM4 that lacks the *para*-Cl substitution on the phenyl ring (**Fig. 3e**). As expected, extended conformations in solution are observed for the two new OXM analogues and no long-range ROEs or nuclear overhauser enhancement signals (NOEs) were observed (**Figs. 3e, S14-27, Table S3**). Consequently, neither OXM3 nor OXM4 are active on CCR6. Finally, we also characterized the solution conformation of OXM2 used in the cryo-EM studies. Despite the absence of the indane moiety, a curved conformation that largely resembles the bound conformation was revealed in its solution state (**Fig. 3e, S28-34, Table S3**). The curved conformation of OXM2 in solution was further confirmed by long-range NOEs observed between the bicyclo[1.1.1]pentane (Hs-21,24,25) and the aromatic hydrogens from the *p*-trifluoromethyl phenyl ring (Hs-16,17,19,20) (**Fig. S35**). In sum, the NMR studies support the hypothesis that a low-energy curved conformation in the free state is critical for OXM analogues to be active on CCR6.

### SQA1 binds to an intracellular pocket

In contrast to the OXM analogues, an intracellular binding site is revealed for the SQA molecule SQA1 (**Figs. 2d, 4**). The binding poses of SQA1 from both structures are nearly identical. Therefore, the CCR6/SQA1/OXM1 complex that led to a higher-resolution reconstruction was used for analysis hereafter. Side chains interacting with SQA1 are from TMs 1, 2, 3, 6, 7, and helix 8 (**Figs. 4a, b**). CCR6 has a glycine residue G320^8.47^ as the TM7-helix 8 linker. The absence of a side chain at the 8.47 position provides a pocket that allows the squaramide core to bind. Indeed, a CCR6 mutant bearing G320^8.47^V completely abolished SQA1 binding (**Figs. 4c, d**). Extensive polar interactions are observed between the squaramide-picolinamide core of SQA1 and CCR6, including the salt bridges between D82^2.40^ and the squaramide NHs, as well as H-bonds mediated by side chains of S79^2.37^, T81^2.39^, R143^3.50^, and K322^8.49^ with the 3’-hydroxyl-picolinamide moiety. Consequently, SQA1 binding was abolished on mutants bearing single point mutations such as D82^2.40^N, S79^2.37^A, T81^2.39^L, R143^3.50^A, or K322^8.49^A (**Figs. 4c, d**). On the other hand, binding of the 1-(4-isopropyl furan-2-yl)propyl moiety is largely mediated by hydrophobic interactions with side chains from TMs 1, 2, 7, and helix 8. Particularly, the 1-propyl binds in a hydrophobic pocket formed by the side chains of L85^2.43^, Y316^7.53^, and V67^1.53^. CCR6 mutants bearing mutations in this pocket such as L85^2.43^A or Y316^7.53^A no longer respond to SQA1 (**Figs. 4c, d**).

**Figure 4.**
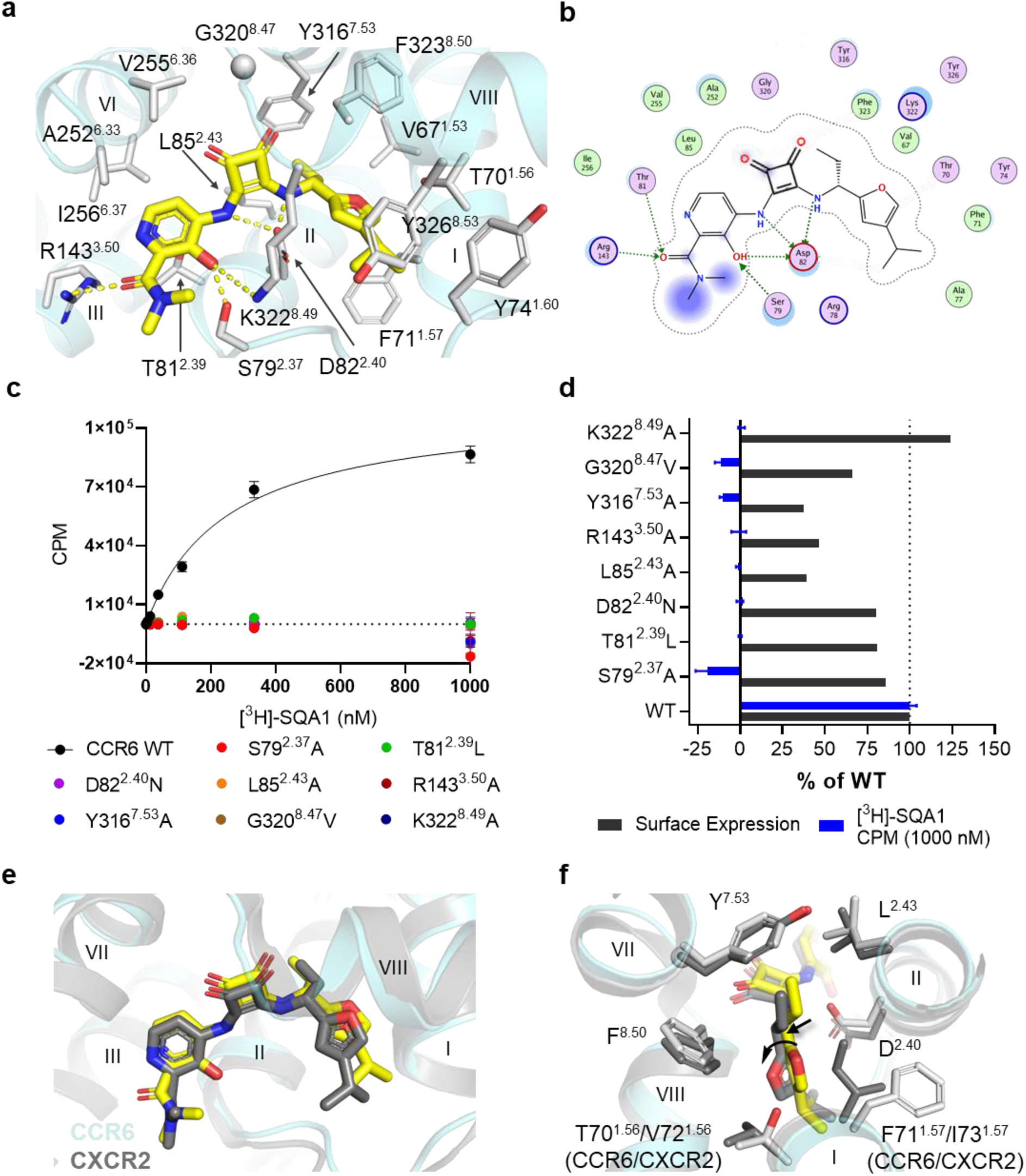
SQA1 binds to an intracellular pocket. **a**, Structure of SQA1 (yellow carbon sticks) bound to CCR6 (aquamarine ribbon) from the higher-resolution CCR6/SQA1/OXM1 cryo-EM structure. Side chains interacting with SQA1 are shown by white carbon sticks. Hydrogen bonds are highlighted by yellow dashed lines. **b**, Protein-ligand interaction diagram for SQA1 prepared with MOE^36^. Definitions of the schematic representation are same as that in Fig. 3b and d. **c**, Saturation binding curves of [^3^H]-SQA1 on CCR6 WT and mutants as indicated. Binding of [^3^H]-SQA1 on WT provided a K_D_ of 250 ± 22 nM. Data shown as mean ± s.d. are representative of three independent experiments (*n* = 3). No significant binding was observed at all doses measured on the CCR6 mutants. **d**, Normalized surface expression (dark grey bars) and CPM (counts per minute) of [^3^H]-SQA1 measured at 1,000 nM ligand concentration (blue bars) for CCR6 mutants transiently expressed in HEK293 cells. Data of WT was used as the baseline for normalization calculation. **e**, Overlay of CCR6 and CXCR2 (PDB ID 6LFL) inactive structures bound by SQA1. **f**, Major differences of the binding pockets in CCR6/- and CXCR2/SQA1-bound structures. Side chains of pocket residues are shown in sticks. In **e** and **f**, CXCR2 and SQA1 from 6LFL are shown by dark grey ribbon and dark grey sticks. CCR6 structure is shown by the same cartoon representations as in **a**. Data for graphs **c-d** are available as source data.

A co-crystal structure of CXCR2 in complex with SQA1 was reported previously^32^. Comparison of the bound structures of SQA1 on CCR6 and CXCR2 reveal a highly similar mode adopted by the molecule in binding to a mostly conserved pocket shared by the two receptors (**Figs. 4e, S36**). As observed in CCR6, extensive polar interactions are also seen in the CXCR2 structure, mediated by the squaramide-picolinamide core with conserved side chains shared by the two receptors. The most striking difference in the ligand binding pose is for the 1-(4-isopropyl furan-2-yl)propyl moiety (**Fig. 4f**). In CXCR2, this moiety is pushed about 1 Å downward towards the intracellular side compared to the bound position in CCR6. Movement of the 1-(4-isopropyl furan-2-yl)propyl moiety is likely caused by the difference of side chains at the 1.57 position, which is F71^1.57^ in CCR6 and is replaced by I73^1.57^ in CXCR2. The aliphatic isoleucine side chain reaches deeper into the 1-propyl binding pocket, resulting in a smaller subpocket in CXCR2 that pushes the 1-propyl group down. Furthermore, to avoid clashing into the isoleucine side chain, the 4-isopropyl furan of SQA1 in CXCR2 undergoes a rotation away from TM1 compared to the bound structure in CCR6.

### SQA and OXM antagonize CCR6 via different allosteric mechanisms

The bound CCR6 structures are resolved in inactive conformations (**Fig. 5a**). A comparison of the SQA- and OXM-bound CCR6 structures to the previous CCL20-bound structure reveals that the binding sites of the OXM and SQA analogues do not overlap with the CCL20 binding site (**Fig. 5b**), indicating both OXM and SQA molecules are allosteric antagonists of CCR6. Compared to the active structure, a significant rearrangement of the CCR6 7TM bundle is observed in the inactive structures, which includes a 4-5 Å inward movement of the entire TM6 towards the center of the TM bundle and a 4.6-Å outward movement at the IC tip of TM7 (**Figs. 5c, d**). These changes result in a closed IC pocket that is not compatible with G protein coupling. Furthermore, the conserved functional motifs such as P226^5.50^-M133^3.40^-F263^6.44^, D142^3.49^-R143^3.50^-Y144^3.51^, and N312^7.49^-P313^7.50^-x-x-Y316^7.53^ are now resolved in conformations resembling those observed in most other inactive class A GPCR structures (**Figs. 5e-g**)^33^. In the inactive structures of CCR7^18^ and CCR9^16^ reported previously, a conserved N^6.52^-Y^3.32^-Q^6.48^ H-bond network was observed as a hallmark of the inactive state^13^. In the inactive CCR6, a similar H-bond network mediated by Y125^3.32^, Q267^6.48^, and N271^6.52^ is also observed (**Fig. 5h**). Notably, both Y125^3.32^ and N271^6.52^ are residues located in the OXM binding pocket, and the side chains of these residues mediate specific protein-ligand interactions via hydrophobic packing and a H-bond interaction with the oxomorpholine core, respectively. Therefore, binding of OXM analogues likely further stabilizes the Y125^3.32^-Q267^6.48^-N271^6.52^ H-bond network that is important for the inactive state of CCR6.

**Figure 5.**
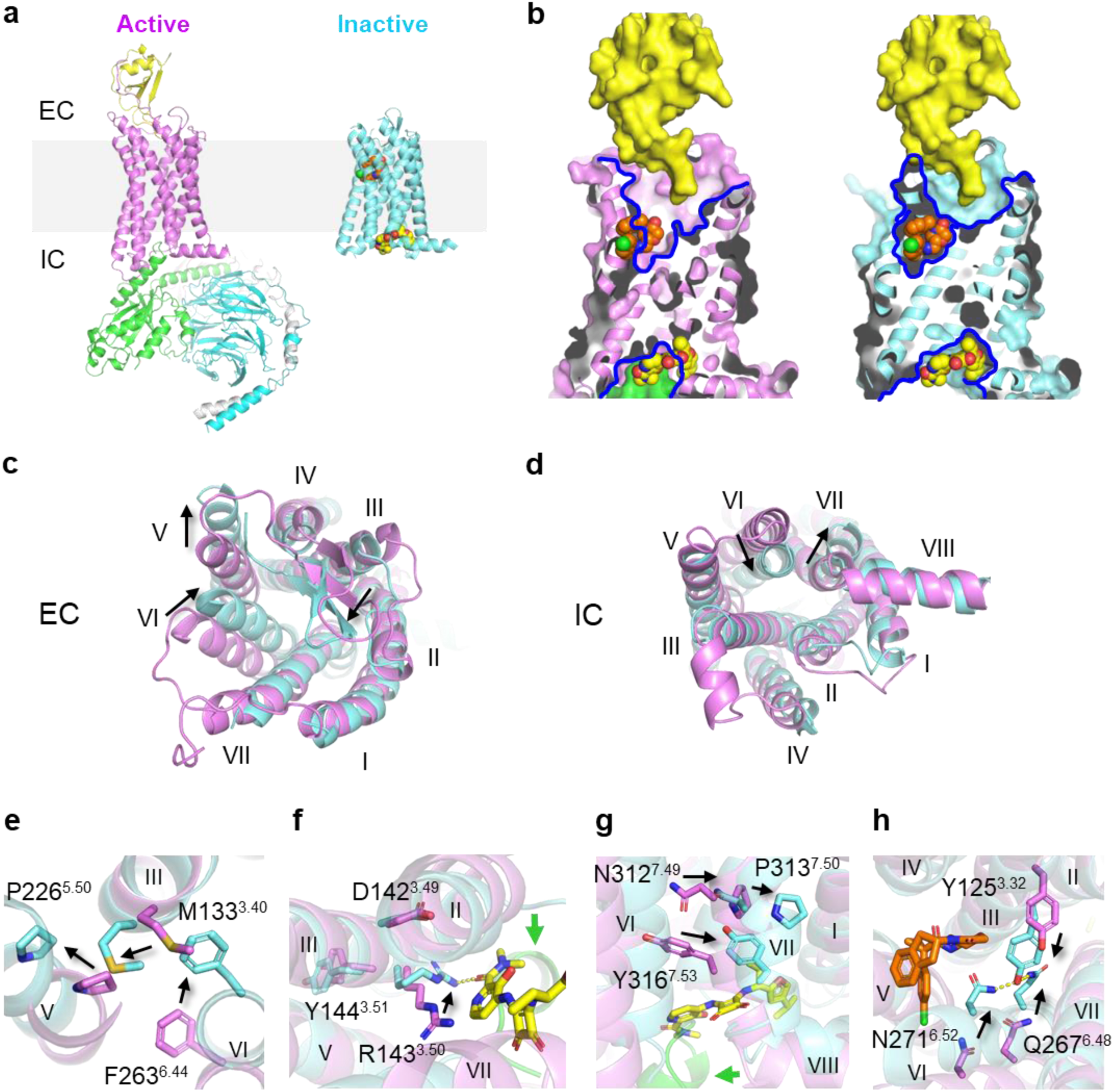
Different mechanisms of allosteric antagonism of CCR6 by OXM and SQA. **a**, Overall structures of the active CCR6 (magenta, PDB ID 6WWZ, left) in complex with its cognate chemokine ligand CCL20 (yellow) and a heterotrimeric G protein colored by subunit (Gαo, green, Gβ, cyan, and Gγ, light grey), and the inactive CCR6 (aquamarine, right) in complex with SQA1 (yellow carbon spheres) and OXM1 (orange carbon spheres). The light grey box highlights the location of cell membranes. **b**, Overlays of CCL20, OXM1, and SQA1 bound structures onto the active (left) and the inactive (right) CCR6 structures. Structures are presented in the same color codes as in **a**. Blue lines depict pockets revealed by the structures. **c-d**, Structural overlays of the active (magenta) and inactive (aquamarine) CCR6 structures from **c**, EC and **d**, IC views. **e-h**, Conformational changes of conserved functional motifs, **e**, P226^5.50^-M133^3.40^-F263^6.44^, **f**, D142^3.49^-R143^3.50^-Y144^3.51^, **g**, N312^7.49^-P313^7.50^-x-x-Y316^7.53^, and **h**, Y125^3.32^-Q267^6.48^-N271^6.52^ revealed by structural overlay of active (magenta) and inactive (aquamarine) CCR6. In **c-h**, significant TM movements and side chain conformational changes from active to inactive states are highlighted by black arrows.

Bound to an inactive IC pocket, SQA1 conformationally and sterically hinders the coupling of the G protein (**Fig. 5b**). Furthermore, SQA1 engages the side chains of R143^3.50^ and Y316^7.53^, which are part of the D^3.49^R^3.50^Y^3.51^ and the N^7.49^P^7.50^xxY^7.53^ motifs involved in G protein binding and CCR6 activation. In the SQA1 co-structures, R143^3.50^ and Y316^7.53^ are found in inactive conformations (**Figs. 5f, g**). Therefore, SQA1 binding spatially competes with G protein binding and prevents conformational changes of the TMs and the critical activation motifs, which altogether inhibits CCL20-mediated CCR6 activation. In addition to SQA1, a few other intracellular antagonists have been reported for other CKRs, such as CXCR1/2^15,34^, CCR2^17^, CCR7^18^, and CCR9^16^, likely all utilizing a highly similar molecular mechanism to regulate their corresponding receptors.

Located in the extracellular half of the 7TM, the OXM pocket is about 9 Å away from the CCL20 binding site (**Fig. 5g**). Binding of the OXM analogues, however, reshapes the orthosteric pocket of CCR6 to a conformation that is vastly different from that observed in the CCL20-bound form (**Figs. 5b, c**). Particularly, by interacting with N186 from ECL2 and the C-terminal tip of the loop, OXM1 or OXM2 binding brings the β-hairpin of ECL2 4 Å closer to the center of the EC pocket, a conformation that would clash with the N-terminus of CCL20 resolved in the active structure. Moreover, sandwiched in between TMs 5 and 6, the *para*-substituted phenyl groups of OXM1 and OXM2 wedge TM5 8 Å away from the center of the 7TM bundle, which makes room to accommodate the 4-5 Å inward movement of the entire TM6 associated with receptor inactivation. ECL3 provides an important docking site for the N-loop and the 30s-loop of CCL20 during receptor activation. The large movement of TM6 observed in the OXM analogue-bound structures results in an EC tip conformation of the helix that is now no longer compatible with CCL20 docking. Altogether, OXM binding stabilizes a distinct receptor conformation of the EC region from that utilized by CCL20, thereby allosterically inhibiting CCL20-mediated CCR6 activation.

Positive cooperativity between orthosteric agonists and G protein is a hallmark of GPCR activation. Cooperative stabilization of inactive CCR2 between orthosteric and allosteric antagonists has also been reported^17^. However, crosstalk between different allosteric ligands has not been well characterized to date. To understand the communication between the extra- and intracellular allosteric pockets discovered on CCR6, we examined binding of [^3^H]SQA1 to WT CCR6 with and without OXM1. To our surprise, the presence of OXM1 did not enhance the binding of [^3^H]SQA1 (Fig. S37), in contrast to the report that the presence of an orthosteric CCR2 antagonist increased binding of the intracellular antagonist, CCR2-RA^17^. We also studied the impact of SQA1 on the binding of OXM1 using SPR with a stabilized CCR6. Despite a slight increase in on-rate and a slight decrease in off-rate of OXM1 to CCR6 preincubated with SQA1, the overall change of the dissociation constant (K_D_) was within noise (Figs. S38). Taken together, these binding data suggest minimal communication occurred between the two allosteric pockets occupied by the OXM and SQA molecules, differing from the cooperativity observed between orthosteric agonists and intracellular signaling transducers.

## Discussion

Despite the significant advancement of cryo-EM and its tremendous success in active GPCR structure determination^35^, limited examples have been reported on the application of cryo-EM to small molecule-bound inactive GPCR structure determination. The challenges are twofold: the small size of the receptors alone, and the highly dynamic nature of this protein family. Here we present a successful example of the application of cryo-EM in tackling inactive GPCR structure determination for small molecule drug discovery. A BRIL fusion was introduced to replace the ICL3 with a carefully designed junction to achieve optimal rigidity. The BRIL/Fab/Nb module is proven to be an effective fiducial marker that allows robust particle alignment in cryo-EM^28^. Thermostabilizing mutations together with a ternary complex strategy of incorporating two different allosteric antagonists led to a highly stabilized CCR6. As a result, two high-resolution structures in complex with analogues from the SQA and OXM series were determined in this study, providing the first known examples of inactive CCR6 structures bound by small molecules. The structures reveal that SQA and OXM utilize different allosteric mechanisms to inhibit CCR6 activation. Importantly, the allosteric pockets of CCR6 discovered in our work are also likely present in other CKRs. Therefore, the lessons described herein are useful for structure-based drug discovery of small molecules targeting CCR6 and other related CKRs.

## Methods

No statistical methods were used to predetermine sample size. The experiments were not randomized. The investigators were not blinded to allocation during experiments and outcome assessment.

### CCR6 thermostabilization and characterization

HEK293T cells used for transient transfection for thermostabilization experiments were cultured in DMEM + GlutaMAX™-I with 4.5 g/L D-Glucose and Pyruvate (Gibco 31966-021), supplemented with 10 % (v/v) of fetal calf serum (FCS). Transient transfections were carried out using GeneJuice (Merck) in accordance with manufacturer’s protocols. Full-length WT CCR6 used as a template for thermostabilization was cloned into pcDNA3.1-based plasmid vector to allow transient expression with C-terminal GFP-TEV-myc-His×10 tags. Thermostability of transiently expressed CCR6 constructs in HEK293T cells was assayed by solubilized [^3^H]-SQA1 binding. Cells were solubilized in detergent (1 % (w/v) *n*-Dodecyl-*β*-D-Maltopyranoside (DDM, Anatrace), *n*-Decyl-*β*-D-Maltopyranoside (DM, Anatrace), or 2 % (w/v) *n*-Nonyl-*β*-D-Glucopyranoside (NG, Anatrace) with cholesteryl hemisuccinate (CHS, Sigma) added at 0.012 % (w/v) as appropriate) prior to incubation with radioligand and heating, and unbound radioligand was subsequently removed using microscale immobilized metal affinity chromatography (IMAC). Melting temperature (T*_m_*) was defined as the temperature at which 50 % of maximal radioligand binding was measured. Stabilizing mutants were made, identified, and combined as described previously^20,21^. The thermostabilized construct CCR6_Nα7.1 used in the cryo-EM study contains the following seven mutations: L47^1.33^A, L86^2.44^A, V96^2.54^A, S140^3.47^A, R159A, F236^5.60^A, G295^7.32^A (superscripts indicate Ballesteros– Weinstein numbering for GPCRs^22^). The thermostabilized construct CCR6_Nβ10.2 used in the crosstalk SPR study contains F73^1.59^A, V276^6.57^A, and A279^6.60^L in addition to the seven mutations of Nα7.1.

The affinity of [^3^H]-SQA1 binding to CCR6 constructs was determined by saturation binding. Increasing concentrations of [^3^H]-SQA1 were incubated for 2 h at 25 °C with aliquots of HEK293T cells transiently expressing CCR6 (or mock transfected control) resuspended in 50 mM HEPES pH 7.5, 150 mM NaCl, 5 mM MgCl_2_ and protease inhibitors. For the crosstalk study, the cells were prebound by 10 μM of OXM1 for 1 h before addition of [^3^H]-SQA1. The cell suspension with ligand mix was then chilled to 4 °C prior to receptor solubilization in 1 % DDM and 0.012 % CHS, and unbound radioligand was separated using microscale IMAC. Non-specific binding to mock-transfected controls was subtracted from binding to CCR6-containing samples prior to calculation of K_d_ in GraphPad Prism 9.0.0, using one site specific curve fitting.

### Expression, membrane preparation, and protein purification

The thermostabilized CCR6_Nα7.1-BRIL chimeric construct was cloned into a modified pFastBac1 vector (Invitrogen) with a haemagglutinin (HA) signal sequence followed by a cleavable FLAG-10×His tags. Protein expression was done in Spodoptera frugiperda (*Sf9*) insect cells (ATCC) using the Bac-to-Bac Baculovirus Expression System (Invitrogen). The insect cells were infected at a density of 2-3 × 10^6^ cells/ml with high titer viral stock multiplicity of infection (MOI) of 5.0. At 48 h post infection, cells were collected. Cell pellet was resuspended by dounce homogenization in buffer containing 10 mM HEPES pH 7.5, 150 mM NaCl, 10 mM MgCl_2_, 20 mM KCl, nuclease (Pierce) and EDTA-free cOmplete^TM^ protease inhibitor cocktail (Roche). Cell debris were removed by centrifugation at 1000×g for 20 min. The supernatant was then subjected to ultracentrifugation at 158,420×g for 30 min and the membrane pellet was resuspended by dounce homogenization in 50 mM HEPES pH 7.5, 150 mM NaCl, 10% glycerol, and EDTA-free cOmplete^TM^ protease inhibitor cocktail (Roche). The resuspension was solubilized at 4 °C for 4 h with 2% (w/v) Lauryl Maltose Neopentyl Glycol (LMNG, Anatrace) and 0.2% (w/v) CHS in the presence of 2 mg/ml iodoacetamide and 50 μM of the compounds. The sample was clarified by ultracentrifugation at 340,252×g for 30 min at 4 °C, and the supernatant was batch bound to TALON cobalt-affinity resin (Takara) overnight at 4 °C. The resin was washed with successively lower concentrations of detergent and eluted in buffer containing 50 mM HEPES pH 7.5, 500 mM NaCl, 0.003% LMNG, 0.0003% CHS, 300 mM imidazole, 50 μM OXM1 or OXM2, and 50 μM SQA1. The elution fractions were concentrated and incubated with anti-BRIL Fab and Fab-Nb in a molar ratio of 1 : 1.2 : 1.5 on ice for 1 hour. The complex was then subjected to size-exclusion chromatography (SEC) using a Superose 6 GL Increase 10/300 (Cytiva) in buffer containing 50 mM HEPES pH 7.5, 150 mM NaCl, 0.003% LMNG, 0.0003% CHS, 50 μM of OXM1 or OXM2 and 50 μM of SQA1. The SEC peak fraction was collected and concentrated to 1 mg/ml for cryo-EM grid preparation.

### Cryo-EM sample preparation and data acquisition

For both samples, cryo-EM grids were prepared with a Vitrobot Mark IV (Thermo Scientific) by applying 3 µl of sample to freshly glow-discharged Quantifoil Au R1.2/1.3 200-mesh grids and plunge freezing into liquid ethane. The grids were prepared using a blot force of +3, a blot time of 2 s, and an absorption time of 20 s. The sample chamber was kept at 4°C and 100% humidity.

The CCR6/SQA1/OXM1 dataset was collected using EPU (Thermo Scientific) on a Titan Krios operating at 300 KeV and equipped with a Selectris energy filter (Thermo Scientific) at a slit width of 10 eV and a Falcon 4 direct electron detector (Thermo Scientific) at a magnification of 215,000× for a magnified pixel size of 0.575 Å per pixel. Data were collected using a defocus range of −0.8 to −2.4 μm and a dose of 7.93 electrons per pixel per s for a total exposure time of 1.76 s divided into 60 subframes for a total accumulated dose of 40 electrons per Å^2^.

The CCR6/SQA1/OXM2 dataset was collected using EPU (Thermo Scientific) on a Titan Krios G3i operating at 300 KeV and equipped with a Bioquantum imaging filter at a slit width of 20 eV and a K3 direct electron detector (Gatan) at a magnification of 130,000× for a magnified pixel size of 0.654 Å per pixel. Data were collected using a defocus range of −0.4 to −1.8 μm and a dose of 18.27 electrons per pixel per s for a total exposure time of 1.06 divided into 45 subframes for a total accumulated dose of 45 electrons per Å^2^.

### Cryo-EM data processing

Data processing was carried out using cryoSPARC (Structura Biotechnology). For the CCR6/SQA1/OXM1 dataset, 12,852 movies were imported into cryoSPARC and subjected to patch motion correction and patch CTF estimation. Exposures with CTF fits better than 5 Å were selected (12,776 movies) and blob picking was used to select and extract 5,332,464 particles. These particles were subjected to multiple rounds of 2D classification. Ab initio model generation was carried out with 476,361 particles and 3 classes. The best model was used as a template for heterogeneous refinement starting with 2,168,721 particles using one good class model and 3 junk models. When most of the particles were sorted to the good class (487,232 particles), the particles were re-extracted using the full box size of 520 pix. These particles (482,484 particles) were subjected to nonuniform refinement, resulting in a map of 2.63 Å.

For the CCR6/SQA1/OXM2 dataset, 15,650 movies were subjected to patch motion correction and patch CTF estimation. Exposures with CTF fits better than 5 Å were selected (14,790 movies) and blob picking was used to select and extract 10,786,639 particles. These particles were subjected to multiple rounds of 2D classification. Ab initio model generation for three classes was carried out with 129,004 particles selected from 2D classification. The best model was used as a template for heterogeneous refinement starting with 1,127,335 particles using one good class model and 3 junk models. After three rounds of heterogeneous refinement, most of the particles sorted to the good class (371,301 particles), the particles were re-extracted using the full box size of 520 pix (368,745 particles). These particles were subjected to nonuniform refinement, resulting in a map of 3.02 Å.

### Model building and refinement

For each of the CCR6 structures presented, an atomic model predicted by AlphaFold2^37^ (thermostabilized CCR6_Nα7.1_ICL3_BRIL fusion construct sequence) was rigid-body fit into the map density. Starting models of the anti-BRIL Fab and the hinge-binding nanobody were obtained from PDB entry 6WW2. Following the docking of each model and ligands into the EM density map, iterative manual adjustment and real-space refinement was carried out using Coot^38^. Subsequently, global refinement and minimization in real space was performed using *phenix.real_space_refine* in Phenix^39^. The final models were visually inspected for fit to the map, and Molprobity^40^ was used for geometry validation. All structure figures were generated by using PyMOL^41^, UCSF Chimera^42^, and UCSF ChimeraX^43^.

### Intrinsic fluorescence-based thermal shift assay

Purified CCR6 ICL3-BRIL fusion protein (9 μM) was incubated with 10 μM of the specified compounds in buffer containing 50 mM HEPES pH 7.5, 500 mM NaCl, and 0.003% LMNG/0.0003% CHS. After a 5 min incubation at 25 °C, the samples were loaded into capillary tubes for Tycho NT.6 instrument (NanoTemper). Tryptophan fluorescence at emission wavelengths of 350 nm and 330 nm were measured as the sample temperature was ramped from 35 °C to 95 °C according to the preprogrammed instrument protocol. The ratio of the emission fluorescence was used to determine the T_i_ for the melting curves reported by the instrument software.

### CPM-based thermal shift assay

CPM-based thermal shift assay was conducted for CCR6 WT by a differential scanning fluorimetry assay adapted from a previous publication^44^ using UNit (Unchained Labs). In brief, 5 μM of protein was incubated for 5 min at 25 °C with the presence of 10 μM of testing compound or 1% DMSO as control in buffer containing 50 mM HEPES pH 7.5, 500 mM NaCl, 0.05% DDM, and 2 μM of the thiol-specific fluorochrome *N*-[4-(7-diethylamino-4-methyl-3-coumarinyl)phenyl]-maleimide (CPM) dye. After incubation, the samples were heated from 20 °C to 80 °C at a temperature ramping rate of 2 °C/min, during which the CPM fluorescence (excitation 384 nm, emission 470 nm) was recorded. Melting temperature (T_m_) is defined as the temperature at which 50% of the protein was denatured based on Boltzmann sigmoidal fitting.

### OXM discovery by SQA-based virtual screen

A field-based 3D similarity search as implemented in the Blaze module within the Cresset software^45–47^ was performed using the small molecule x-ray conformation^48^, of an SQA analogue as a query. Briefly the method uses electrostatics and shape of the query ligand to rapidly search large chemical collections for molecules with similar properties. The basis of electrostatic potential calculations is the polarizable XED force field which allows an accurate description of the charges around atoms distribution. Each molecule is described as a Field Point using the local extrema of four molecular fields: positive electrostatic, negative electrostatic, surface (van der Waals interactions), and hydrophobic. Unlike traditional electrostatic field methods that place single point partial charges at atom centers, this field redefines charges towards a multipole electron distribution around the atom. The molecular fields developed from this representation are condensed to their local field points and used to rapidly search and score scaffolds with similar electronic properties for molecules in our internal database. The OXM analogue OXM1 with a similarity score of 0.74 was identified from this approach.

### NMR experiments using ROE techniques

Rotating-frame nuclear Overhauser enhancements (ROEs) or nuclear Overhauser effect (NOE) experiments were collected using previously reported methods^29^ in C*D*Cl_3_ or DMSO-*d*_6_. The absolute values for the ROE/NOE intensities were measured relative to the irradiated peak which was arbitrarily assigned a value of −100. Calibration for this experiment used a ROE/NOE intensity for the adjacent aromatic proton at a distance of 2.8 Å.

### RDC NMR solution conformations

The proton and carbon assignments for OXM1, OXM2, OXM3, and OXM4 were completed using a combination of 1D proton, 2D COSY, 2D HSQC, 2D HMBC and 1D carbon experiments using standard Bruker pulse sequences and a concentration of 7 mg/200 μl. The assignments are shown in Supplementary Information (Figs. S1-S5, S8-12, S15-19, S22-26). All NMR experiments were collected on a Bruker AVANCE III spectrometer operating at a ^1^H-Larmor frequency of 600.1 MHz that was equipped with a 5 mm TCI helium cryoprobe using Topspin 3.2. The chemical shifts are referenced to residual solvent (deuterated DMSO). Both ^1^*J*_CH_ and ^1^*J*_NH_’s^49^ were collected for the four OXM analogues using compressed DMSO-d_6_ swollen poly-HEMA gels in New Era Enterprises compression devices. For the proton-carbon and proton-nitrogen *J*-couplings, two *J*-scaled Bird (JSB) HC-HSQC spectra were collected for each compound, see Supplementary Information (Figs. S6-S7, S13-14, S20-21, S27-28). The first JSB HC-HSQC was collected on the compound in DMSO-d_6_ alone (isotropic media) while the second JSB HC-HSQC was collected on the compound in a poly-HEMA gel (anisotropic media). The residual dipolar coupling (RDC) values (^1^*D*) for proton-carbon and proton-nitrogen J-couplings were determined using the formula: ^1^*D* = ^1^*T* – ^1^*J* as reported previously^30^. Overall, RDCs (^1^*J*_CH_ and ^1^*J*_NH_) in a range of −8.9 to +13.3 Hz were measured for each compound in DMSO-d_6_ as shown in Supplementary Information (Table S3). The RDC data was fit to each member of a conformational ensemble that consisted of 80-160 low energy structures that were generated in a solvent dependent manner based on molecular dynamic simulations. Two files were generated for each compound: a linear conformer file and a U-shaped conformer file. The RDC restraints were fit to each file independently and the best results (lowest Q) are reported. The RDC data was fit using Mestre MSpin/Stereofitter software using the input files provided (see Supplementary Information)^50^. With this software, the Cornilescu *Q*-factor (*Q*)^51^ was used to judge the goodness of the fit for each conformation. The conformations with the best fit had *Q* = 0.005 for OXM1, *Q* = 0.097 for OXM2, *Q* = 0.001 for OXM3, and *Q* = 0.035 for OXM4. In addition, the experimental RDC’s correlated well with the back-calculated values as shown in Table S3.

### Conformer generation for RDC fitting experiments

All structures were first subject to geometry optimization in Macromodel^52^ using the OPLS3e forcefield and water as solvent, with default parameters and convergence criteria. A conformational search was performed on each structure using the mixed MCMM/low-mode search as implemented in the Macromodel program with 1000 steps per rotatable bond and an energy window of 21 kJ mol−1 for retention of conformers. The conformer files (OXM1, OXM2, OXM3, and OXM4) are provided in a supplementary file for each compound.

### OXM binding measurement by SPR

Binding affinity and kinetics of OXM1 were measured with the CCR6_Nα7.1 construct using a Surface Plasmon Resonance (SPR)-based binding assay on Biacore T200 instrument (Cytiva). The steps of sensor preparation and protein capture were carried out at 25 °C and the binding interactions were measured at 10 °C. The CCR6 protein was initially captured via immobilized metal affinity chromatography (IMAC) followed by amine coupling onto an activated Ni-NTA sensor chip treated by a 1:1 mixture of 0.05 M *n*-hydroxysuccinimide (NHS) and 0.2 M 1-ethyl-3-(3-dimethylaminopropyl) carbodiimide (EDC). The experiments were carried out in 50 mM HEPES buffer (pH 7.5) containing 150 mM NaCl, 0.1% DDM, and 3% DMSO. OXM1 was injected in a concentration series consisting of 6-point, 3-fold dilutions from 10 μM to 0.014 μM. The data were processed and analyzed using Biacore Biaeval and Scrubber 2.0 software. The binding affinities and on and off rates were calculated by globally fitting the sensorgrams to a 1:1 binding model.

### Crosstalk study by SPR

The crosstalk SPR experiments were performed using a Biacore T200 Instrument (Cytiva) equipped with a NiHC 200M sensor chip (Xantec) at 25 °C. The run buffer for all experiments, unless otherwise stated, was 50 mM HEPES pH 7.5, 500 mM NaCl, 1 mM MgCl_2_, 0.03% DDM, 0.003% CHS, 5% DMSO. The CCR6_Nβ10.2 protein was immobilized on the sensor chip by Ni-NTA pull down and amine coupling. 0.5 M EDTA pH 8.0 was injected for 60 s at 5 µL/min followed by a 60 s injection of run buffer containing 0.5 mM NiCl_2_. The surface was activated with a 1:1 mixture of 0.1 M NHS and 0.5 M EDC at a flow rate of 10 µL/min. The receptors were injected at a concentration of 0.5 µM at 5 µL/min until a surface response of 2000 RU was achieved. The surface was equilibrated for 10 h in run buffer before commencing experiments. OXM1 binding affinity was measured in a 5-point, 2-fold dilution series in single cycle kinetics format with a top concentration of 1 µM in run buffer with and without 100 nM SQA1. The flow rate of binding measurement was 50 µL/min. The association phase was 120 s and dissociation phase was between 1800 s. All experiments were run in triplicate. The data were processed using Biacore T200 Evaluation Software (Version 2.0, Cytiva). Responses were corrected for volume exclusion effects. Sensorgrams were reference and blank subtracted before fitting with 1:1 binding model which accounted for drift, bulk shift and mass transport. Sensorgram images were prepared using GraphPad Prism software.

### Protein surface expression measurement

HEK293 cells transiently expressing CCR6 constructs or mock transfected control after 72 h post transfection were detached using an enzyme-free dissociation buffer. Cells were blocked with 0.1 mg/ml mouse IgG prior to staining with BV-421 labeled anti-human CCR6 antibody (Clone 11A9, BD Biosciences) for 1 h at 4 °C. After washing, cells were analyzed on a BD LSRFortessa cytometer (BD Biosciences) for cell surface expression of CCR6 by determining the mean fluorescence intensity (MFI) of BV-421 using FlowJo™ Software (Becton, Dickinson and Company).

### Radioligand saturation binding in HEK293 or CHO-K1 cells

Saturation binding experiments were performed on HEK293 cells transiently expressing human CCR6 48 h post transfection, or CHO-K1 cells stably expressing human CCR6 (DiscoverX). HEK293 cells were plated at 80,000/well in growth media (DMEM + 10% Serum + 20 mM HEPES) in white/clear bottom PDL coated 96 well plates (Corning); CHO-K1 cells were plated at 50,000/well in white/clear bottom 96 well Isoplates (PerkinElmer) in growth media a day before the experiments. On the day of assay, cells in the assay well were treated with either 0.3-0.4 % DMSO (for total binding) or 90-120 μM of unlabeled SQA1 (for non-specific binding), followed by addition of a serial dilution of [^3^H]-SQA1 in growth media. For each dose of [^3^H]-SQA1, triplicate reactions were prepared for both total and non-specific binding conditions. After 3 hours of incubation at room temperature, cells were washed 3 times with wash buffer (PBS + 1 mM CaCl_2_ + 0.1% BSA) before adding Microscint-20 (PerkinElmer) for tritium concentration measurement using a TriLux MicroBeta2 plate reader (PerkinElmer). Data was analyzed with GraphPad Prism software using a one-site saturation binding equation.

### CCR6+ T cell chemotaxis assay

Human CD4^+^CCR6^+^CXCR3^-^ T cells were isolated from human donor leukopaks using EasySep™ Human Th17 Cell Enrichment Kit (StemCell Technologies, 18162). To obtain large quantities of cells, CCR6+ T cells were activated with Dynabeads Human T-activator (Gibco, 11132D) at a density of 1 × 10^6^ cell/ml in growth media (RPMI1640 media with 10% serum, 4 ng/ml IL-2) with a 1:1.5 cell to bead ratio. On day 4 post activation, Dynabeads were removed from the culture. The activated T cells were maintained at 1‒2 × 10^6^ cells/ml for 15 days by feeding fresh growth media when needed. The CCR6+ T cell chemotaxis assay was carried out on day 12 to 15 post T cell activation using the 96 well ChemoTx^®^ disposable chemotaxis system (Neuroprobe 101-5) according to the manufacturer’s protocol. After one wash with assay buffer (1 × HBSS containing 20 mM HEPES and 0.25% BSA), cells were incubated with test compounds for 30 minutes at room temperature prior to initiation of chemotaxis. CCL20 induced CCR6^+^ human T cell chemotaxis with an EC_50_ = 0.19 nM (pEC50 = 9.7 ± 0.9, mean ± s.d., *n* = 5). CCL20 was used at 0.5 nM in all measurements. For IC50 determination, the top and bottom of the chemotaxis chamber contained the same concentration of compound. DMSO was kept constant at 0.1% (v/v) in all wells. The final concentration of CCL20 (Peprotech, 300-29A) in the bottom chamber is 0.5 nM. The fully assembled chemotaxis plate was placed in a cell culture incubator at 37 °C, 5% CO_2_ for 1 h. After incubation, the top filter was removed, followed by a quick freeze of the bottom chamber at −80 °C for 1 h. The migrated cells in the bottom chamber were stained with CyQUANT dye (Life Technologies, C7026) for cell number determination. Data was analyzed with GraphPad Prism Software using a non-linear regression analysis of dose response curves for IC50 determination.

### GPCR β-arrestin assay panel

OXM1 was tested at 1 μM for agonistic and antagonistic activity on a panel of GPCRs in *gpcr*MAX℠ GPCR assay panel using PathHunter^®^ β-arrestin enzyme complementation technology as described in Eurofins Discovery website (gpcrMAX-(discoverx.com)).

**Preparation of [^3^H]-SQA1 and Synthesis of different OXM analogues, OXM1, OXM2, OXM3, and OXM4** are in supplementary information.

### Data availability statement

The cryo-EM density maps for the complexes are deposited in the Electron Microscopy Data Bank (EMDB) under accession codes EMD-xxxxx (CCR6/SQA1/OXM1) and EMD-xxxxx (CCR6/SQA1/OXM2). The coordinates for the models are deposited in the Worldwide Protein Data Bank (wwPDB) under accession codes XXXX (CCR6/SQA1/OXM1) and XXXX (CCR6/SQA1/OXM2). Solution conformer library files of the OXM analogues are provided as source data in .sdf format. Other source data are also provided with this paper. All the other data supporting the findings of this study are available within the article and its Supplementary Information files and from the corresponding author upon reasonable request.

## Acknowledgements

Cryo-EM data for the CCR6/SQA1/OXM2 complex was collected at the Cryo-EM Facility at MIT.nano. We thank K. Fennell and H. Zhao for recombinant protein expression support; K. Schildknegt, T. Kenakin, and E.J. Corey for consultancy support for the CCR6 antagonist project; C. Tang, Q. Jin, and N. Fadeyi for synthesis support; X. Qiu and D. Hepworth, for project management support. We thank all members of Pfizer CCR6 research project team for suggestions and comments.

## Author contributions

D.J.W. performed protein purification, thermal shift experiments, prepared cryo-EM grids, collected and processed cryo-EM data, determined the structures, and assisted with preparing the manuscript; B.S.G. supervised discovery and design of the OXM analogues; K.A.F. performed NMR studies; W.L. designed and supervised all pharmacology experiments; J.A. performed radioligand binding and CCR6 mutagenesis studies; M.E.S. supervised discovery of the SQA analogue; R.J.U. performed ligand-based virtual screening and generated conformer libraries for the OXM analogues; J.V. performed protein purification; K.K.C. performed T cell chemotaxis experiments; R.D. performed protein surface expression measurements; P.V.S performed SPR experiments; F.V. and R.K.F. performed T cell chemotaxis experiments; A.D., A.F., C.C., G.C., J.J.M., J.I.T., P.N., P.M., and V.L. performed synthesis; D.L., E.S., and C.O. performed StaR engineering, characterization, and saturation bindings; B.J.H. performed crosstalk SPR studies; G.S.M. purified CCR6 protein for crosstalk SPR studies. I.W.W. collected cryo-EM data of the SQA1/OXM2 complex; S.H. assisted with cryo-EM data collection of the SQA1/OXM1 complex; M.R. and R.M.C. supervised StaR engineering and characterization; M.F.C. managed structural and biophysical studies and reviewed the data; G.B. analyzed and interpreted pharmacology data; A.T. supervised and managed the project; H.W. designed and supervised all structural and biophysical studies, determined the structures, analyzed the results and wrote the manuscript.

## Additional information

**Extended data** is available for this paper at (web).

**Supplemental information** is available for this paper at (web).

**Reprints and permissions information** is available at (web).

## References

1 Hughes, C. E. & Nibbs, R. J. B. A guide to chemokines and their receptors. FEBS J 285, 2944–2971 (2018). 10.1111/febs.14466

2 Bonecchi, R. & Graham, G. J. Atypical Chemokine Receptors and Their Roles in the Resolution of the Inflammatory Response. Front Immunol 7, 224 (2016). 10.3389/fimmu.2016.00224

3 Murdoch, C. & Finn, A. Chemokine receptors and their role in inflammation and infectious diseases. Blood 95, 3032–3043 (2000).

4 Chow, M. T. & Luster, A. D. Chemokines in cancer. Cancer Immunol Res 2, 1125–1131 (2014). 10.1158/2326-6066.CIR-14-0160

5 Murphy, P. M. et al. International union of pharmacology. XXII. Nomenclature for chemokine receptors. Pharmacol Rev 52, 145–176 (2000).

6 Paulissen, S. M., van Hamburg, J. P., Dankers, W. & Lubberts, E. The role and modulation of CCR6+ Th17 cell populations in rheumatoid arthritis. Cytokine 74, 43–53 (2015). 10.1016/j.cyto.2015.02.002

7 Schutyser, E., Struyf, S. & Van Damme, J. The CC chemokine CCL20 and its receptor CCR6. Cytokine Growth Factor Rev 14, 409–426 (2003).

8 Homey, B. et al. Up-regulation of macrophage inflammatory protein-3 alpha/CCL20 and CC chemokine receptor 6 in psoriasis. J Immunol 164, 6621–6632 (2000). 10.4049/jimmunol.164.12.6621

9 Skovdahl, H. K. et al. Expression of CCL20 and Its Corresponding Receptor CCR6 Is Enhanced in Active Inflammatory Bowel Disease, and TLR3 Mediates CCL20 Expression in Colonic Epithelial Cells. PLoS One 10, e0141710 (2015). 10.1371/journal.pone.0141710

10 Ranasinghe, R. & Eri, R. Modulation of the CCR6-CCL20 Axis: A Potential Therapeutic Target in Inflammation and Cancer. Medicina (Kaunas) 54 (2018). 10.3390/medicina54050088

11 Katchar, K., Kelly, C. P., Keates, S., O’Brien M, J. & Keates, A. C. MIP-3alpha neutralizing monoclonal antibody protects against TNBS-induced colonic injury and inflammation in mice. Am J Physiol Gastrointest Liver Physiol 292, G1263–1271 (2007). 10.1152/ajpgi.00409.2006

12 Hedrick, M. N. et al. CCR6 is required for IL-23-induced psoriasis-like inflammation in mice. J Clin Invest 119, 2317–2329 (2009). 10.1172/jci37378

13 Wasilko, D. J. et al. Structural basis for chemokine receptor CCR6 activation by the endogenous protein ligand CCL20. Nat Commun 11, 3031 (2020). 10.1038/s41467-020-16820-6

14 Rennard, S. I. et al. CXCR2 Antagonist MK-7123. A Phase 2 Proof-of-Concept Trial for Chronic Obstructive Pulmonary Disease. Am J Respir Crit Care Med 191, 1001–1011 (2015). 10.1164/rccm.201405-0992OC

15 Gonsiorek, W. et al. Pharmacological characterization of Sch527123, a potent allosteric CXCR1/CXCR2 antagonist. J Pharmacol Exp Ther 322, 477–485 (2007). 10.1124/jpet.106.118927

16 Oswald, C. et al. Intracellular allosteric antagonism of the CCR9 receptor. Nature 540, 462–465 (2016). 10.1038/nature20606

17 Zheng, Y. et al. Structure of CC chemokine receptor 2 with orthosteric and allosteric antagonists. Nature 540, 458–461 (2016). 10.1038/nature20605

18 Jaeger, K. et al. Structural Basis for Allosteric Ligand Recognition in the Human CC Chemokine Receptor 7. Cell 178, 1222–1230 e1210 (2019). 10.1016/j.cell.2019.07.028

19 Li, W. et al. A Novel C-C Chemoattractant Cytokine (Chemokine) Receptor 6 (CCR6) Antagonist (PF-07054894) Distinguishes between Homologous Chemokine Receptors, Increases Basal Circulating CCR6(+) T Cells, and Ameliorates Interleukin-23-Induced Skin Inflammation. J Pharmacol Exp Ther 386, 80–92 (2023). 10.1124/jpet.122.001452

20 Magnani, F. et al. A mutagenesis and screening strategy to generate optimally thermostabilized membrane proteins for structural studies. Nat Protoc 11, 1554–1571 (2016). 10.1038/nprot.2016.088

21 Robertson, N. et al. The properties of thermostabilised G protein-coupled receptors (StaRs) and their use in drug discovery. Neuropharmacology 60, 36–44 (2011). 10.1016/j.neuropharm.2010.07.001

22 Ballesteros, J. A. & Weinstein, H. Integrated methods for the construction of three-dimensional models and computational probing of structure-function relations in G protein-coupled receptors. Methods in Neurosciences 25, 63 (1995). 10.1016/S1043-9471(05)80049-7

23 Roth, C. B., Hanson, M. A. & Stevens, R. C. Stabilization of the human beta2-adrenergic receptor TM4-TM3-TM5 helix interface by mutagenesis of Glu122(3.41), a critical residue in GPCR structure. J Mol Biol 376, 1305–1319 (2008). 10.1016/j.jmb.2007.12.028

24 Chun, E. et al. Fusion partner toolchest for the stabilization and crystallization of G protein-coupled receptors. Structure 20, 967–976 (2012). 10.1016/j.str.2012.04.010

25 Liu, W. et al. Structural basis for allosteric regulation of GPCRs by sodium ions. Science 337, 232–236 (2012). 10.1126/science.1219218

26 Mukherjee, S. et al. Synthetic antibodies against BRIL as universal fiducial marks for single-particle cryoEM structure determination of membrane proteins. Nat Commun 11, 1598 (2020). 10.1038/s41467-020-15363-0

27 Ereno-Orbea, J. et al. Structural Basis of Enhanced Crystallizability Induced by a Molecular Chaperone for Antibody Antigen-Binding Fragments. J Mol Biol 430, 322–336 (2018). 10.1016/j.jmb.2017.12.010

28 Tsutsumi, N. et al. Structure of human Frizzled5 by fiducial-assisted cryo-EM supports a heterodimeric mechanism of canonical Wnt signaling. Elife 9 (2020). 10.7554/eLife.58464

29 Butts, C. P. et al. Interproton distance determinations by NOE--surprising accuracy and precision in a rigid organic molecule. Org Biomol Chem 9, 177–184 (2011). 10.1039/c0ob00479k

30 Farley, K. A. et al. Cyclic Peptide Design Guided by Residual Dipolar Couplings, J-Couplings, and Intramolecular Hydrogen Bond Analysis. J Org Chem 84, 4803–4813 (2019). 10.1021/acs.joc.8b02811

31 Horst, R., Farley, K. A., Kormos, B. L. & Withka, J. M. NMR spectroscopy: the swiss army knife of drug discovery. J Biomol NMR 74, 509–519 (2020). 10.1007/s10858-020-00330-0

32 Liu, K. et al. Structural basis of CXC chemokine receptor 2 activation and signalling. Nature (2020). 10.1038/s41586-020-2492-5

33 Wu, H. et al. Structure of the human kappa-opioid receptor in complex with JDTic. Nature 485, 327–332 (2012). 10.1038/nature10939

34 Salchow, K. et al. A common intracellular allosteric binding site for antagonists of the CXCR2 receptor. Br J Pharmacol 159, 1429–1439 (2010). 10.1111/j.1476-5381.2009.00623.x

35 Danev, R. et al. Routine sub-2.5 A cryo-EM structure determination of GPCRs. Nat Commun 12, 4333 (2021). 10.1038/s41467-021-24650-3

36 Molecular Operating Environment (MOE), 2022.02 Chemical Computing Group ULC, 910-1010 Sherbrooke St. W., Montreal, QC H3A 2R7, Canada. (2023).

37 Jumper, J. & Hassabis, D. Protein structure predictions to atomic accuracy with AlphaFold. Nat Methods 19, 11–12 (2022). 10.1038/s41592-021-01362-6

38 Emsley, P., Lohkamp, B., Scott, W. G. & Cowtan, K. Features and development of Coot. Acta Crystallogr D Biol Crystallogr 66, 486–501 (2010). 10.1107/S0907444910007493

39 Liebschner, D. et al. Macromolecular structure determination using X-rays, neutrons and electrons: recent developments in Phenix. Acta Crystallogr D Struct Biol 75, 861–877 (2019). 10.1107/S2059798319011471

40 Chen, V. B. et al. MolProbity: all-atom structure validation for macromolecular crystallography. Acta Crystallogr D Biol Crystallogr 66, 12–21 (2010). 10.1107/S0907444909042073

41 Schrödinger, L. & DeLano, W. PyMoL, <http://www.pymol.org/pymol> (2020).

42 Pettersen, E. F. et al. UCSF Chimera--a visualization system for exploratory research and analysis. J Comput Chem 25, 1605–1612 (2004). 10.1002/jcc.20084

43 Pettersen, E. F. et al. UCSF ChimeraX: Structure visualization for researchers, educators, and developers. Protein Sci 30, 70–82 (2021). 10.1002/pro.3943

44 Alexandrov, A. I., Mileni, M., Chien, E. Y., Hanson, M. A. & Stevens, R. C. Microscale fluorescent thermal stability assay for membrane proteins. Structure 16, 351–359 (2008). 10.1016/j.str.2008.02.004

45 Blaze, version, Cresset, Litlington, Cambridgeshire, UK;, <https://www.cresset-group.com/software/blaze/> (

46 Cheeseright, T. J., Mackey, M. D., Melville, J. L. & Vinter, J. G. FieldScreen: virtual screening using molecular fields. Application to the DUD data set. J Chem Inf Model 48, 2108–2117 (2008). 10.1021/ci800110p

47 Cheeseright, T., Mackey, M., Rose, S. & Vinter, A. Molecular field extrema as descriptors of biological activity: definition and validation. J Chem Inf Model 46, 665–676 (2006). 10.1021/ci050357s

48 Groom, C. R., Bruno, I. J., Lightfoot, M. P. & Ward, S. C. The Cambridge Structural Database. Acta Crystallogr B Struct Sci Cryst Eng Mater 72, 171–179 (2016). 10.1107/S2052520616003954

49 Gil-Silva, L. F., Santamaria-Fernandez, R., Navarro-Vazquez, A. & Gil, R. R. Collection of NMR Scalar and Residual Dipolar Couplings Using a Single Experiment. Chemistry 22, 472–476 (2016). 10.1002/chem.201503449

50 Navarro-Vazquez, A. MSpin-RDC. A program for the use of residual dipolar couplings for structure elucidation of small molecules. Magn Reson Chem 50 **Suppl 1**, S73–79 (2012). 10.1002/mrc.3905

51 Cornilescu, G., Delaglio, F. & Bax, A. Protein backbone angle restraints from searching a database for chemical shift and sequence homology. J Biomol NMR 13, 289–302 (1999). 10.1023/a:1008392405740

52 Macromodel, Schrodinger, LLC, New York, <http://www.schrodinger.com> (2011).

