## Supplementary Information for "Structural basis for CCR6 modulation by allosteric antagonists"

### Preparation of [$^3$H]-SQA1 and Synthesis of different OXM analogues, OXM1, OXM2, OXM3, and OXM4

#### Preparation of [$^3$H]-SQA1.

**General synthetic conditions:** Reagents and solvents were obtained from Aldrich and/or Alfa and were used without further purification. Solvents were commercial anhydrous grades and were used as received. All reactions were conducted with continuous magnetic stirring under an atmosphere of dry nitrogen unless otherwise specified. Chromatographic purifications were affected by medium pressure ("flash") chromatography on silica gel unless noted otherwise. Compounds were characterized by proton (1H) NMR spectra using Bruker spectrometers and are reported in parts per million (ppm) relative to the residual resonances of the deuterated solvent. All $^{13}$C NMR spectra were proton decoupled.

**(*S*)-*N*-((4-bromofuran-2-yl)methylene)-2-methylpropane-2-sulfinamide:** To a solution of 4-bromo-2-furaldehyde (1.0 g, 5.7 mmol) in THF (29 ml) was added $Ti(OEt)_4$ (2.78 ml, 11.4 mmol) at ambient temperature. The mixture was stirred for 20 min and (*S*)-2-methyl-2-propanesulfinamide (727 mg, 6.0 mmol) was added and the resulting mixture was stirred for 16 h. The reaction was quenched with water (10 ml). The mixture was filtered, and the filter cake was washed with ethyl acetate (20 ml × 2). The combined filtrate was evaporated under reduced pressure to afford crude residue that was purified by column chromatography (90:10 heptanes:ethyl acetate to 100% ethyl acetate) to afford the title compound (1.2 g, 76%) as a white solid. $^1$H NMR (400 MHz, $CDCl_3$) δ 1.28 (s, 9H), 7.06 (s, 1H), 7.65 (s, 1H), 8.35 (s, 1H); $^{13}$C NMR (101 MHz, $CDCl_3$) δ 22.6, 58.2, 102.2, 120.1, 144.7, 149.4, 151.2; LCMS (ESI): >99% (UV), $[M+H]^+$ m/z 278.1

**(*S*)-*N*-((*R*)-1-(4-bromofuran-2-yl)allyl)-2-methylpropane-2-sulfinamide:** To a solution of (*R*)-*N*-((4-bromofuran-2-yl)methylene)-2-methylpropane-2-sulfinamide (1.0 g, 3.6 mmol) in dichloromethane (20 ml) was added dropwise a solution of vinyl magnesium bromide (11.5 ml, 1.0 M) at -78 °C. The resulting solution was stirred at -78 °C for 1 h. The reaction was quenched by sat. aq. $NH_4Cl$ (10 ml) at 0 °C. The resulting mixture was concentrated to afford crude residue which was purified by flash chromatography (90:10 heptanes:Ethyl acetate to 100% ethyl acetate) to afford the title compound (200 mg, 20%) as the minor diastereomeric product. $^1$H NMR (400 MHz, $CDCl_3$) δ 1.22 (s, 9H), 3.44 (d, *J* = 5.5 Hz, 1H), 4.97-5.00 (m, 1H), 5.29-5.41 (m, 2H), 6.05 (ddd, *J* = 17.1, 10.2, 6.6 Hz, 1H), 6.33 (s, 1H), 7.40 (s, 1H); $^{13}$C NMR (101 MHz, $CDCl_3$) δ 22.5, 56.0, 56.2, 100.1, 111.2, 118.6, 135.5, 140.8, 154.3; MS (ESI): $[M+H]^+$ m/z 307.0.

**(*S*)-2-methyl-N-((*R*)-1-(4-(prop-1-en-2-yl)furan-2-yl)allyl)propane-2-sulfinamide:** A mixture of (*S*)-N-((*R*)-1-(4-bromofuran-2-yl)allyl)-2-methylpropane-2-sulfinamide (200 mg, 0.65 mmol), potassium carbonate (271 mg, 1.63 mmol), isopropenylboronic acid pinacol ester (274 mg, 1.63 mol) and Pd(dppf)$Cl_2$ (38 mg, 0.05 mmol) in dioxane (17.8 ml) and water (4.0 ml) was degassed by bubbling $N_2$ gas through the solution for 10 min. The reaction was then refluxed (100 °C) for 5 h. After cooling to ambient temperature, water (10 ml) was added and the mixture was extracted ethyl acetate (20 ml × 2). The organic layers were combined and evaporated under reduced pressure and then purified by column chromatography (90:10 heptanes:ethyl acetate to 100% ethyl acetate) to afford the title compound as an oil (70 mg, 40%). $^1$H NMR (400 MHz, $CDCl_3$) δ 1.19 (s, 9H), 2.00 (s, 3H), 3.46 (br s, 1H), 4.85-4.96 (m, 1H), 5.00 (br s, 1H), 5.19 (s, 1H), 5.27-5.35 (m, 1H), 5.40 (d, $J$ = 17.2 Hz, 1H), 6.10 (ddd, $J$ = 16.88, 10.24, 6.83 Hz, 1H), 6.44 (s, 1H), 7.38 (s, 1H); $^{13}$C NMR (101 MHz, $CDCl_3$) δ 20.9, 22.6, 55.6, 56.0, 105.5, 111.1, 118.7, 128.3, 134.7, 135.3, 138.9, 154.2.

**(*R*)-1-(4-(propan-2-yl-1,2-$t_2$)furan-2-yl)propan-2,3-$t_2$-1-amine hydrochloride salt :** To a solution of (*S*)-2-methyl-N-((*R*)-1-(4-(prop-1-en-2-yl)furan-2-yl)allyl)propane-2-sulfinamide (5 mg, 0.019 mmol) in THF (1 ml) in a tritiation flask was added $Pd(OH)_2$/C (10 mg). The reaction was stirred under an atmosphere of tritium gas (5 Ci) for 3 h. The reaction was filtered, and the filtrate was evaporated under reduced pressure. The residue was repeatedly dissolved in ethanol and evaporated under reduced pressure to remove residual reagents. The resulting residue was dissolved in ethyl acetate (0.3 ml) and 12 M HCl (3 μl) was added. The resulting solution was stirred for 40 min. The reaction then concentrated under reduced pressure. The residue was repeatedly dissolved in ethanol and evaporated under reduced pressure to remove residual reagents to provide the title compound that was used in the next step without further purification.

**4-((3,4-dioxo-2-(((1*R*)-1-(4-(propan-2-yl-1,2-$t_2$)furan-2-yl)propyl-2,3-$t_2$)amino)cyclobut-1-en-1-yl)amino)-3-hydroxy-*N*,*N*-dimethylpicolinamide ([$^3$H]-SQA1):** To a solution of (*R*)-1-(4-(propan-2-yl-1,2-$t_2$)furan-2-yl)propan-2,3-$t_2$-1-amine hydrochloride salt (assumed 0.019 mmol) in ethanol (0.3 ml) was added diisopropylethylamine (30 μl) followed by 4-((2-ethoxy-3,4-dioxocyclobut-1-en-1-yl)amino)-3-hydroxy-N,N-dimethylpicolinamide (4.5 mg, 0.015 mmol). The reaction was stirred for 5 h at ambient temperature. The reaction was evaporated under reduced pressure and then initially purified by column chromatography (90:10 dichloromethane:methanol). The resulting material was further purified by HPLC (Gemini C18 25 × 1 cm with 0.1% trifluoracetic acid in water (v/v) as the mobile phase A; 0.1% trifluoracetic acid in acetonitrile (v/v) as the mobile phase B; 20% B to 100% B over 100 min at 3 ml/min flow rate) to provide the title compound which was dissolved in ethanol. The radiochemical purity was determined by high performance liquid chromatography and has a specific activity of 106Ci/mmol. The title compound was dispensed and stored as 8 × 2mCi packs as an ethanol solution at 10 mCi/ml. HPLC (Inertsil ODS3 5 μm 250 × 2.6 mm at 22 °C with 0.1% trifluoracetic acid in water (v/v) as the mobile phase A; 0.1% trifluoracetic acid in acetonitrile (v/v) as the mobile phase B; 20% B to 80% B over 20 min at 1 ml/min flow rate) RT 13.483 min; 97.5%; HRMS (ESI): $[M+H]^+$ m/z 435.2219.

**Preparation of OXM1**

**(*R*)-*N*-(3-chloro-2-hydroxypropyl)-1-(4-chlorophenyl)cyclopropane-1-carboxamide:** To a solution of 1-(4-chlorophenyl)-1-cyclopropanecarboxylic acid (2.00 g, 10.17 mmol) in DMF (68.5 ml) was added (*R*)-1-amino-3-chloropropan-2-ol (1.00 g, 6.85 mmol), HATU (3.91 g, 10.3 mol), and *N*-methyl morpholine (2.64 ml, 24.0 mmol). The reaction was stirred at ambient temperature overnight. The reaction was quenched with saturated aqueous ammonium chloride solution and extracted with ethyl acetate. The organic layers were combined and washed with water (100 ml × 2) and saturated aqueous NaCl solution then was evaporated under reduced pressure. The crude residue was purified by column chromatography (90:10 heptanes:ethyl acetate to 100% ethyl acetate) to afford the title compound (1.61 g, 82%). $^1$H NMR (400 MHz, $CDCl_3$) δ 1.05-1.11 (m, 2H), 1.62-1.65 (m, 2H), 3.27-3.34 (m, 1H), 3.81-3.87 (m, 2H), 5.69 (br s, 1H), 7.34-7.39 (m, 4H); MS (ESI): $[M+H]^+$ m/z 288.2.

**(*R*)-1-(4-chlorophenyl)-N-(oxiran-2-ylmethyl)cyclopropane-1-carboxamide:** A solution of (*R*)-N-(3-chloro-2-hydroxypropyl)-1-(4-chlorophenyl)cyclopropane-1-carboxamide (135 mg, 0.47 mmol) in THF (0.47 ml) in a screw cap vial was cooled to -40 °C. To the reaction vial was added potassium-*t*-butoxide (33.8 mg, 0.47 mmol) and the reaction was stirred for 10 min. The reaction was filtered through silica pad which was washed with 5% MeOH in DCM. The filtrate was concentrated under reduced pressure. The crude residue was purified by column chromatography (90:10 heptanes:ethyl acetate to 20:80) to afford the title compound (66 mg, 56%). $^1$H NMR (400 MHz, $CDCl_3$) δ1.00-1.08 (m, 2H), 1.61 (t, *J* = 3.4 Hz, 2H), 2.47 (dd, *J* = 4.7, 2.6 Hz, 1H), 2.73 (t, *J* = 4.3 Hz, 1H), 2.99-3.04 (m, 1H), 3.19-3.25 (m, 1H), 3.60-3.66 (m ,1H), 5.45 (s, 1H), 7.40-7.30 (m, 4H); MS (ESI): $[M+H]^+$ m/z 252.2.

**(*R*)-1-(4-chlorophenyl)-N-((4-(2,3-dihydro-1H-inden-2-yl)-5-oxomorpholin-2-yl)methyl) cyclopropane-1-carboxamide (OXM1):** To a solution of (*R*)-1-(4-chlorophenyl)-N-(oxiran-2-ylmethyl)cyclopropane-1-carboxamide (850 mg, 3.38 mmol) in toluene (6.75 ml) was added indan-2-amine (450 mg, 3.38 mmol) and the reaction was heated at 100 °C for 1 h. The reaction was then allowed to cool to ambient temperature to which was added triethylamine (0.57 ml, 4.05 mmol) and 2-chloroacetyl chloride (0.27 ml, 3.38 mmol). The reaction was stirred overnight at ambient temperature. To the reaction was added sodium methylate (205 in methanol, 2.32 ml, 10.1 mmol). The reaction was stirred for 1.5 h. The reaction was quenched with 1N HCl (1 ml) and diluted with dichloromethane. The organic layer was separated and the aqueous was extracted with dichloromethane (3×). The organic layers were combined and concentrated under reduced pressure. The crude residue was initially purified by column chromatography (heptanes:ethyl acetate). The resulting material was further purified by SFC (Phenomenex Bipenyl 5 μm 250 × 2.1 mm with $CO_2$ as the mobile phase A; methanol as the mobile phase B; 10% B over 100 min at 80 ml/min flow rate) to provide the title compound as a white solid (578 mg, 40%). HPLC (Kinetic C18 2.6 μm 100 × 3.0 mm with 0.1% formic acid in water (v/v) as the mobile phase A; 0.1% formic acid in acetonitrile (v/v) as the mobile phase B; 5% B for 0.5 min then to 100% B over 4 min and hold for 1.5 min at 0.75 ml/min flow rate) RT 5.173 min; >99%; MS (ESI): $[M+H]^+$ m/z 425.2; Chiral HPLC (Chiral Tech AD-H 5 μm 250 × 4.6 mm with $CO_2$ as the mobile phase A; 0.2% ammonia in methanol (v/v) as the mobile phase B; 5% B for 1.0 min then to 60% B over 8 min and hold for 0.5 min at 3.0 ml/min flow rate) RT 8.699 min; >99 EE ((*S*)-1-(4-chlorophenyl)-*N*-((4-(2,3-dihydro-1H-inden-2-yl)-5-oxomorpholin-2-yl)methyl)cyclopropane-1-carboxamide prepared from (*R*)-1-amino-3-chloropropan-2-ol RT = 7.557 min); $^1$H NMR (400 MHz, $(CD_3)_2SO$) δ 0.89-0.95 (m, 2H), 1.23-1.29 (m, 2H), 2.85-2.90 (m, 1H), 2.94-3.00 (m, 2H), 3.05-3.17 (m, 4H), 3.72-3.79 (m, 1H), 4.07 (dd, *J* = 8.0 8.0 Hz, 2H), 5.27-5.34 (m, 1H), 6.85-6.91 (m, 1H), 7.18-7.27

(m, 6H), 7.30-7.35 (m, 2H). $^{13}C$ NMR (151 MHz, DMSO-$d_6$) δ 14.7, 14.8, 29.5, 34.5, 35.7, 41.1, 43.4, 51.8, 66.8, 71.5, 124.2, 124.3, 126.6, 126.6, 128.5, 131.9, 138.7, 140.9, 140.9, 165.3, 172.3. HRMS (ESI/QTOF) m/z: $[M+H]^+$ Calcd for $C_{24}H_{25}ClN_2O_3$ 425.1627; Found 425.1624; $[\alpha]_D^{23}$ (*c* 0.003, MeOH) = -33.3.

**Preparation of OXM2.**

***tert*-butyl (*S*)-(3-(bicyclo[1.1.1]pentan-1-ylamino)-2-hydroxypropyl)carbamate:** To a solution of bicyclo[1.1.1]pentan-1-amine (16.68 g, 139.4 mmol) in isopropanol (500 ml) was added DIPEA (18 g, 139.3 mmol) followed by a solution of *tert*-butyl (R)-(oxiran-2-ylmethyl)carbamate (24 g, 138.6 mmol) in isopropanol (200 ml) at room temperature. The reaction was stirred for 16 h and then concentrated under reduced pressure. The crude residue was purified by column chromatography (2:8 ethyl acetate in petroleum ether to 100% ethyl acetate) twice to provide the title compound as a yellow solid (7.6 g, 21%). $^1H$ NMR (400 MHz, $CDCl_3$) δ 1.44 (s, 9H), 1.70 – 1.80 (m, 6H), 2.48 – 2.53 (m, 1H), 2.68 – 2.72 (m, 1H), 3.05 – 3.11 (m, 1H), 3.26 – 3.32 (m, 1H), 3.66 – 3.73 (m ,1H), 4.98 (br S, 1H).

***tert*-butyl (*R*)-((4-(bicyclo[1.1.1]pentan-1-yl)-5-oxomorpholin-2-yl)methyl)carbamate:** To a solution of *tert*-butyl (R)-(3-(N-(bicyclo[1.1.1]pentan-1-yl)-2-chloroacetamido)-2-hydroxypropyl)carbamate (7.7 g, 23.1 mmol) in THF (300 ml) at 5 °C was added NaH (60% in mineral oil, 3.7 g, 92.5 mmol) in portions. The reaction was then stirred at room temperature for 5 h. The reaction was cooled to 5 °C and quenched with saturate aqueous $NH_4Cl$ (50 ml) and water (50 ml). The organic layer was separated, and the aqueous layer was extracted with ethyl acetate (50 ml × 2). The combined organic layers were concentrated under reduced pressure. The residue was purified by column chromatography (petroleum ether to 50% ethyl acetate in petroleum ether) to provide the title compound as a white solid (4.34 g, 63%). LCMS (Waters Xbridge C180 30 × 2.1 mm with 0.05% ammonia hydroxide in water (v/v) as the mobile phase A; acetonitrile as the mobile phase B; 0% B to 95% over 0.6 min then 95% B for 0.8 min) RT 2.510 min; >95%; MS (ESI): $[M+H-56]^+$ 241.2; $^1H$ NMR (400 MHz, $CDCl_3$) δ 1.46 (s, 9H), 2.14 - 2.19 (m, 6H), 2.48 (s, 1H), 3.14 - 3.20 (m, 3H), 3.43 - 3.46 (m, 1H), 3.75 – 3.78 (m, 1H), 4.03-4.20 (m, 2 H), 4.91 (br s, 1H).

**(*R*)-6-(aminomethyl)-4-(bicyclo[1.1.1]pentan-1-yl)morpholin-3-one:** To a solution of *tert*-butyl (*R*)-((4-(bicyclo[1.1.1]pentan-1-yl)-5-oxomorpholin-2-yl)methyl)carbamate (100 mg, 0.34 mmol) in methanol (3 ml) was added HCl (4 M in dioxane, 1 ml) at room temperature. The reaction was stirred for 3 h. The reaction was concentrated under reduced pressure to provide the title compound as a yellow oil (80 mg, 100%) which was used without further purification as the presumed HCl salt. LCMS (Waters Xbridge C18 30 × 2.1 mm with 0.05% $NH_3$ in water (v/v) as the mobile phase A; acetonitrile as the mobile phase B; 0% B to 95% over 0.6 min then 95% B for 0.8 min) RT 0.732 min; 86%; MS (ESI): $[M+H]^+$ m/z 197.1.

**(*R*)-N-((4-(bicyclo[1.1.1]pentan-1-yl)-5-oxomorpholin-2-yl)methyl)-1-(4-(trifluoromethyl) phenyl)cyclopropane-1-carboxamide (OXM2):** To a solution of (*R*)-6-(aminomethyl)-4-(bicyclo[1.1.1]pentan-1-yl)morpholin-3-one (1.50 g, 6.44 mmol) in dichloromethane (50 ml) was added 1-(4-trifluoromethyl)cyclopropane-1-carboxylic acid (Aldrich, 1.48 g, 6.44 mmol), triethylamine (2.61 g, 25.8 mmol), and HATU (2.94 mg, 7.73 mmol) at 15 °C. The reaction was allowed to warm to room temperature and stir for 16 h. The reaction mixture was poured into water (15 ml) and extracted with ethyl acetate (50 ml × 3). The combined organic concentrated under reduced pressure.

The residue was first purified by column chromatography (petroleum ether to 33% ethyl acetate in petroleum ether) then purified by HPLC (Phenomenex Gemini C18 10 μm 250 × 50 mm with 0.5% ammonia hydroxide in water (v/v) as the mobile phase A; acetonitrile as the mobile phase B; 35% B to 55% B over 15 min at 110 ml/min flow rate) to provide the title compound as a white solid (1.39 g, 50%). LCMS (Waters Xselect CSH C18 30 × 2.1 mm with 10 mM ammonium acetate in water as the mobile phase A; 10 mM acetonitrile as the mobile phase B; 0% B for 0.6 min then to 100% over 3.4 min) RT 3.098 min; 100%; MS (ESI): $[M+H]^+$ m/z 409.2; Chiral HPLC (Chiralcel OD-3 150×4.6 mm 3um with $CO_2$ as the mobile phase A; 0.05% diethylamine in ethanol as the mobile phase B; 5% to 40% B over 5 mins, 2.5 ml/min) RT 2.705 min; >99 EE.

$^1$H NMR (400 MHz, $CDCl_3$) δ 1.09 - 1.11 (m, 2H), 1.63 – 1.67 (m, 2H), 2.13 – 2.18 (m, 6H), 2.48 (s, 1H), 3.02 – 3.17 (m, 3H), 3.54 - 3.56 (m, 1H), 3.68 – 3.71 (m, 1H), 3.93 – 4.12 (m, 2H), 5.54 – 5.57 (m, 1H), 7.54 (d, *J* = 8.0 Hz, 2H), 7.66 (d, *J* = 8.0 Hz, 2H). $^{19}$F NMR (400 MHz, $CDCl_3$) δ - 62.65 (s). $^{13}$C NMR (151 MHz, DMSO-$d_6$) δ 14.7, 14.8, 23.4, 30.2, 41.2, 46.1, 51.6, 53.0, 66.8, 71.5, 121.6, 123.4, 125.2, 125.3, 125.3, 125.3, 125.4, 127.0, 127.3, 127.5, 127.7, 127.9, 130.6, 144.8, 144.8, 166.2, 171.9.

**Preparation of OXM3**

***tert*-butyl (3-(benzyl(2,3-dihydro-1H-inden-2-yl)amino)-2-hydroxypropyl)carbamate:** To a solution of N-benzyl-2,3-dihydro-1H-inden-2-amine (22.3 g, 100 mmol) in methanol was added *tert*-butyl (oxiran-2-ylmethyl)carbamate (17.3 g, 100 mmol) in methanol. The reaction mixture was refluxed for 2 h, then cooled, and concentrated under reduced pressure. The residue was co-evaporated with dioxane to give the title compound as a yellow oil (30g, 60%) which was used without further purification.

***tert*-butyl (3-((2,3-dihydro-1H-inden-2-yl)amino)-2-hydroxypropyl)carbamate:** To a solution of *tert*-butyl (3-(benzyl(2,3-dihydro-1H-inden-2-yl)amino)-2-hydroxypropyl)carbamate (5.8 g, 15 mmol) in ethyl acetate (100 ml) was added 10% Pd/C (1 g). The mixture was hydrogenated under an atmosphere of hydrogen gas for 2 h. The reaction mixture was filtered through celite. The filtrate was concentrated under reduced pressure to provide the title compound (3g, 80%) as a brown oil and used without further purification.

***tert*-butyl (3-(2-chloro-N-(2,3-dihydro-1H-inden-2-yl)acetamido)-2-hydroxypropyl) carbamate:** To a solution of *tert*-butyl (3-((2,3-dihydro-1H-inden-2-yl)amino)-2-hydroxypropyl) carbamate (30.6 g, 100 mmol) in THF/DMF (1:1) was added diisoprylethylamine (5 ml) and the reaction mixture was cooled to -60 ºC. To the cooled reaction mixture chloroacetyl chloride was added dropwise under an atmosphere of argon gas. The reaction was stirred overnight, purified by column chromatography on silica gel, and concentrated under reduced pressure to provide the title compound (30g, 80%) as a yellow oil.

***tert*-butyl ((4-(2,3-dihydro-1H-inden-2-yl)-5-oxomorpholin-2-yl)methyl)carbamate:** A solution of *tert*-butyl (3-(2-chloro-N-(2,3-dihydro-1H-inden-2-yl)acetamido)-2-hydroxypropyl)carbamate (38 g, 100 mmol) in DMF (300 ml) was cooled and NaH (60% in mineral oil, 5g, 120 mmol) was added in portions. The reaction mixture was allowed to warm to room temperature and stirred for 2 h. Water and ethyl acetate was added and the resulting mixture was concentrated under reduced pressure. The resulting reside was diluted with water and ether. The organic layer was separated, and the aqueous layer was extracted with ether. The combined organic layer was washed with water and brine, then filtered through $Na_2SO_4$. The filtrate was evaporated under reduced pressure and the residue was crystallized with ether with hexanes to provide the title compound (30 g, 60%) as a white solid.

**6-(aminomethyl)-4-(2,3-dihydro-1H-inden-2-yl)morpholin-3-one:** To a solution of *tert*-butyl ((4-(2,3-dihydro-1H-inden-2-yl)-5-oxomorpholin-2-yl)methyl)carbamate (34.6 g, 100 mmol) in dichloromethane (500 ml) was added trifluoroacetic acid. The reaction mixture was stirred for 24 h and then evaporated under reduced pressure. The residue was dissolved in water and made alkaline with saturated $K_2CO_3$. The mixture was extracted with chloroform. The organic layer was dried and evaporated to provide the title compound (20g, 75%).

**2-(4-chlorophenyl)-*N*-((4-(2,3-dihydro-1H-inden-2-yl)-5-oxomorpholin-2-yl)methyl)-2-methylpropanamide (OXM3):** To a solution of 6-(aminomethyl)-4-(2,3-dihydro-1H-inden-2-yl)morpholin-3-one (100 mg, 0.41 mmol) in DCM (6.0 ml) at 0 °C was added 2-(4-chlorophenyl)-2-methylpropanoic acid (81 mg, 0.41 mmol), triethylamine (0.113 ml, 0.81 mmol), and HATU (232 mg, 0.61 mmol). The reaction was stirred for 16 h at 15 °C. The reaction mixture was quenched the addition of aqueous potassium bicarbonate (20 ml). The layers were separated and the aqueous was washed with ethyl acetate (40 ml × 3). The organic layers combined and were dried with sodium sulfate, filtered, and concentrated under reduced pressure. The crude residue was purified by HPLC (DuraShell 5 μm 150 × 25 mm with 0.05% ammonia hydroxide in water (v/v) as the mobile phase A; acetonitrile as the mobile phase B; 46% B to 76% B over 10 min at 25 ml/min flow rate) to provide the title compound as a white solid (48 mg, 28%). LCMS (Waters Xselect CSH C18 5 μm 50 × 2.1 mm with 5% 10 mM ammonia acetate in water (v/v) as the mobile phase A; acetonitrile as the mobile phase B; 0% B for 0.6 min to 100% B over 3.4 min at 0.8 ml/min flow rate ) RT 3.197 min; >99%; MS (ESI): $[M+H]^+$ m/z 427.4; HRMS (ESI/QTOF) m/z: $[M+H]^+$ Calcd for $C_{24}H_{27}ClN_2O_3$ 427.1783; Found 427.1778: $^1$H NMR (400 MHz, ($CD_3OD$) δ 1.47 (s, 6H), 2.87-2.93 (m, 1H), 3.00-3.07 (m, 2H), 3.09-3.23 (m, 3H), 3.26 (d, *J* = 6.0 Hz, 2 H), 3.75-3.81 (m, 1H), 4.10 (d, *J* = 16.6 Hz, 1H), 4.21 (d, *J* = 16.6 Hz, 1H), 5.36-5.44 (m, 1H), 7.14-7.18 (m, 2H), 7.22-7.25 (m, 2H), 7.26-7.30 (m, 4 H). $^{13}$C NMR (151 MHz, DMSO-$d_6$) δ 14.7, 14.8, 23.4, 30.2, 41.2, 46.1, 51.6, 53.0, 66.8, 71.5, 121.6, 123.4, 125.2, 125.3, 125.3, 125.3, 125.4, 127.0, 127.3, 127.5, 127.7, 127.9, 130.6, 144.8, 144.8, 166.2, 171.9.

**Preparation of OXM4**

***N*-((4-(2,3-dihydro-1*H*-inden-2-yl)-5-oxomorpholin-2-yl)methyl)-1-phenylcyclopropane-1-carboxamide (OXM4)**: To a solution of 1-phenylcyclopropane-1-carboxylic acid (19.4 mg, 120 µm) in DCM (~0.1 ml) in a reaction vial was added 6-(aminomethyl)-4-(2,3-dihydro-1H-inden-2-yl)morpholin-3-one (24 mg, 0.1 mmol) in DCM (800 µL) and diisopropylethylamine (~50 µL, 300 mmol). A 0.55 M stock solution of HOPO and EDCI in DCM was separately prepared of which 200 ml (110 µmol) was added to the reaction. The reaction vial was capped and agitated at 50 °C for 16 h. The reaction was concentrated under reduced pressure. The crude residue was purified by HPLC (DuraShell 5 µm 150 × 25 mm with 0.225% formic acid in water (v/v) as the mobile phase A; acetonitrile as the mobile phase B; 9% B to 59% B over 12 min at 35 ml/min flow rate) to provide the title compound as a white solid (10 mg, 26%). LCMS (Waters Xselect CSH C18 5 µm 50 × 2.1 mm with 0.0375% trifluoroacetic acid in water (v/v) as the mobile phase A; 0.01875% trifluoracetic acid in acetonitrile as the mobile phase B; 1% B to 5% over 0.6 min then to 100% B over 3.4 min at 0.8 ml/min flow rate ) RT 3.056 min; >99%; MS (ESI): $[M+H]^+$ m/z 391; $^1$H NMR (600 MHz, $(CD_3)_2SO$) δ 0.93 (d, *J* = 3.1 Hz, 2H), 1.30–1.21 (m, 2H), 2.86 (dd, *J* = 16.4, 6.4 Hz, 1H), 3.01 – 2.92 (m, 3H), 3.06 (dd, *J* = 16.4, 8.6 Hz, 1H), 3.13 – 3.09 (m, 1H), 3.14 (t, *J* = 6.1 Hz, 2H), 3.81 – 3.69 (m, 1H), 4.14 – 3.96 (m, 2H), 5.30 (tt, *J* = 8.5, 6.5 Hz, 1H), 6.71 (t, *J* = 6.0 Hz, 1H), 7.21 – 7.16 (m, 2H), 7.28 – 7.22 (m, 5H), 7.34 – 7.28 (m, 2H); $^{13}$C NMR (151 MHz, DMSO) δ 14.6, 14.7, 30.2, 34.5, 35.7, 36.0, 41.1, 43.4, 44.9, 51.9, 66.8, 71.6, 124.2, 124.3, 126.6, 126.6, 126.7, 127.3, 128.6, 130.0, 139.8, 140.9, 140.9, 165.4, 172.7.

### Supplementary Figures

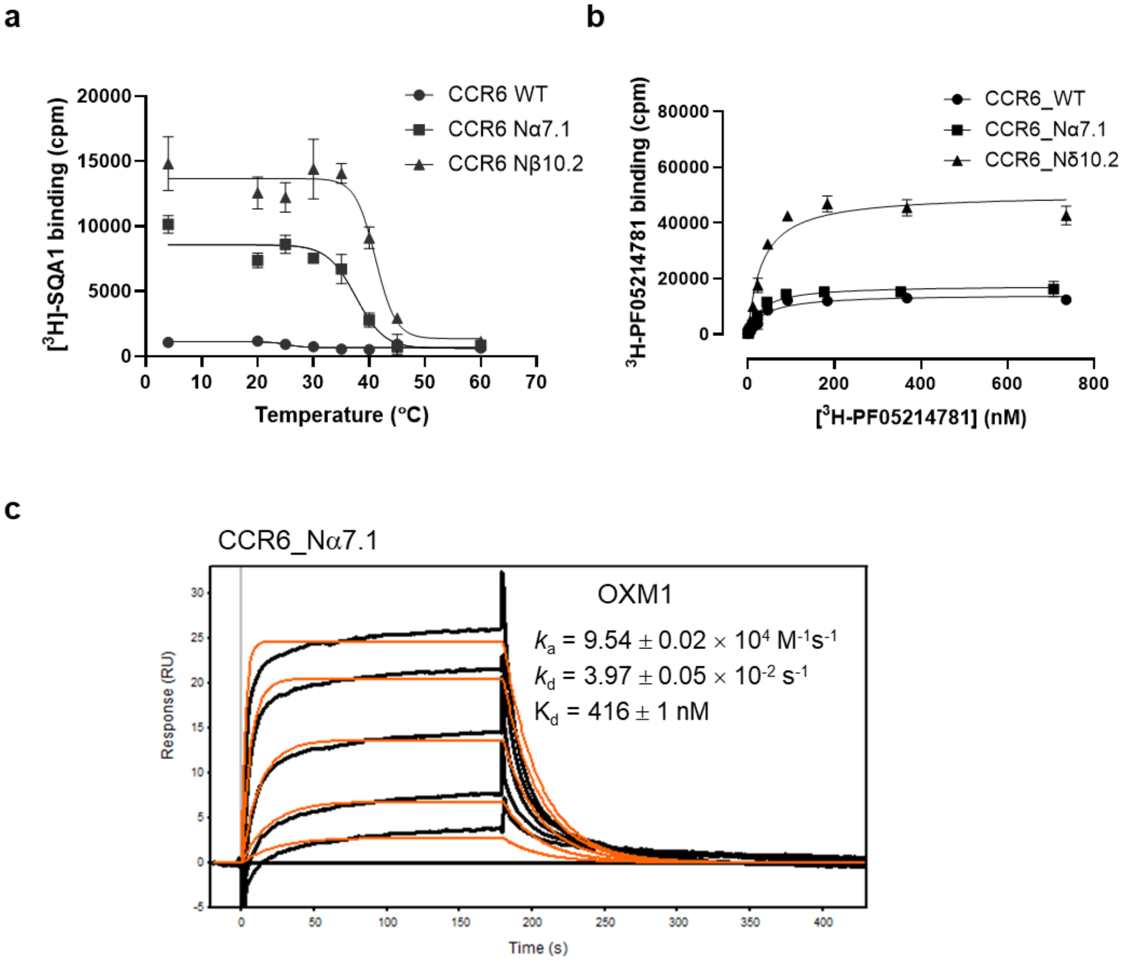

**Figure S1. Thermostability and characterization of CCR6 constructs. a**, Thermostability of human WT and thermostabilized CCR6 constructs in DM with 5 mM MgCl$_2$ are shown. The individual temperature data points are shown in the format of mean ± s.d derived from three independent measurements (*n* = 3). WT and thermostabilized CCR6 produced a mean T$_m$ of 25.2 ± 3.7 °C (WT), 37.6 ± 0.7 °C (Nα7.1), and 41.1 ± 0.4 °C (Nβ10.2). **b**, Saturation binding curves of [$^3$H]-SQA1 to the membranes of HEK293 cells that transiently expressed human WT or thermostabilized CCR6 constructs. Data plotted with average un-transfected control subtracted in the format of mean ± s.d derived from four independent measurements (*n* = 4). There was no significant difference in the affinity of [$^3$H]-SQA1 at WT (Kd = 39.2 ± 4.1 nM), Nα7.1 (Kd = 30.7 ± 1.2 nM), or Nβ10.2 (Kd = 32.8 ± 2.5 nM) CCR6 (*n* = 4). **c**, SPR sensorgrams of OXM1 binding to purified protein of thermostabilized CCR6. A concentration series of OXM1 consisting of 6 three-fold dilutions from 10 to 0.014 µM are shown in black curves (experimental data), fitting using a model of 1:1 binding (orange curves). Data plotted with average non-specific binding control subtracted. Kd in mean ± s.d. and binding kinetics derived from *n* = 3 independent repeats are shown in the figure.

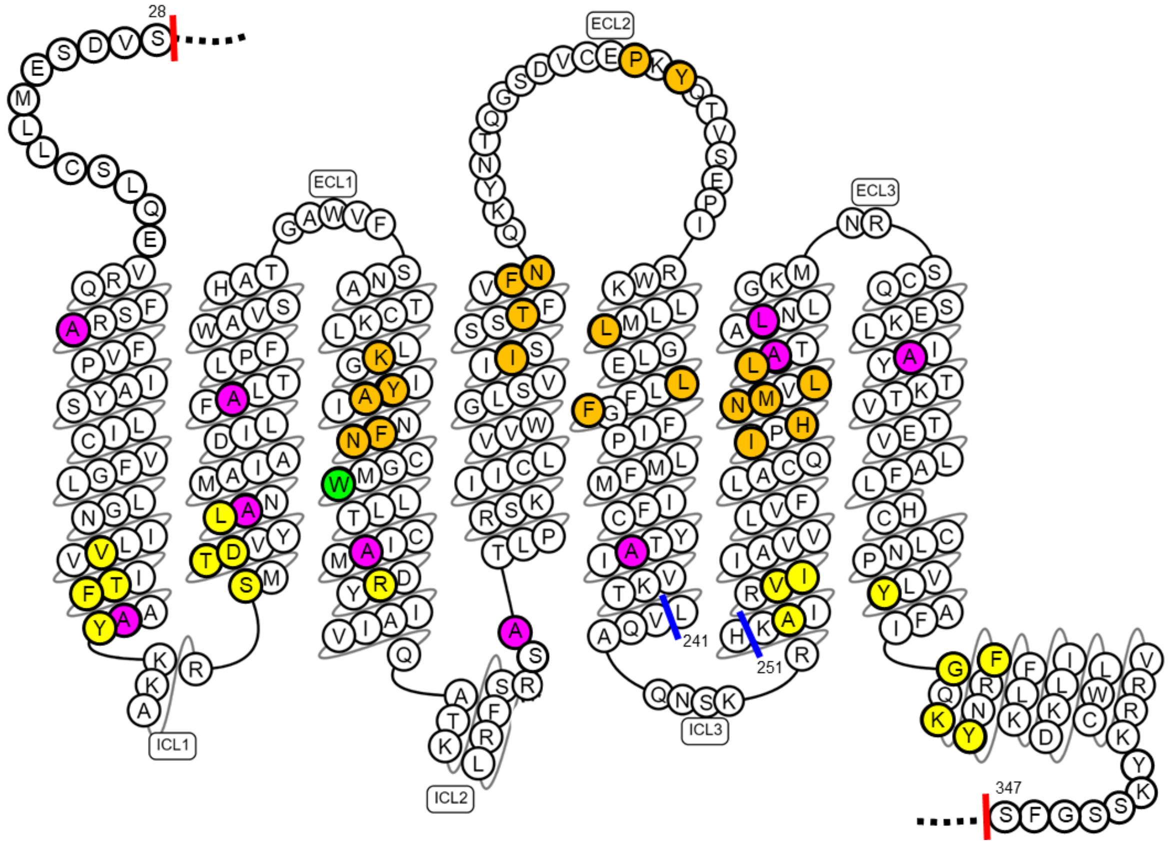

**Figure S2. Snakeplot of the thermostabilized CCR6 constructs.** The thermostabilizing substituations, L47$^{1.33}$A, F73$^{1.59}$A, L86$^{2.44}$A, V96$^{2.54}$A, S140$^{3.47}$A, R159A, F236$^{5.60}$A, V276$^{6.57}$A, A279$^{6.60}$L, and G295$^{7.32}$A are highlighted in magenta. The L134$^{3.41}$W mutation is highlighted in green. Residues interacting with OXM or SQA analogues are highlighted in orange or yellow, repectively. Deletions at ICL3 where the seqeunce of BRIL was inserted are highlighted by blue lines. The termini truncations are highlighted by red lines. The truncated N- and C-termini (Δ1-27, Δ348-374) are not included in the schematic representation. Instead, they are replaced by black dashes.

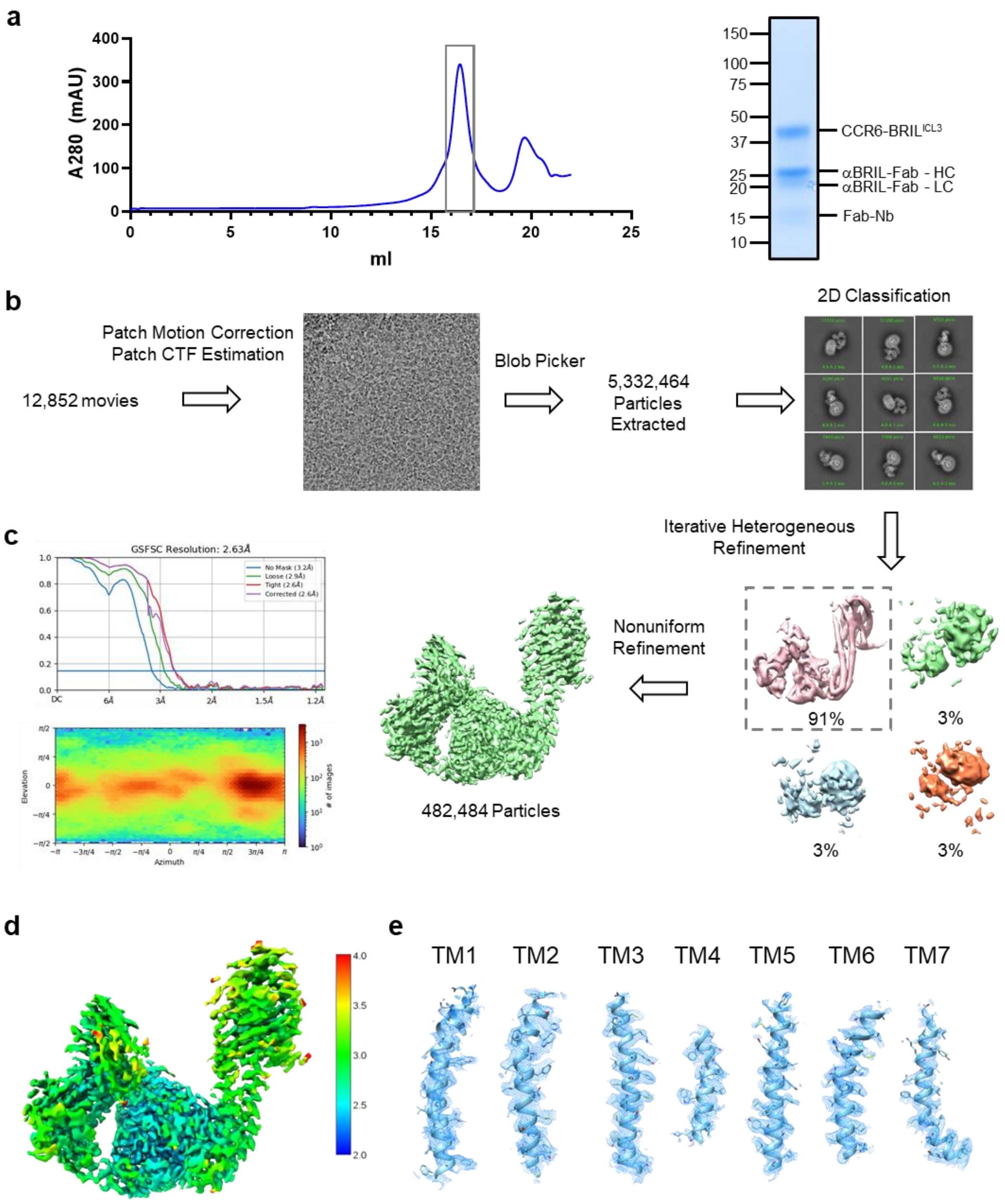

**Figure S3. Cryo-EM data processing workflow for CCR6/SQA1/OXM1.** **a**, SEC profile of CCR6/SQA1/OXM1 complex purification. Fractions used in cryo-EM study are highlighted by the grey box and the corresponding SDS-PAGE image is shown. **b**, Data processing workflow. **c**, Fourier shell correlation curves from gold-standard refinement and particle angular distribution. **d**, Cryo-EM map colored by local resolution. **e**, CCR6 TM helices fitted into the cryo-EM density map.

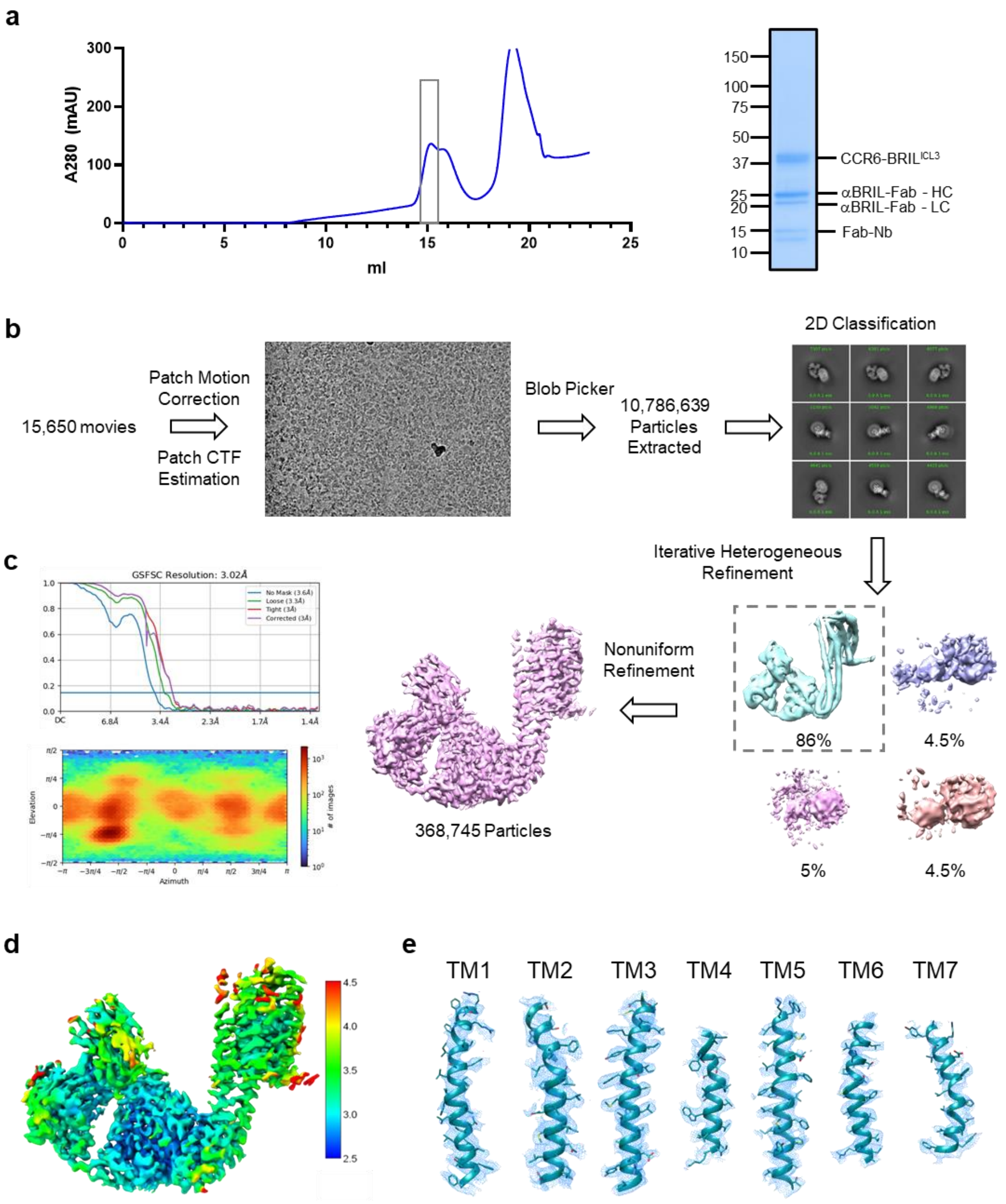

**Figure S4. Cryo-EM data processing workflow for CCR6/SQA1/OXM2. a**, SEC profile of CCR6/SQA1/OXM2 complex purification. Fractions used in cryo-EM study are highlighted by the grey box and the corresponding SDS-PAGE image is shown. **b**, Data processing workflow. **c**, Fourier shell correlation curves from gold-standard refinement and particle angular distribution. **d**, Cryo-EM map colored by local resolution. **e.** CCR6 TM helices fitted into the cryo-EM density map.

| Residue number | 3 | 3 | 3 | 3 | 3 | 4 | 4 | 4 | 4 | 45 | 5 | 5 | 5 | 6 | 6 | 6 | 6 | 6 | 6 |
|---|---|---|---|---|---|---|---|---|---|---|---|---|---|---|---|---|---|---|---|
| Sequence-based () | .29 | .32 | .33 | .36 | .37 | .56 | .60-61 | .63-64 | .64 | .52 | .39-40 | .43-44 | .47 | .49 | .51 | .52 | .53 | .55 | .56 |
| Structure-based (GPCRdb) | x29 | x32 | x33 | x36 | x37 | x56 | x61 | x64 | x65 | x52 | x40 | x44 | x47 | x49 | x51 | x52 | x53 | x55 | x56 |
| [Human] CCR1 | S | Y | Y | L | Y | A | G | F | S | L | A | N | L | T | Y | N | L | I | L |
| [Human] CCR2 | T | Y | H | Y | F | A | G | F | T | P | T | N | L | T | Y | N | I | I | L |
| [Human] CCR3 | S | Y | H | L | Y | A | E | F | Y | A | T | T | L | T | Y | N | V | I | L |
| [Human] CCR4 | S | Y | L | F | Y | A | G | F | S | T | S | N | L | T | Y | N | I | L | F |
| [Human] CCR5 | T | Y | F | F | F | A | G | F | T | S | T | V | L | A | Y | N | I | L | L |
| [Human] CCR6 | K | Y | A | F | N | I | T | F | N | P | L | L | F | I | H | N | M | L | L |
| [Human] CCR7 | F | Y | K | F | F | L | E | Y | S | L | Q | M | F | L | Y | N | G | V | L |
| [Human] CCR8 | S | Y | Y | F | Y | A | L | F | Y | S | N | N | L | V | F | N | V | L | F |
| [Human] CCR9 | N | Y | K | F | Y | L | E | Y | S | M | L | V | F | F | Y | N | C | L | L |
| [Human] CCR10 | S | Y | S | F | H | L | A | F | S | L | A | V | F | L | Y | S | L | L | L |
| [Human] CXCR1 | S | K | E | F | Y | L | F | F | R | E | R | H | F | L | Y | N | L | L | L |
| [Human] CXCR2 | S | K | E | F | Y | L | V | F | R | E | R | Q | F | L | Y | N | L | L | L |
| [Human] CXCR3 | G | F | N | F | Y | F | D | F | L | Y | R | L | F | T | Y | H | L | V | L |
| [Human] CXCR4 | H | Y | T | L | Y | L | D | F | A | R | Q | I | L | L | Y | Y | I | I | S |
| [Human] CXCR5 | I | H | K | F | Y | L | E | F | A | F | R | H | F | S | Y | H | I | I | F |
| [Human] CXCR6 | L | Y | T | F | Y | V | Q | Y | G | Y | L | M | F | M | F | N | L | K | F |
| [Human] $CX_3CR1$ | T | F | F | F | F | V | Q | F | T | G | N | N | F | T | Y | N | V | I | F |
| [Human] XCR1 | N | F | S | L | Y | S | T | H | - | Y | V | N | F | G | Y | N | F | L | F |
| [Human] ACKR1 | Y | W | Y | A | F | L | V | A | S | L | A | V | L | W | H | G | V | L | G |
| [Human] ACKR2 | S | Y | T | F | Y | V | D | F | V | A | R | N | F | F | Y | N | L | L | F |
| [Human] ACKR3 | H | F | S | L | F | V | D | Y | L | S | E | V | F | L | Y | H | V | V | L |
| [Human] ACKR4 | S | Y | T | F | V | L | Q | F | Y | P | Q | I | F | L | Y | N | I | K | F |
| [Human] CCRL2 | I | Y | F | L | Y | A | E | V | Y | F | T | N | L | A | Y | N | I | F | F |

**Figure S5. Structure-based sequence alignment of human chemokine receptors at the OXM binding pocket.** The sequence alignment was performed using servers from www.GPCRdb.org[55].

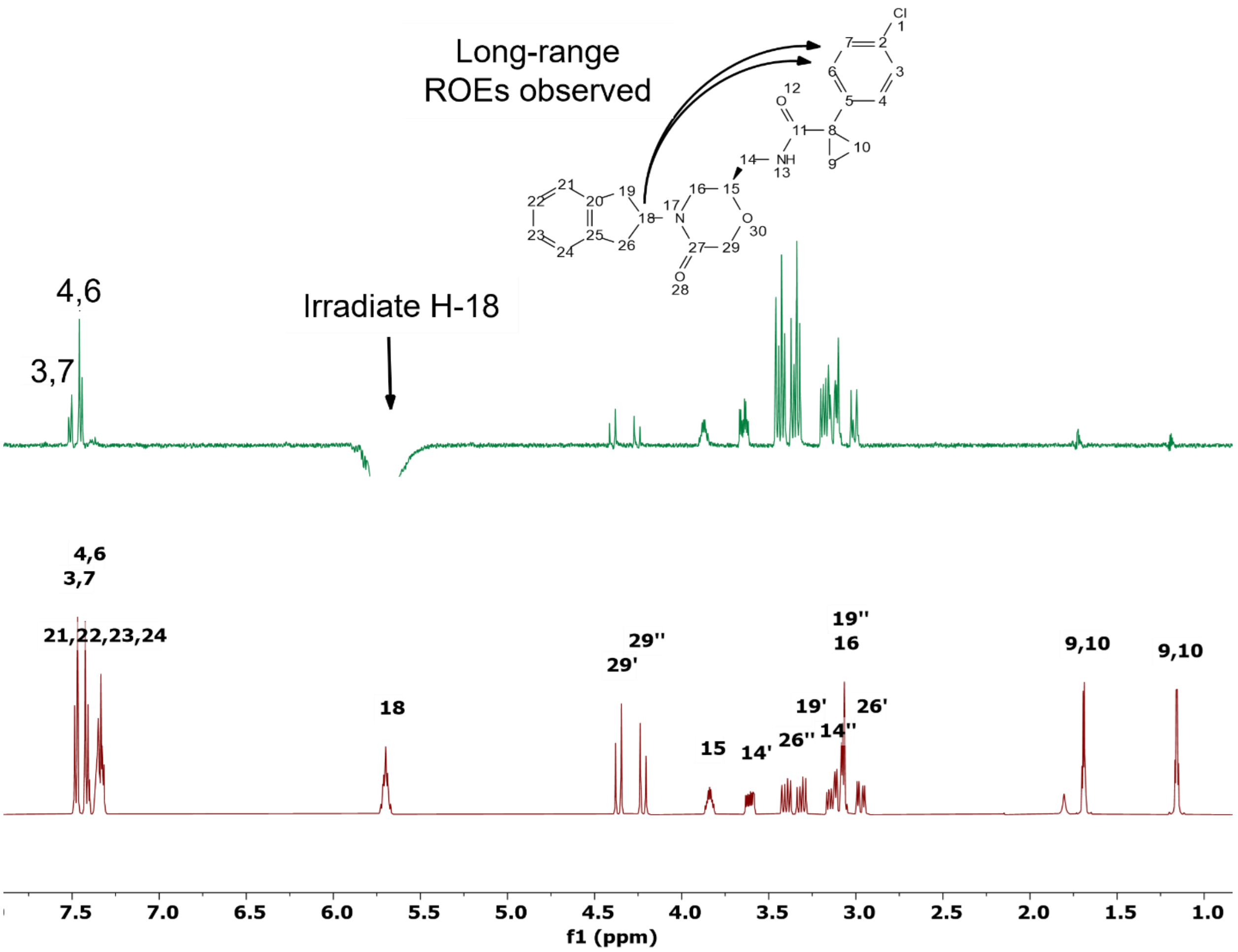

**Figure S6. Rotating-frame Overhauser Enhancement signals (ROEs) observed for OXM1 in $CDCl_3$.** The observation of long-range ROEs from H-18 to H-4/H-6 and to H3/H-7 suggests that OXM1 adopts a U-shaped conformation in solution.

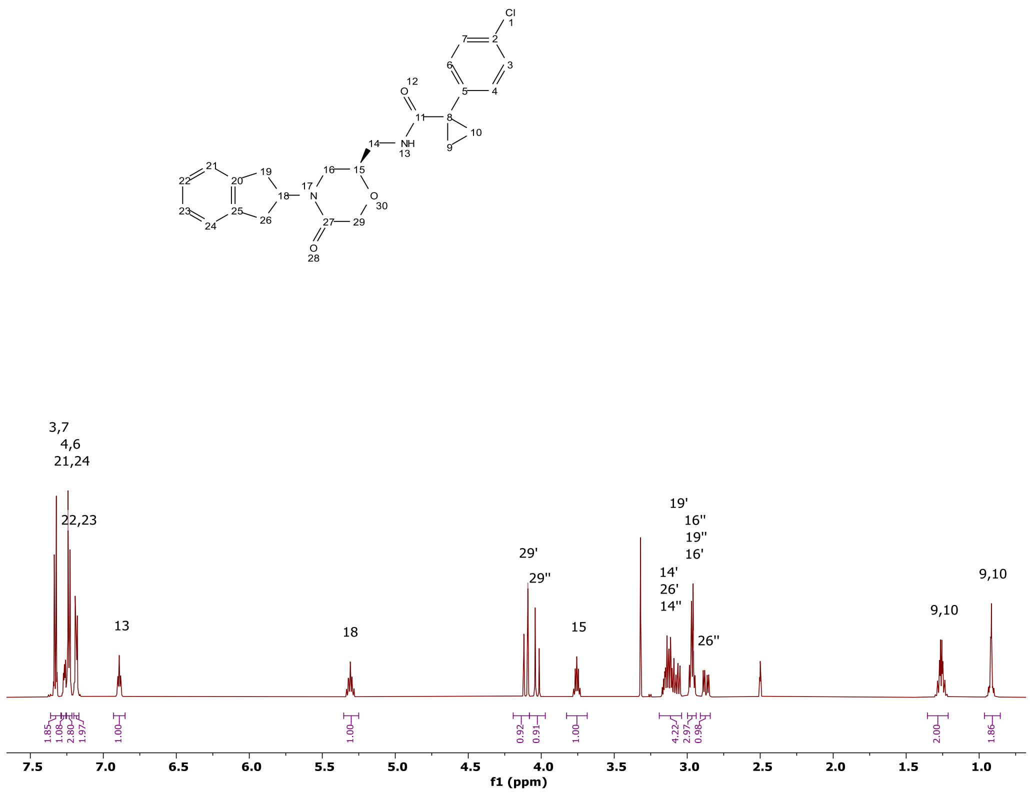

**Figure S7. 600.1 MHz $^1$H spectrum of OXM1 in DMSO-$d_6$.**

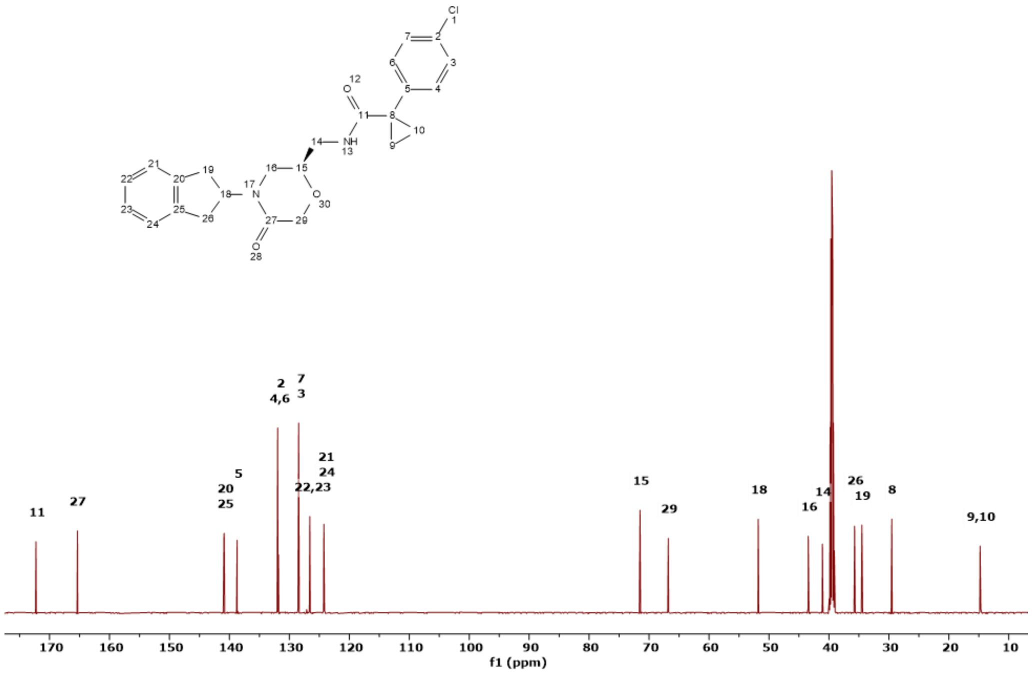

**Figure S8. 150.9 MHz $^{13}$C spectrum of OXM1 in DMSO-$d_6$.**

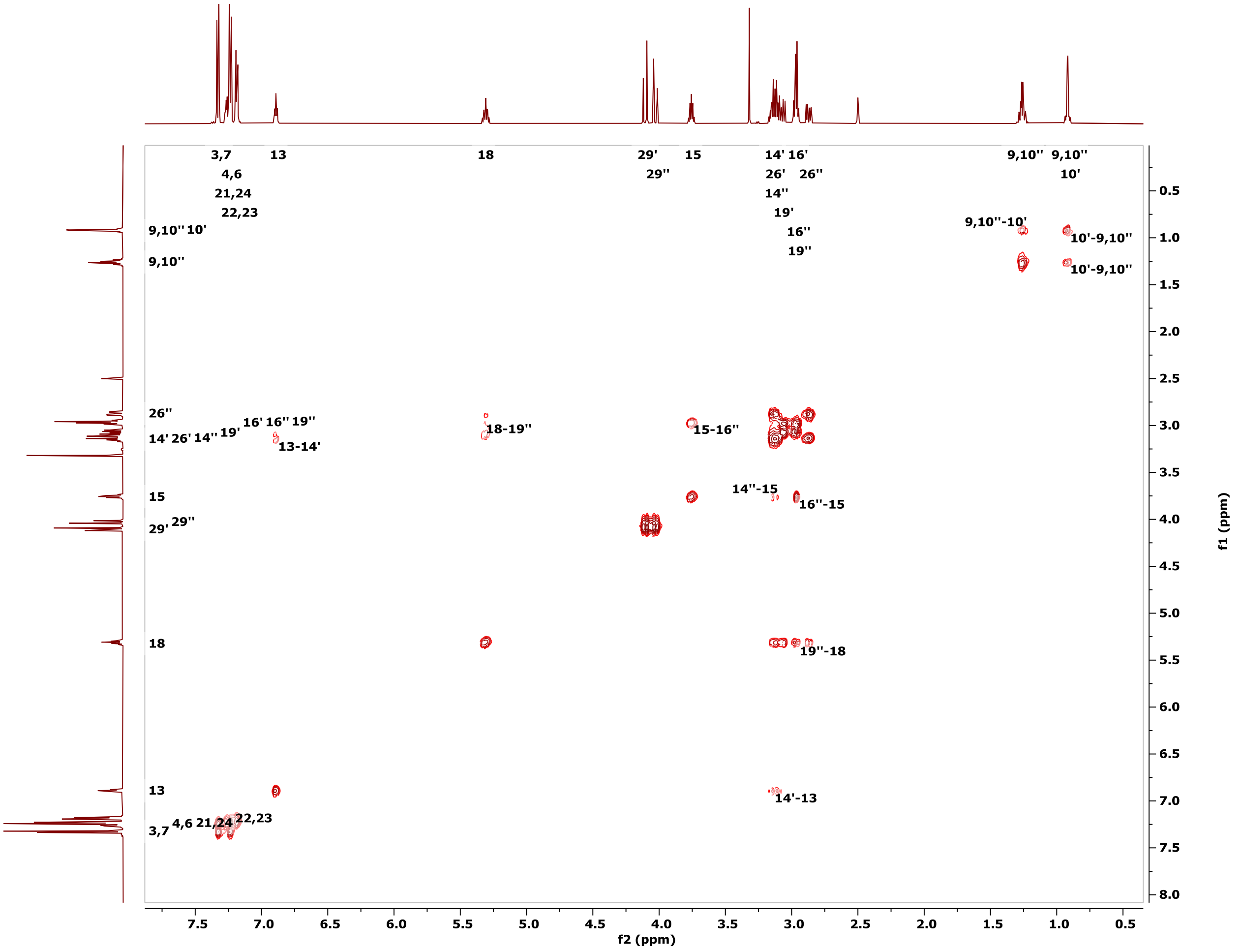

**Figure S9. 600.1 MHz COSY spectrum of OXM1 in DMSO-d6.**

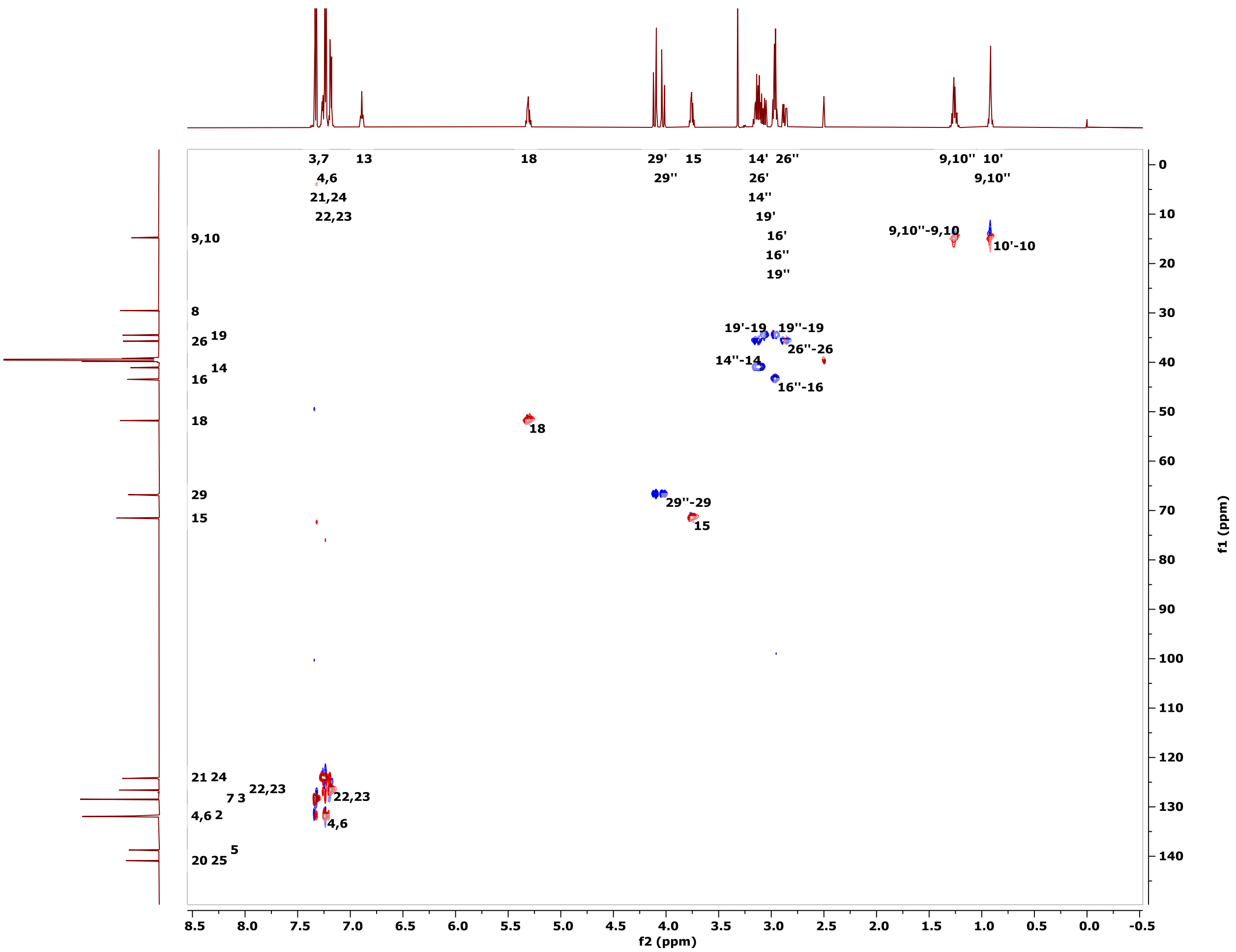

**Figure S10. 600.1 MHz HC-HSQC spectrum of OXM1 in DMSO-d6.**

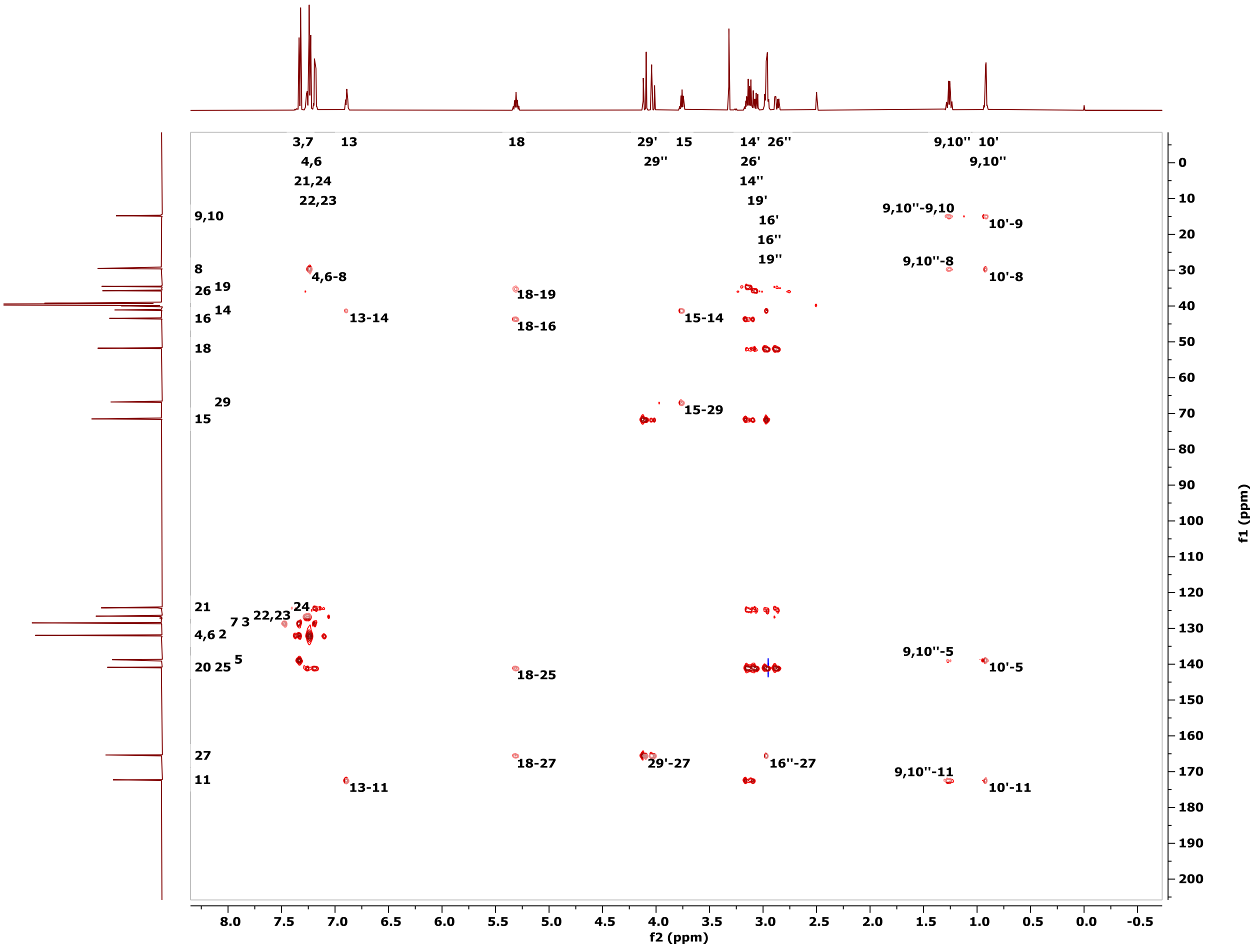

**Figure S11. 600.1 MHz HC-HMBC spectrum of OXM1 in DMSO-d6.**

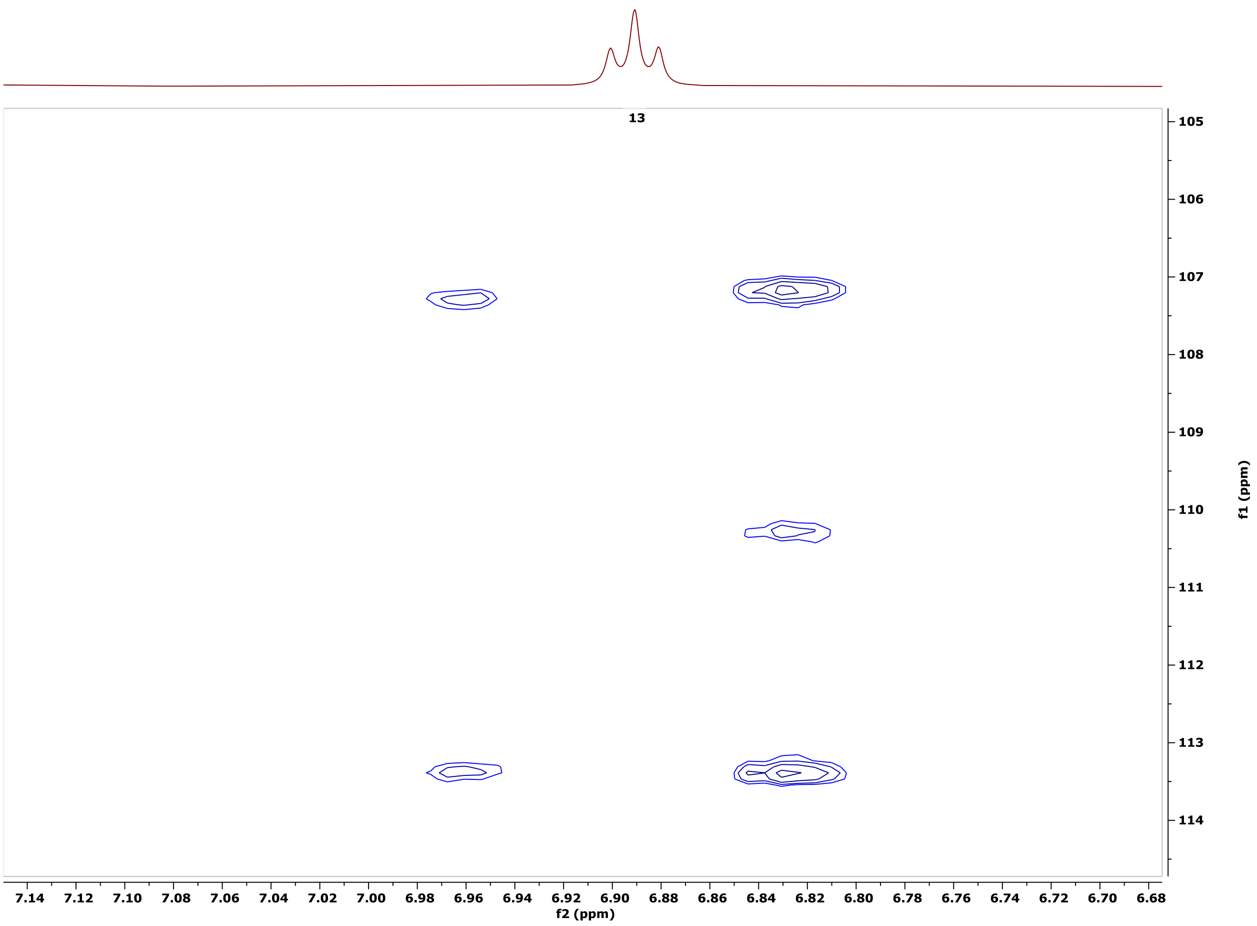

**Figure S12. 600.1 MHz JSBHN-HSQC spectrum of OXM1 in poly-HEMA gel in DMSO-$d_6$.**

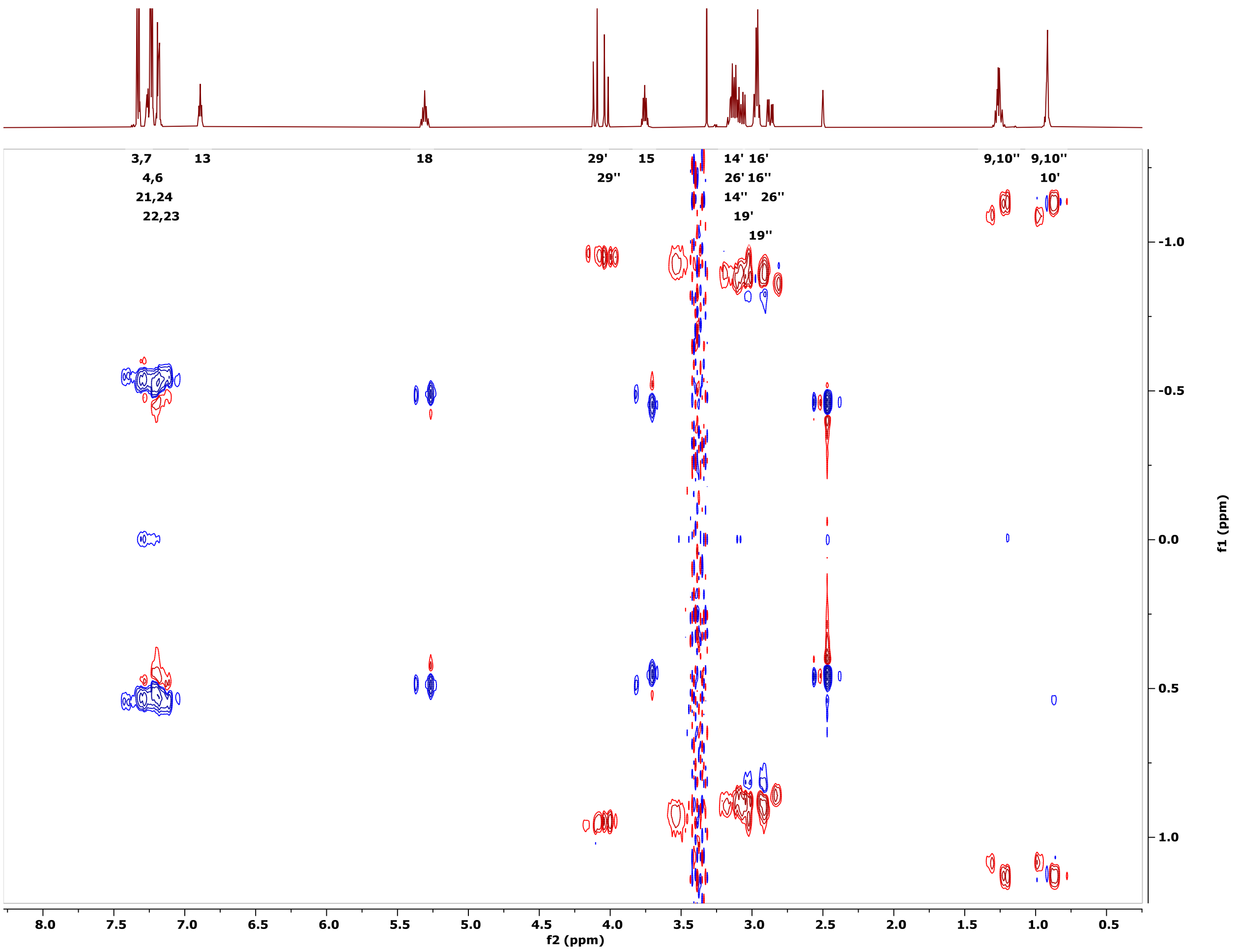

**Figure S13. 600.1 MHz JSBHC-HSQC spectrum of OXM1 in poly-HEMA gel in DMSO-d6.**

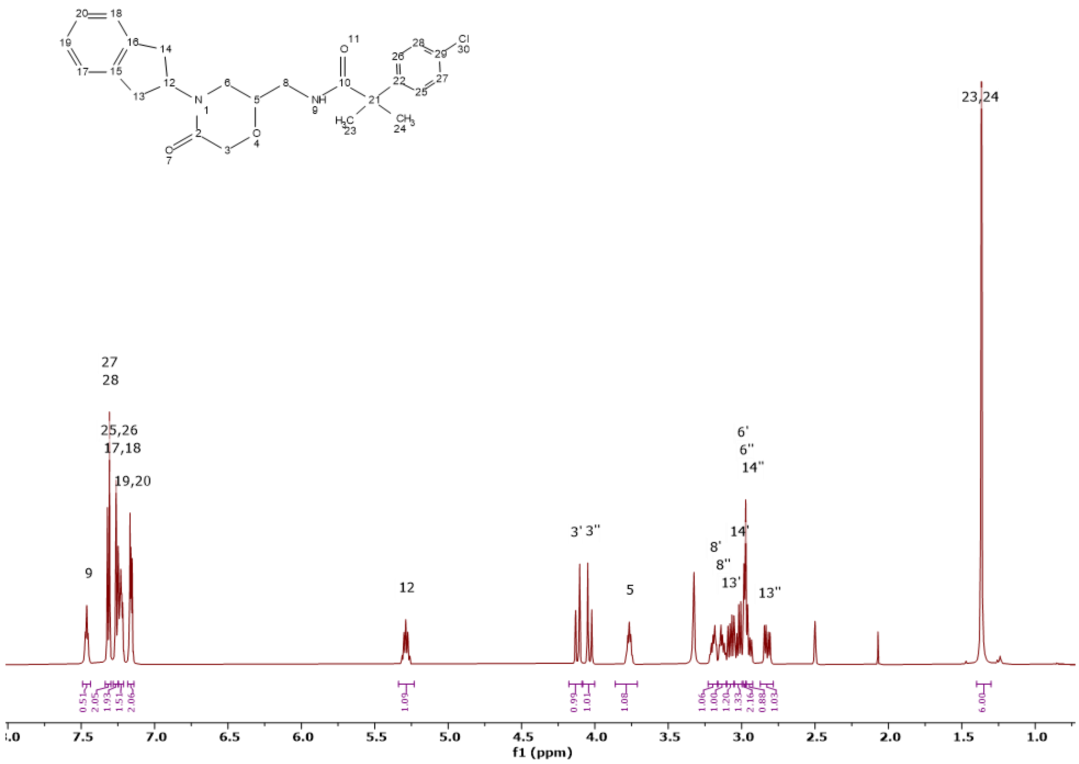

**Figure S14. 600.1 MHz $^{1}$H spectrum of OXM3 in DMSO-d$_6$.**

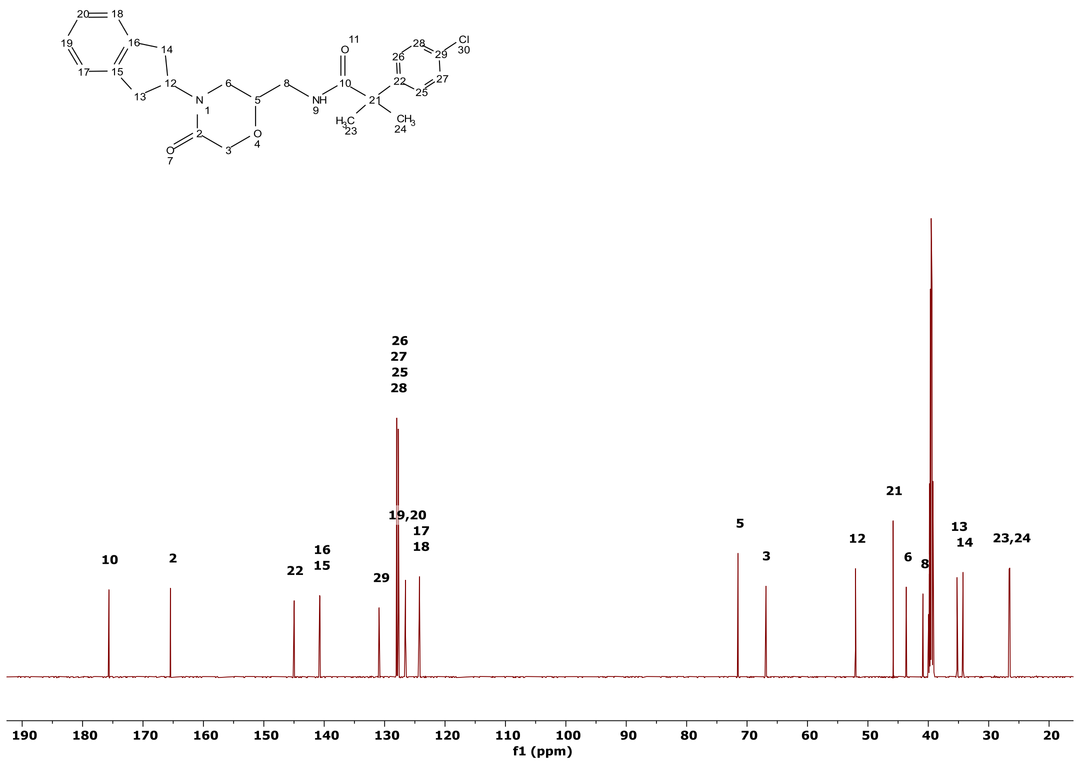

**Figure S15. 150.9 MHz $^{13}$C spectrum of OXM3 in DMSO-d$_6$.**

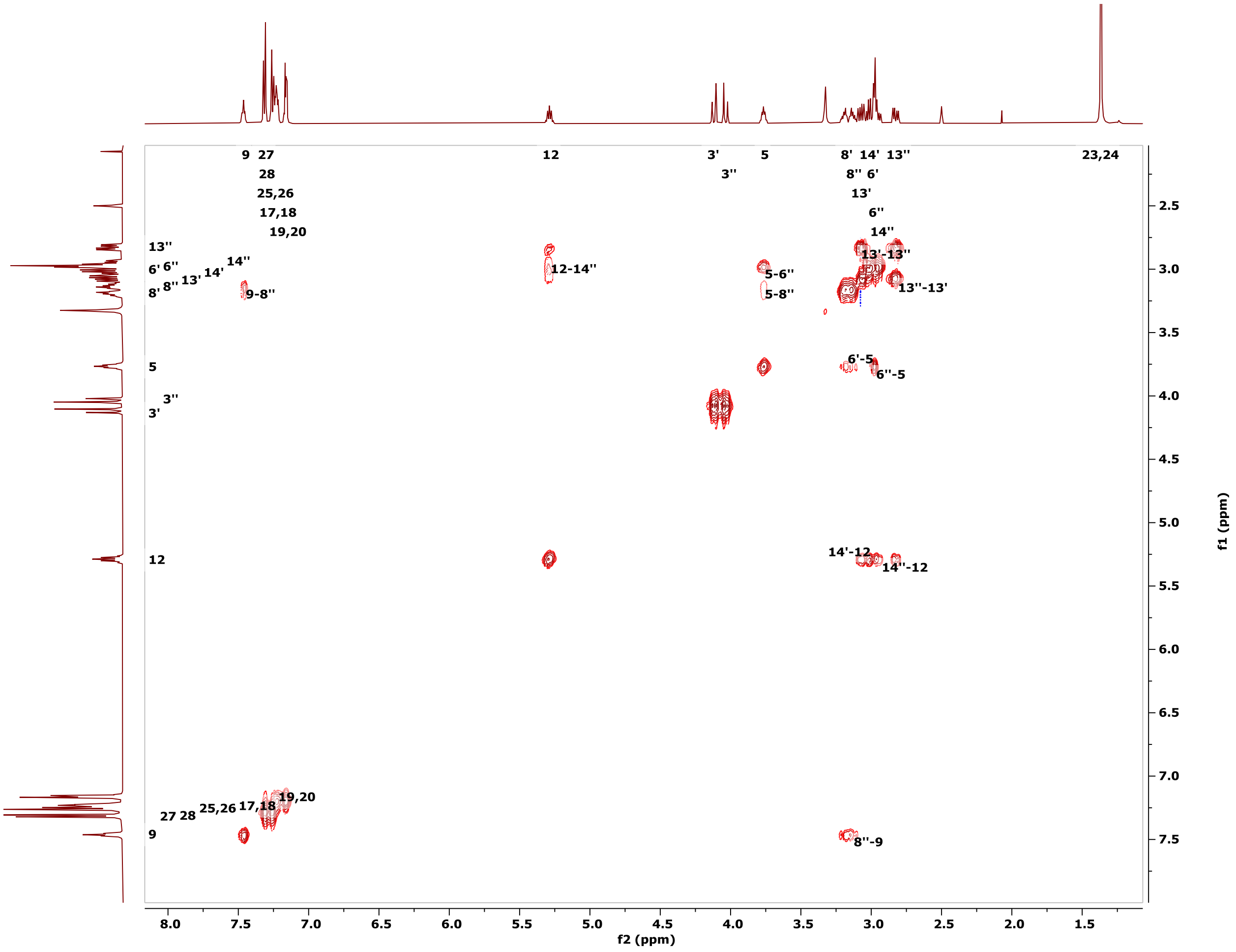

**Figure S16. 600.1 MHz COSY spectrum of OXM3 in DMSO-d6.**

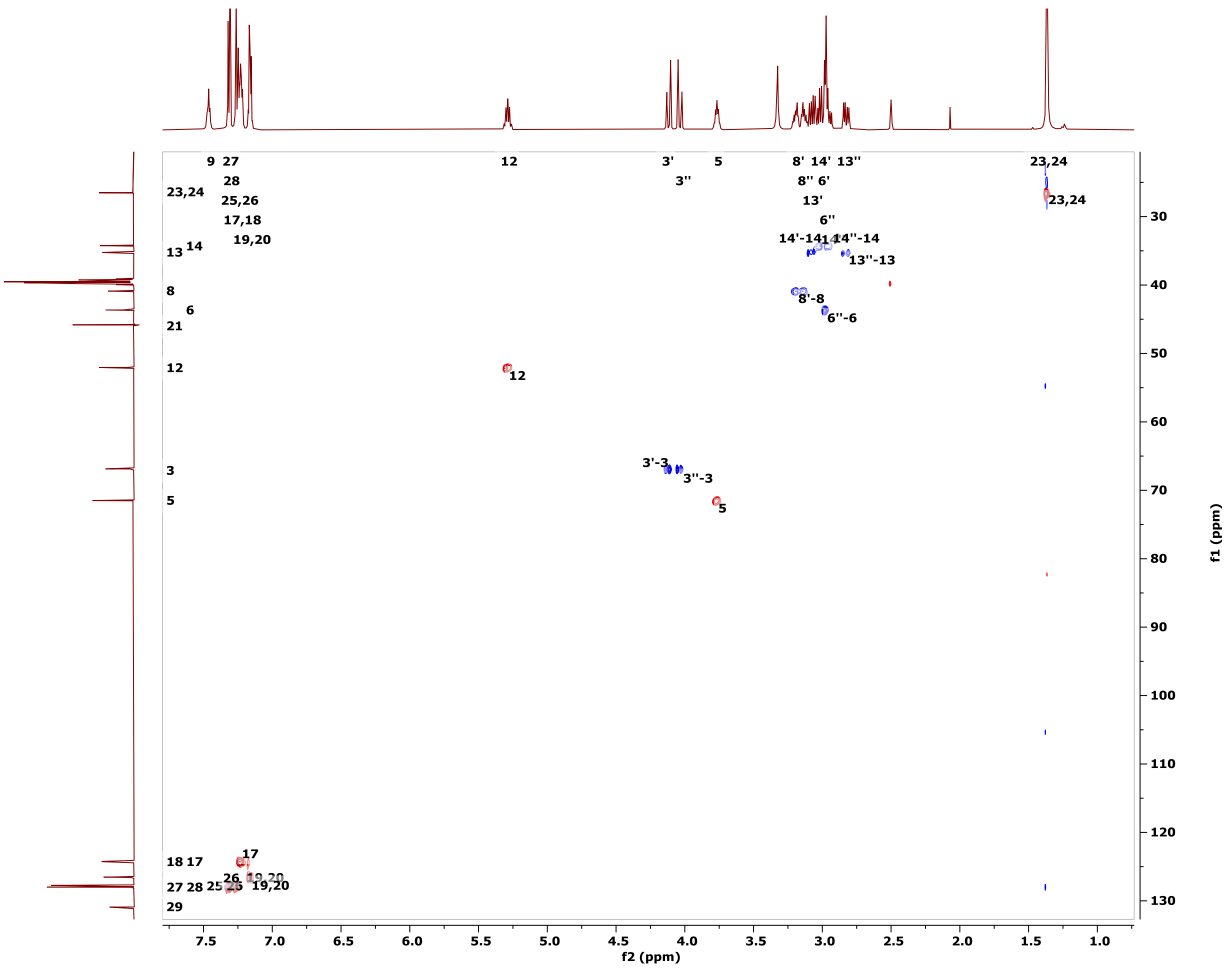

**Figure S17. 600.1 MHz HC-HSQC spectrum of OXM3 in DMSO-d6.**

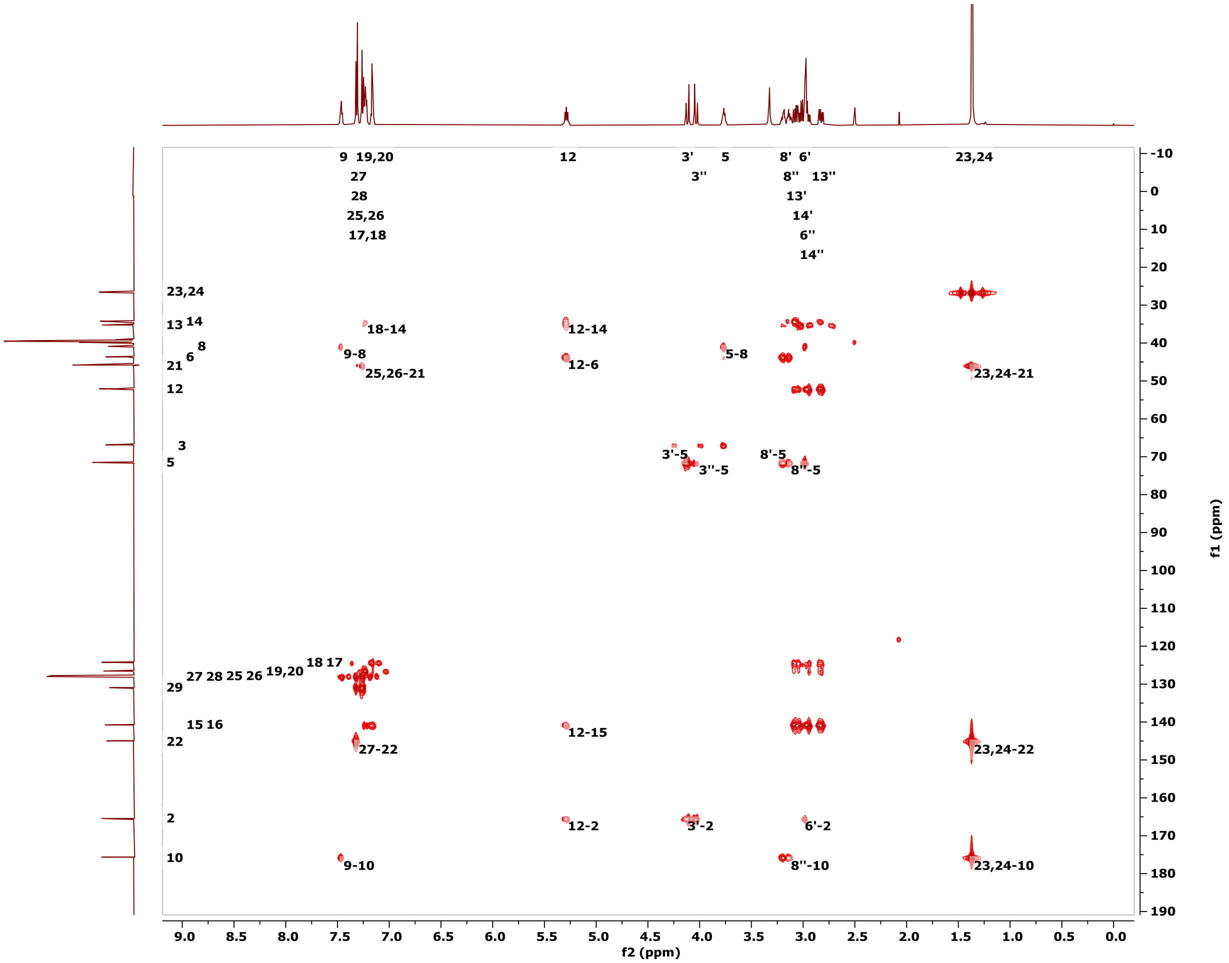

**Figure S18. 600.1 MHz HC-HMBC spectrum of OXM3 in DMSO-d6.**

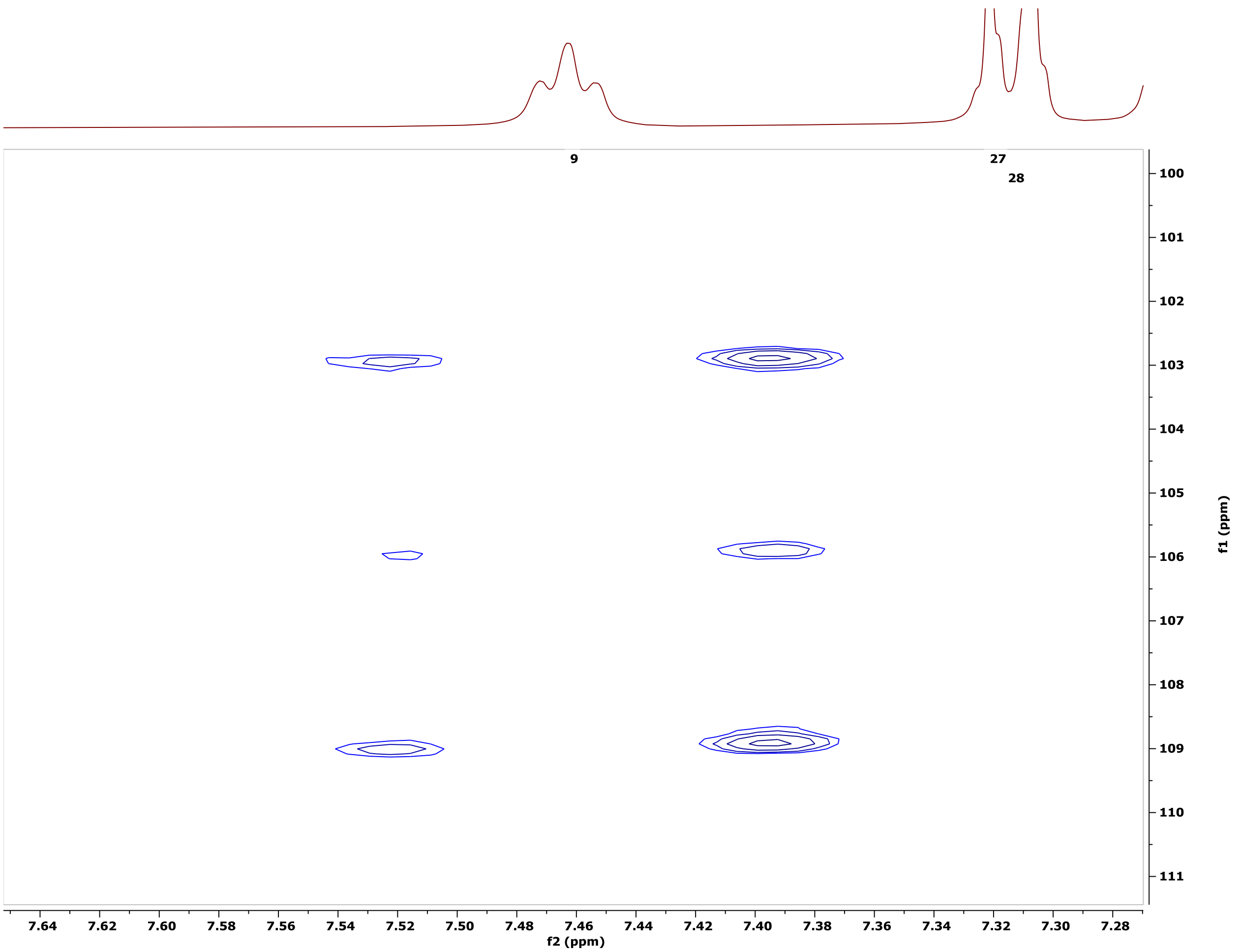

**Figure S19. 600.1 MHz JSBHN-HSQC spectrum of OXM3 in poly-HEMA gel in DMSO-$d_6$.**

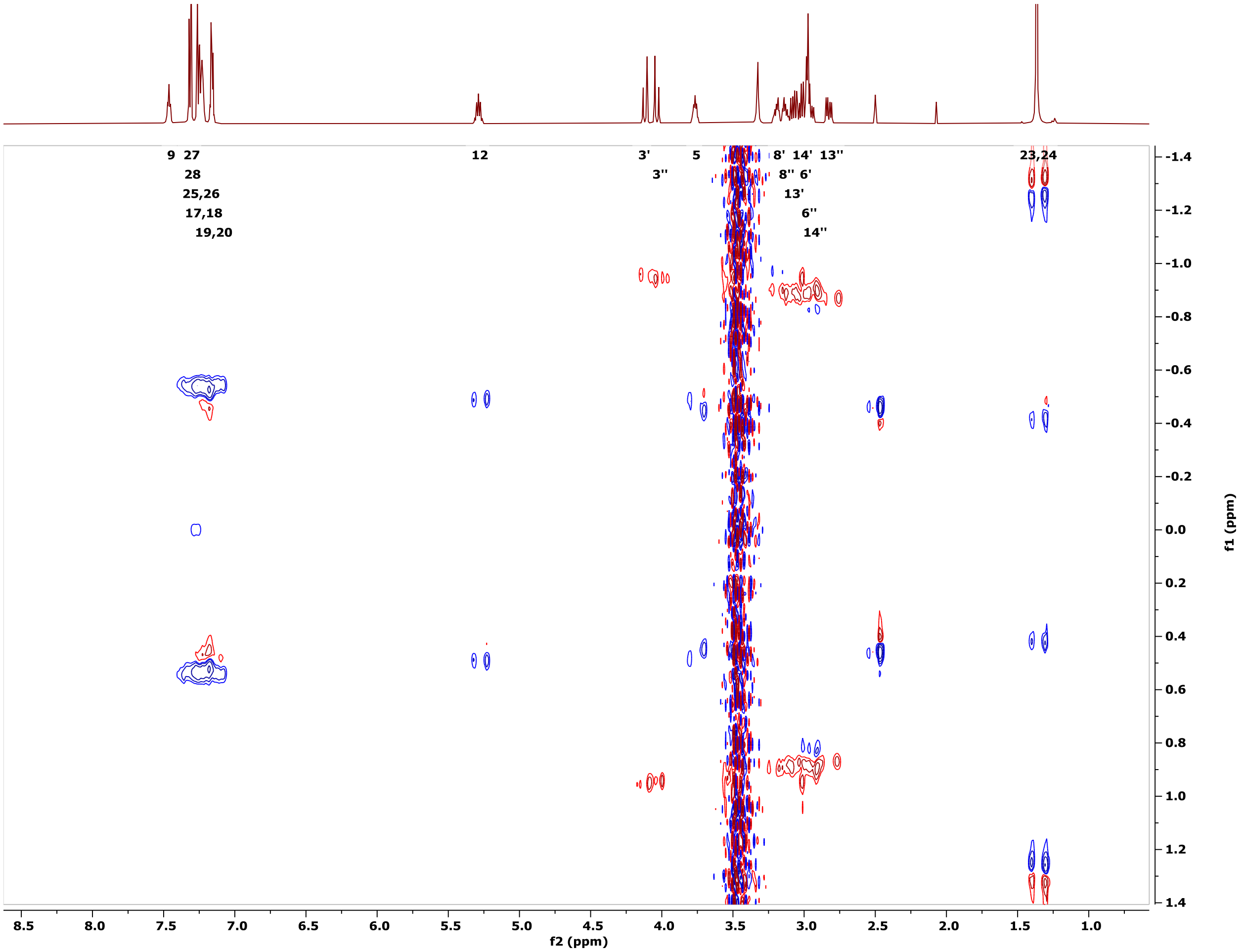

**Figure S20. 600.1 MHz JSBHC-HSQC spectrum of OXM3 in poly-HEMA gel in DMSO-d6.**

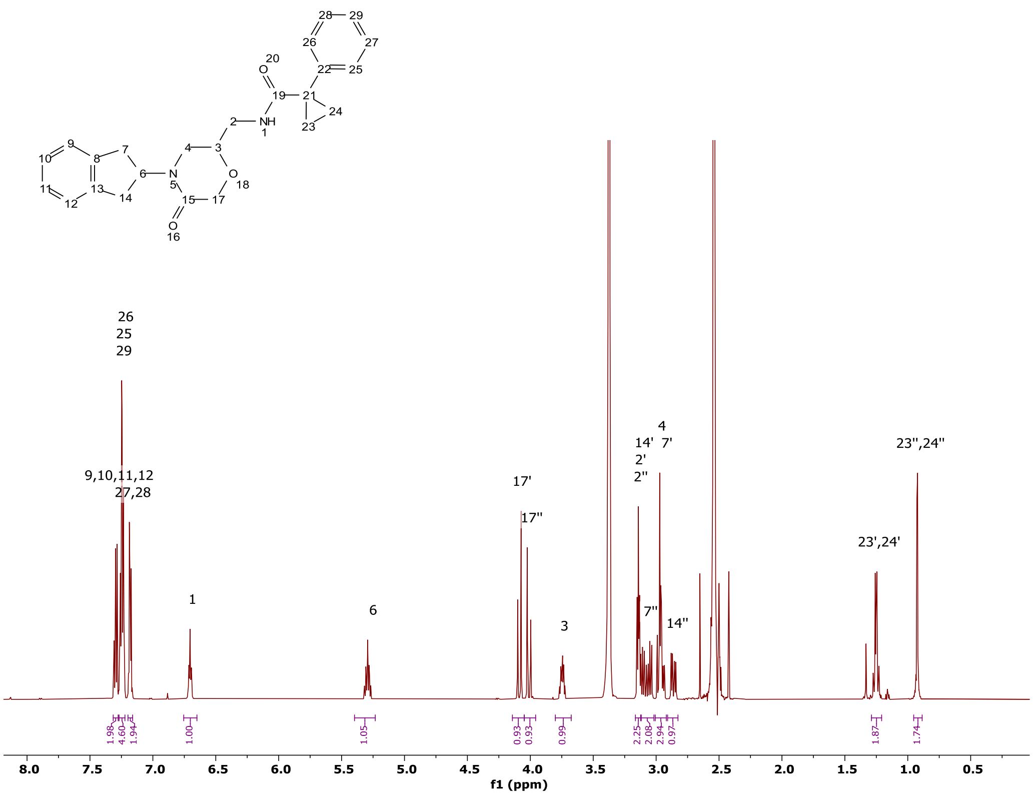

**Figure S21. 600.1 MHz $^{1}$H spectrum of OXM4 in DMSO-d$_6$.**

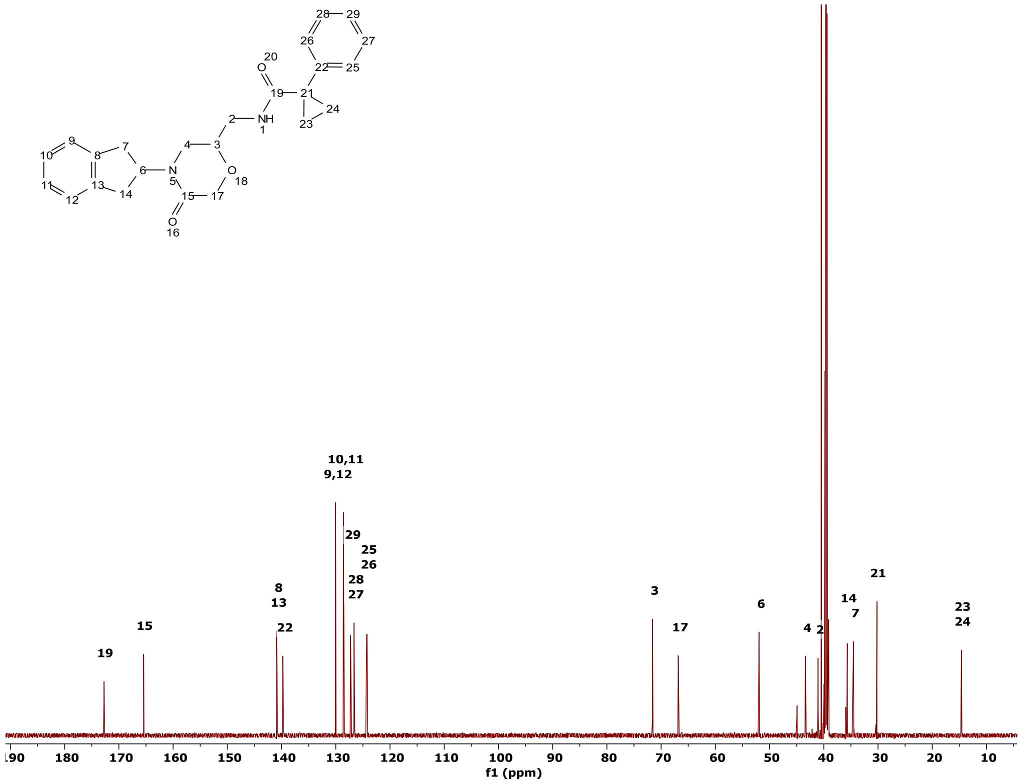

**Figure S22. 150.9 MHz $^{13}$C spectrum of OXM4 in DMSO-d$_6$.**

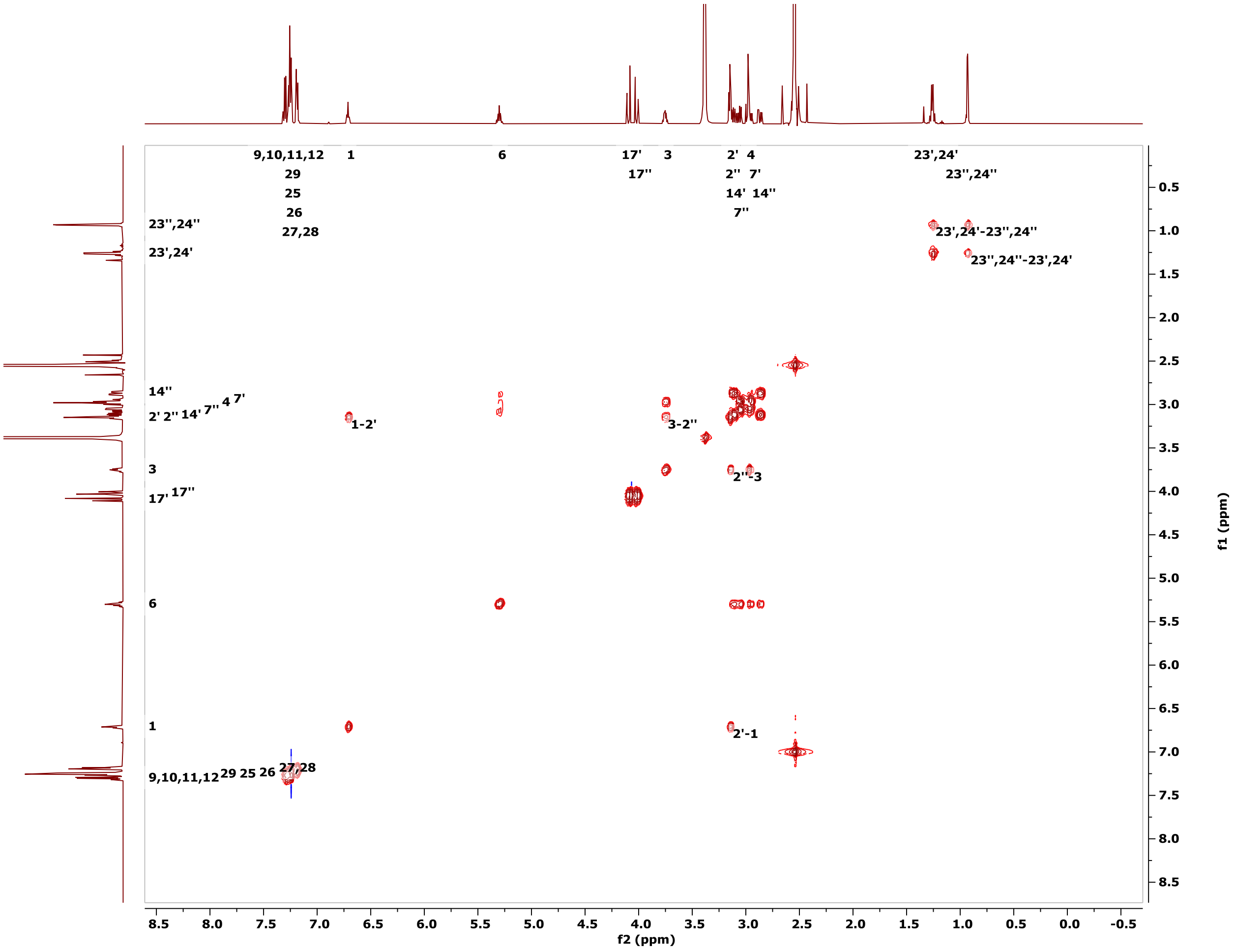

**Figure S23. 600.1 MHz COSY spectrum of OXM4 in DMSO-d6.**

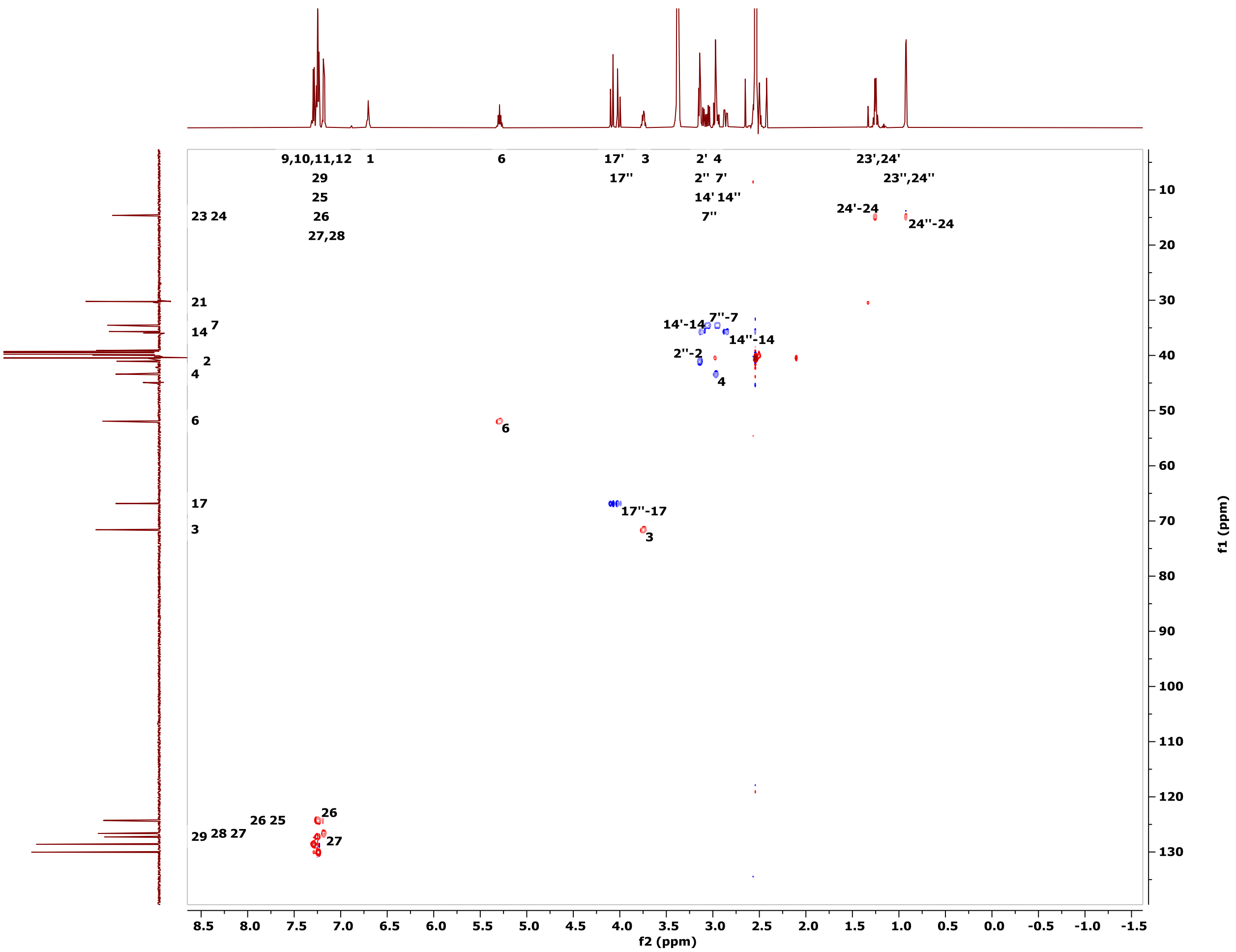

**Figure S24. 600.1 MHz HC-HSQC spectrum of OXM4 in DMSO-d6.**

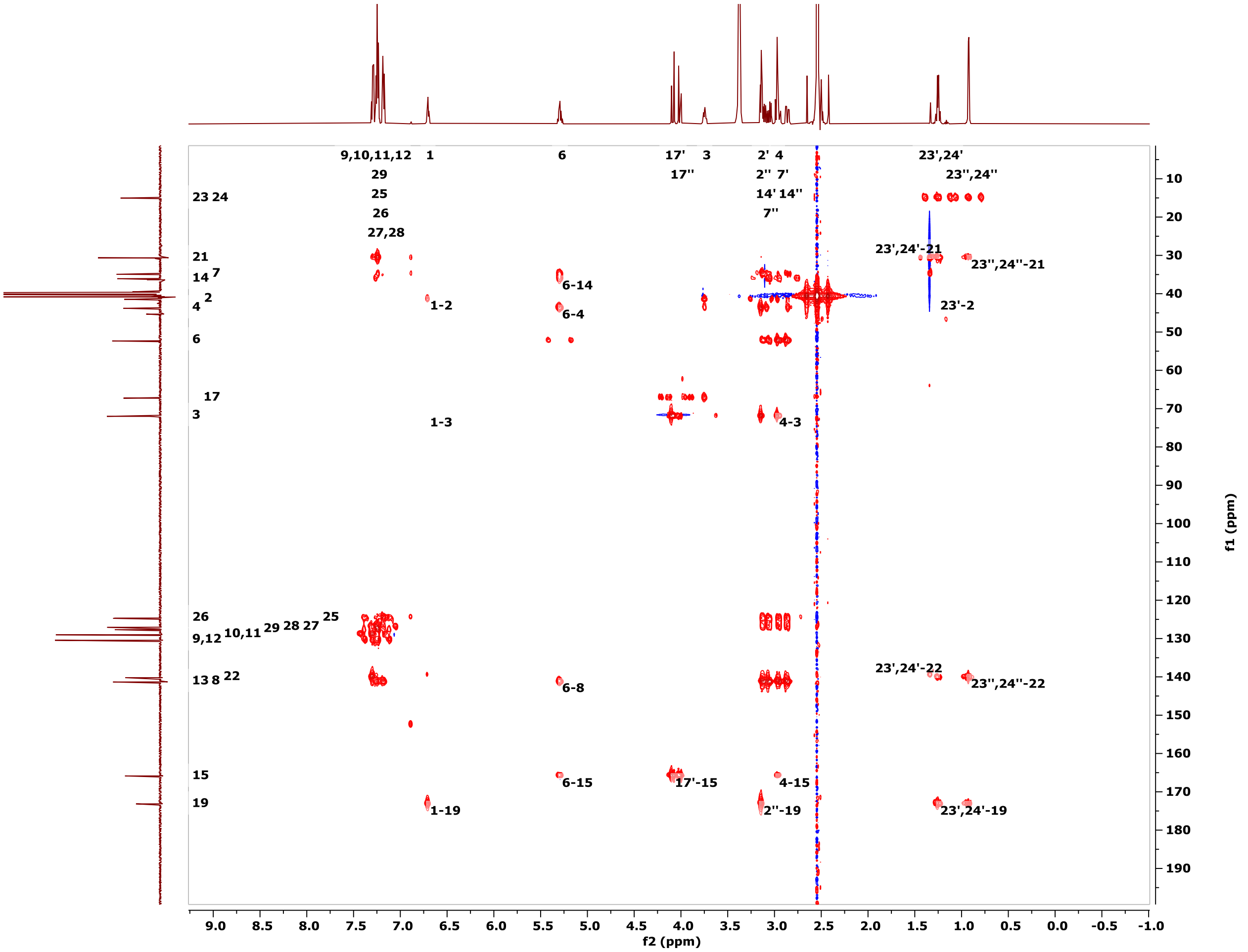

**Figure S25. 600.1 MHz HC-HMBC spectrum of OXM4 in DMSO-d6.**

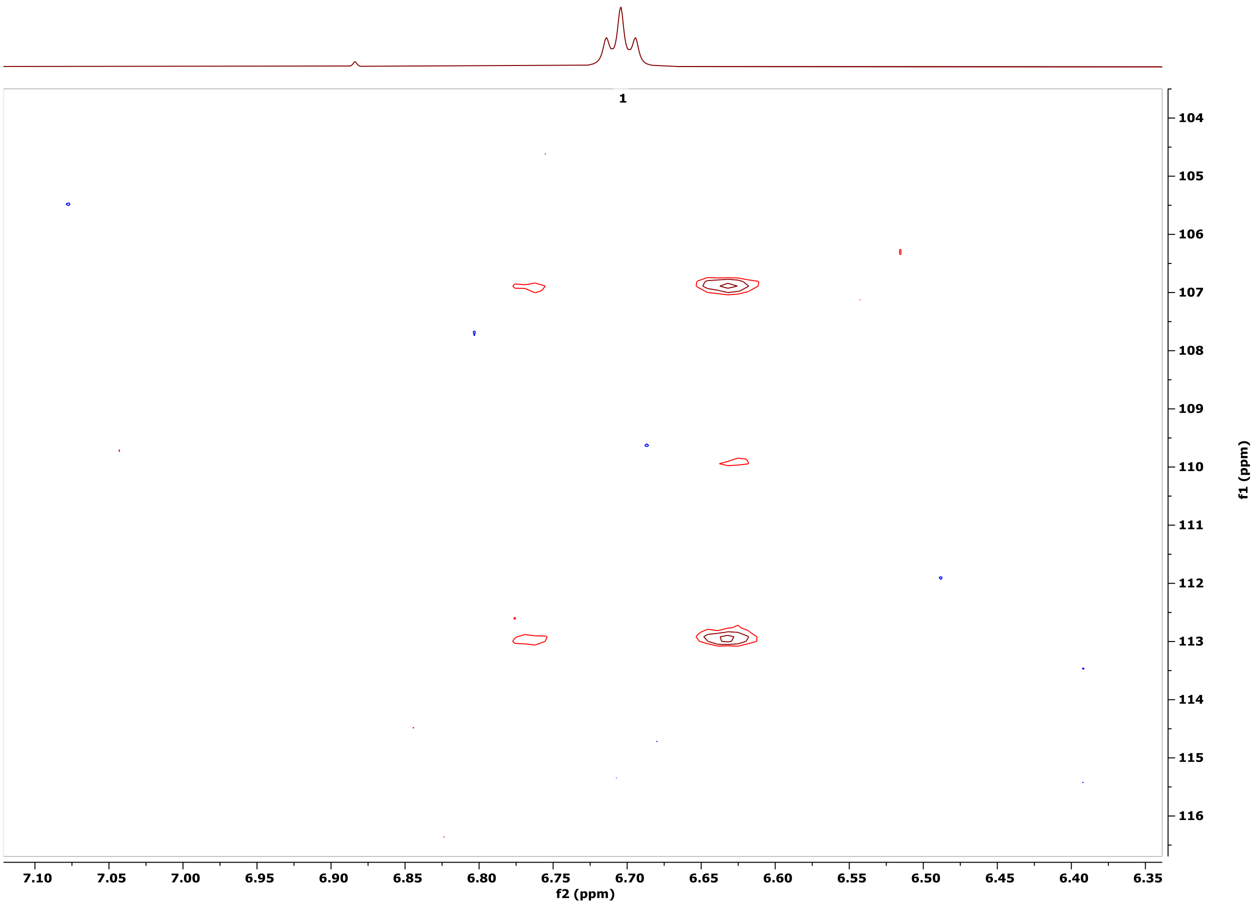

**Figure S26. 600.1 MHz JSBHN-HSQC spectrum of OXM4 in poly-HEMA gel in DMSO-$d_6$.**

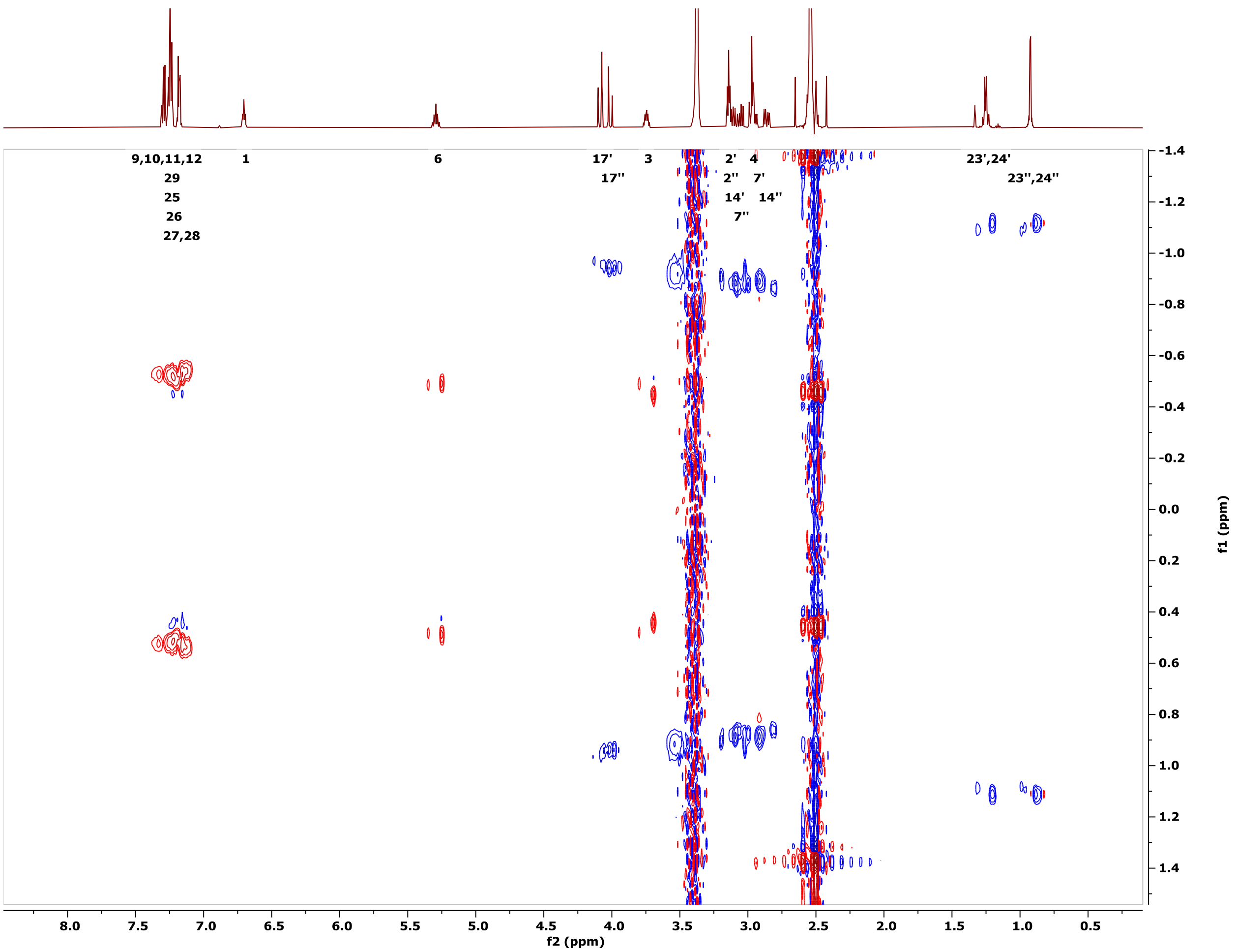

**Figure S27. 600.1 MHz JSBHC-HSQC spectrum of OXM4 in poly-HEMA gel in DMSO-d6.**

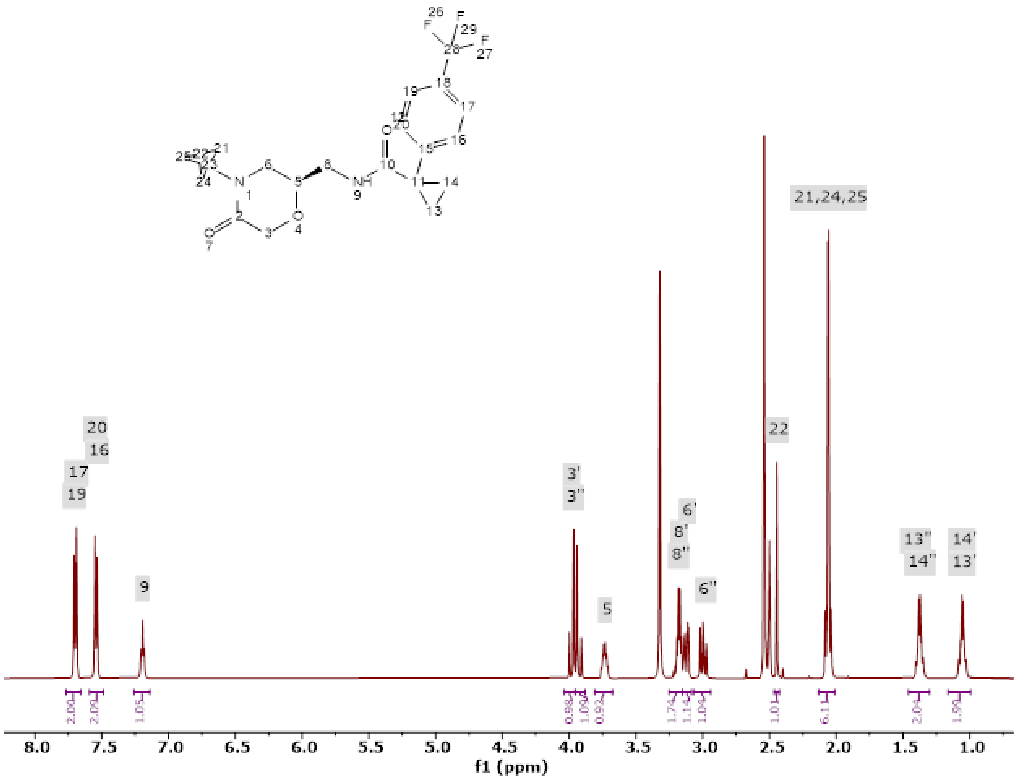

**Figure S28. 600.1 MHz $^1$H spectrum of OXM2 in DMSO-d$_6$.**

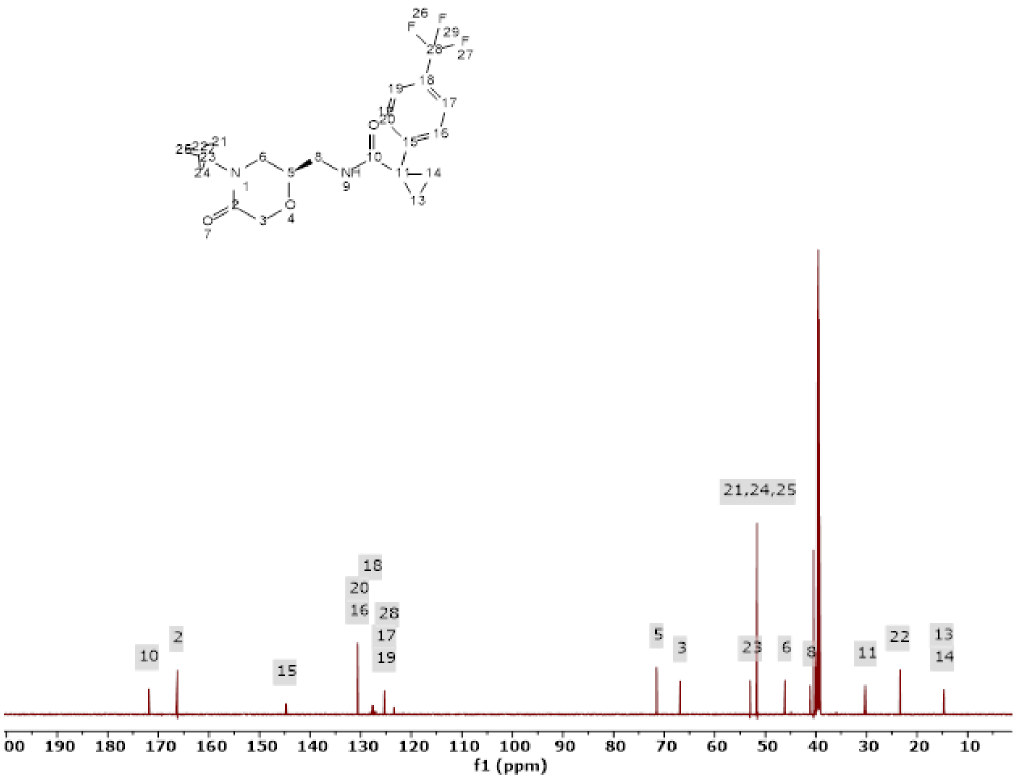

**Figure S29. 150.9 MHz $^{13}$C spectrum of OXM2 in DMSO-d$_6$.**

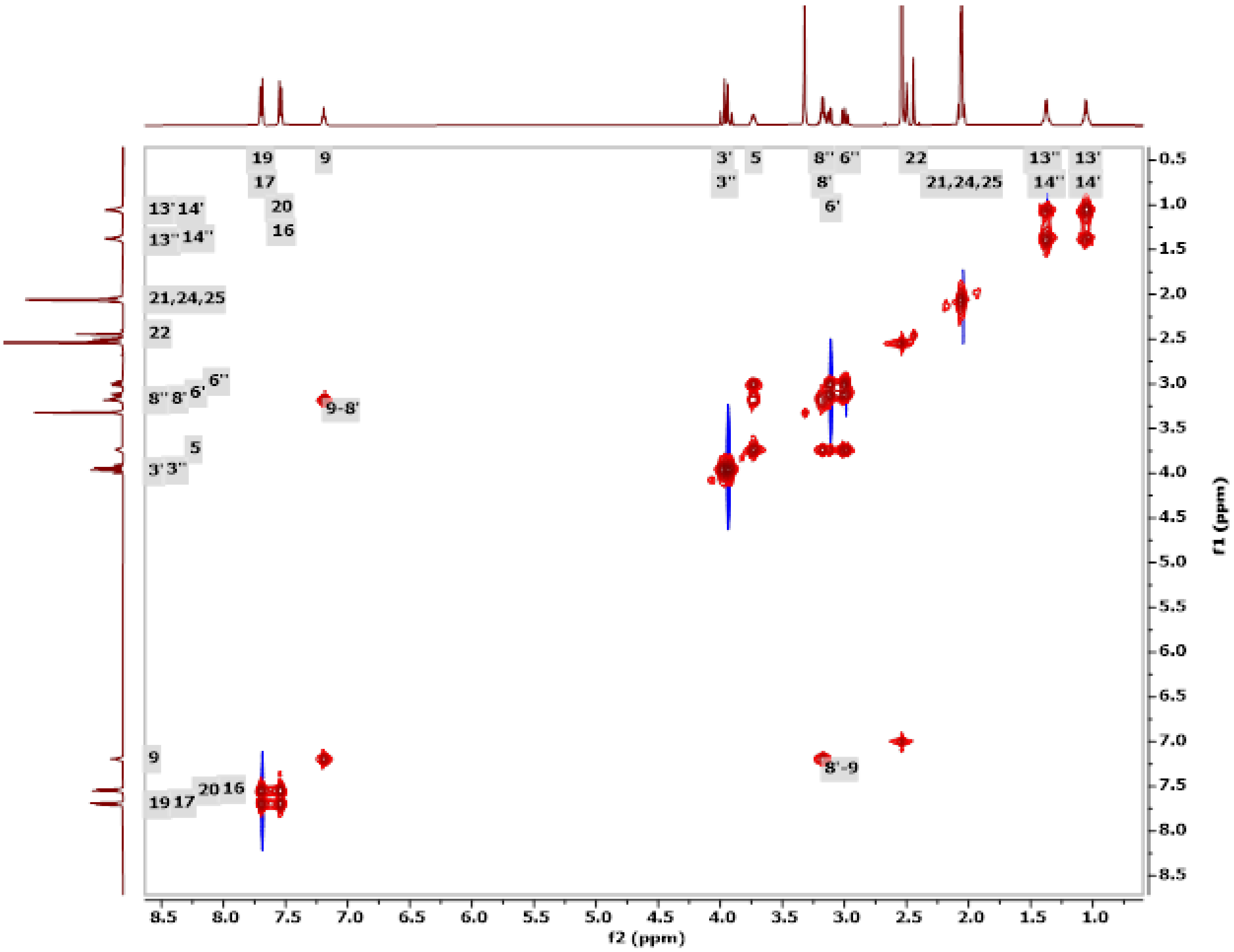

**Figure S30. 600.1 MHz COSY spectrum of OXM2 in DMSO-d6.**

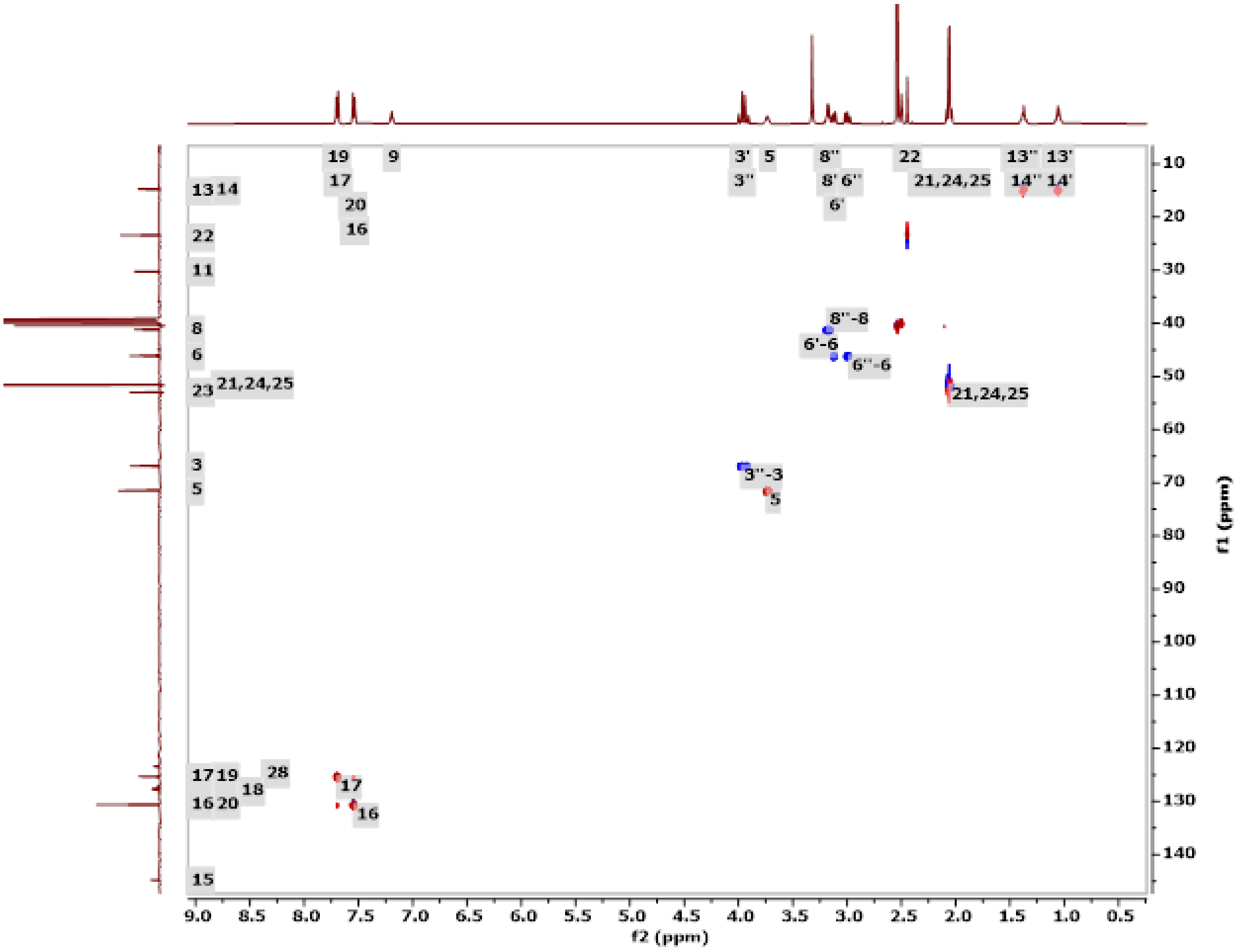

**Figure S31. 600.1 MHz HC-HSQC spectrum of OXM2 in DMSO-d6.**

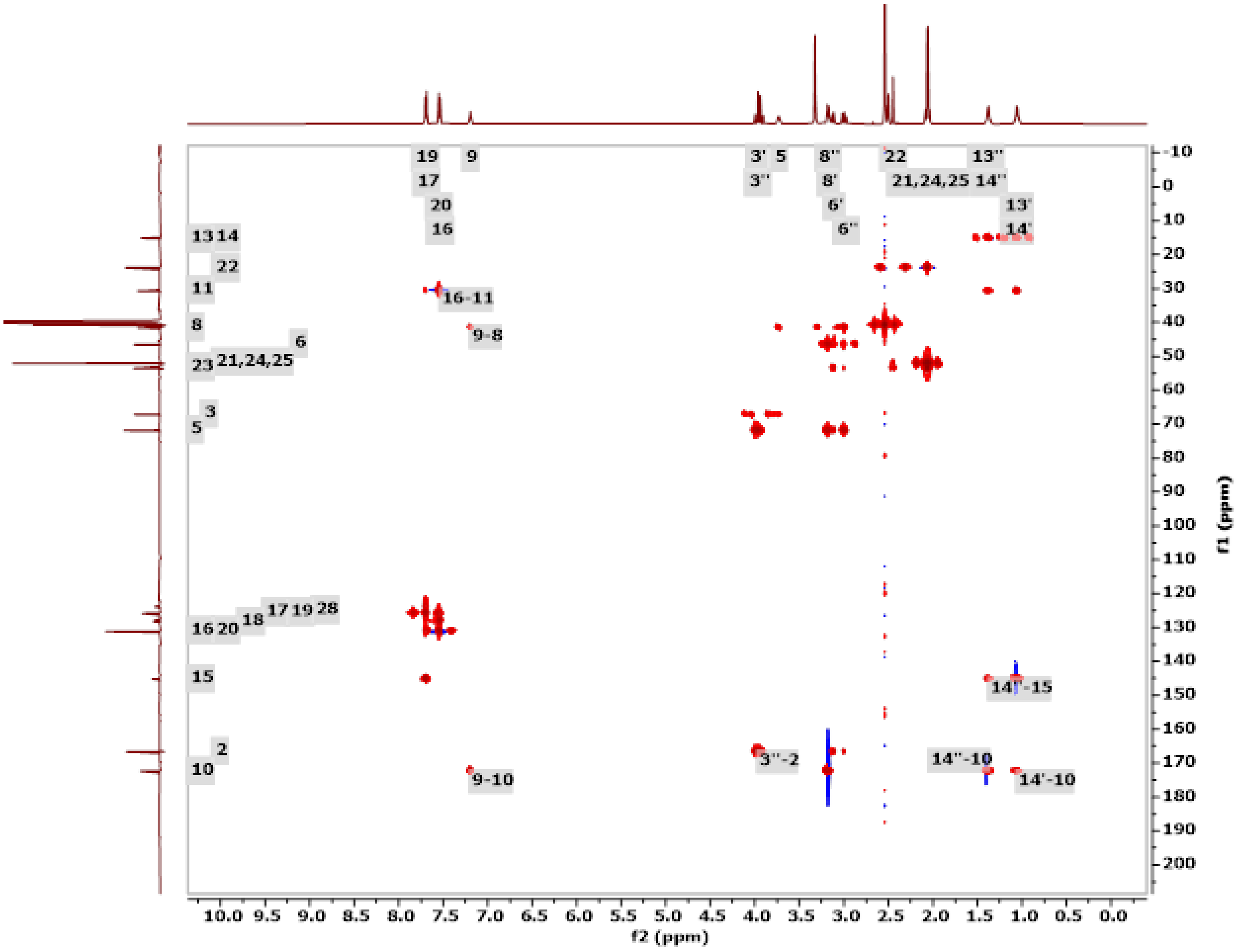

**Figure S32. 600.1 MHz HC-HMBC spectrum of OXM2 in DMSO-$d_6$.**

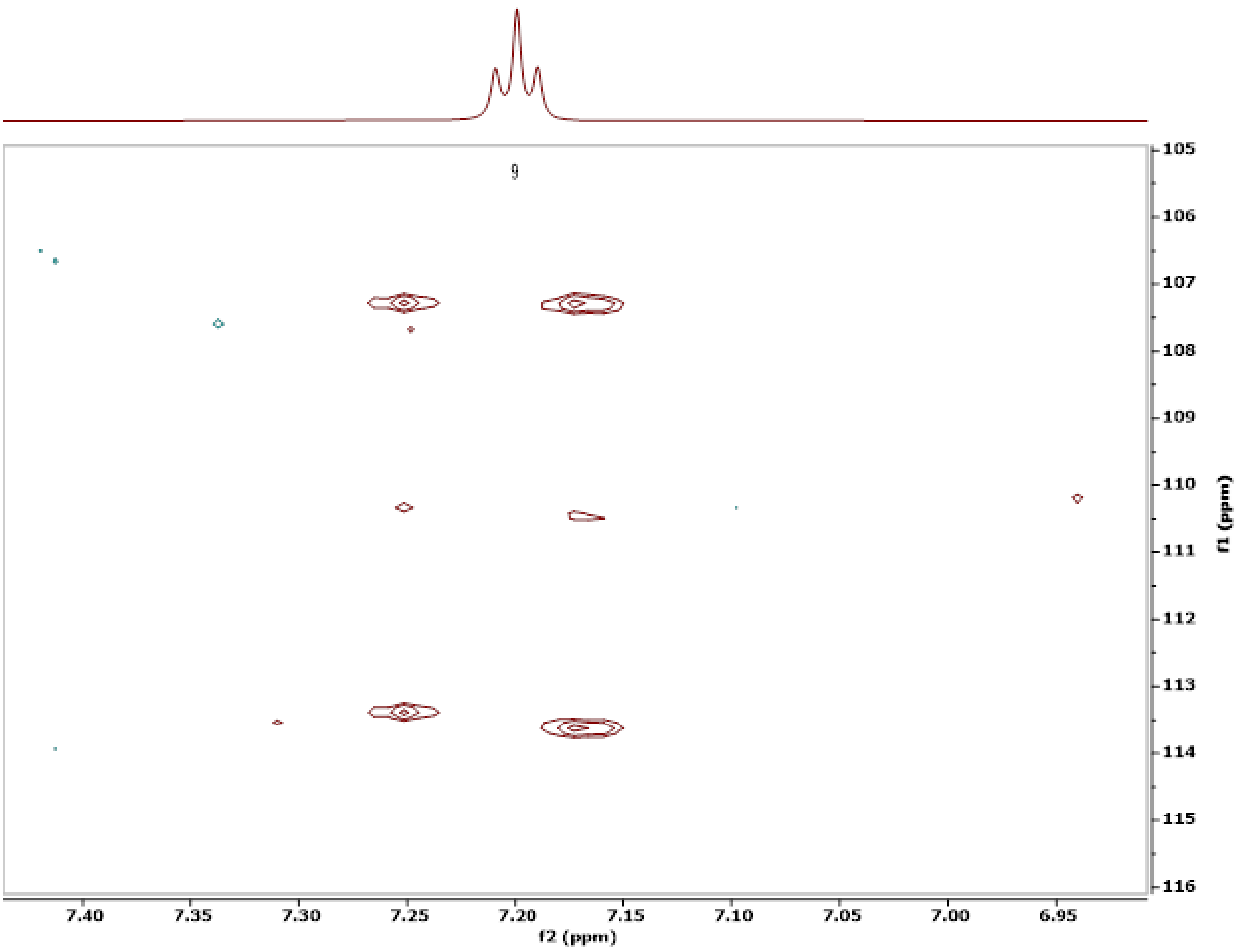

**Figure S33. 600.1 MHz JSBHN-HSQC spectrum of OXM2 in poly-HEMA gel in DMSO-d$_6$.**

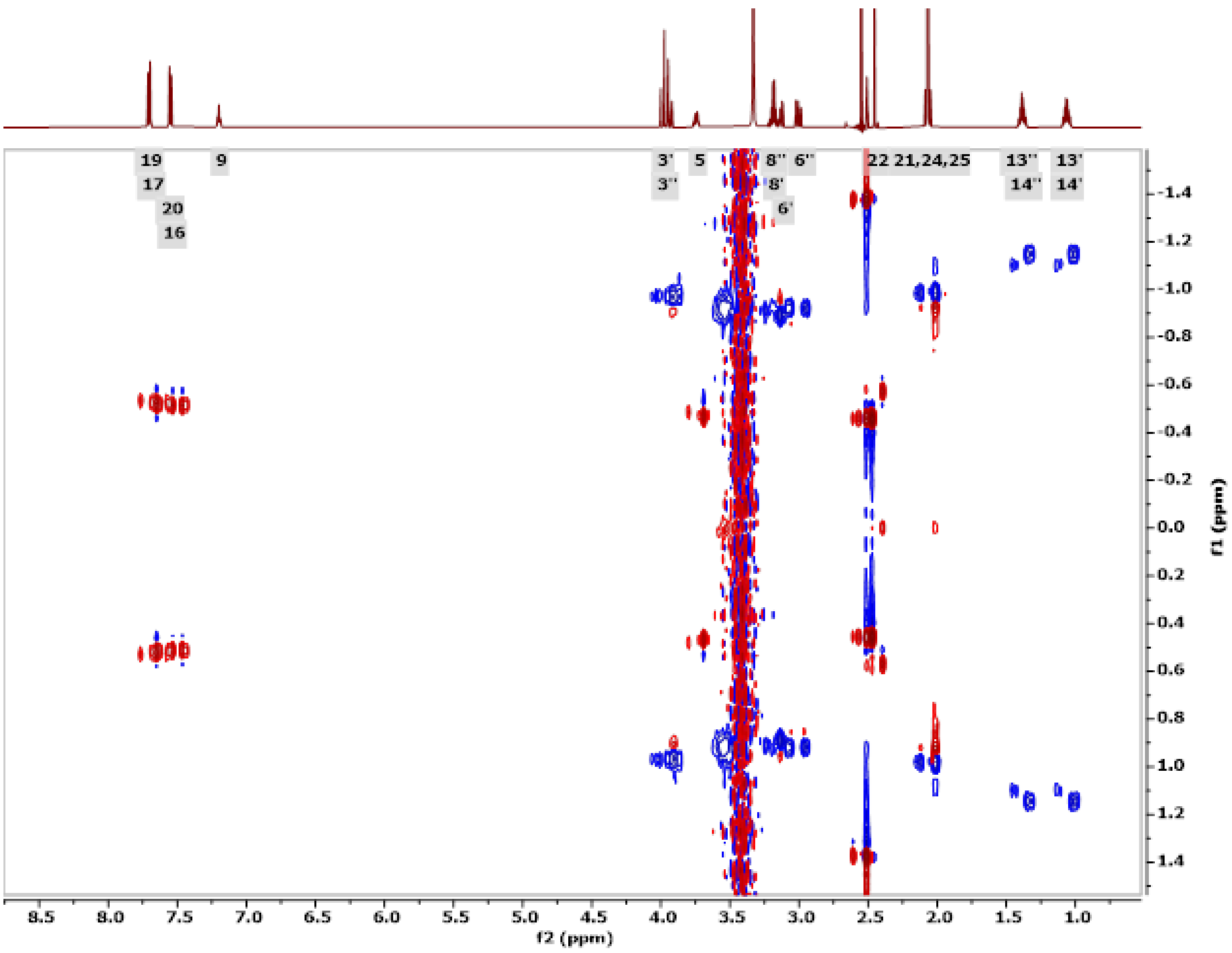

**Figure S34. 600.1 MHz JSBHC-HSQC spectrum of OXM2 in poly-HEMA gel in DMSO-d$_6$.**

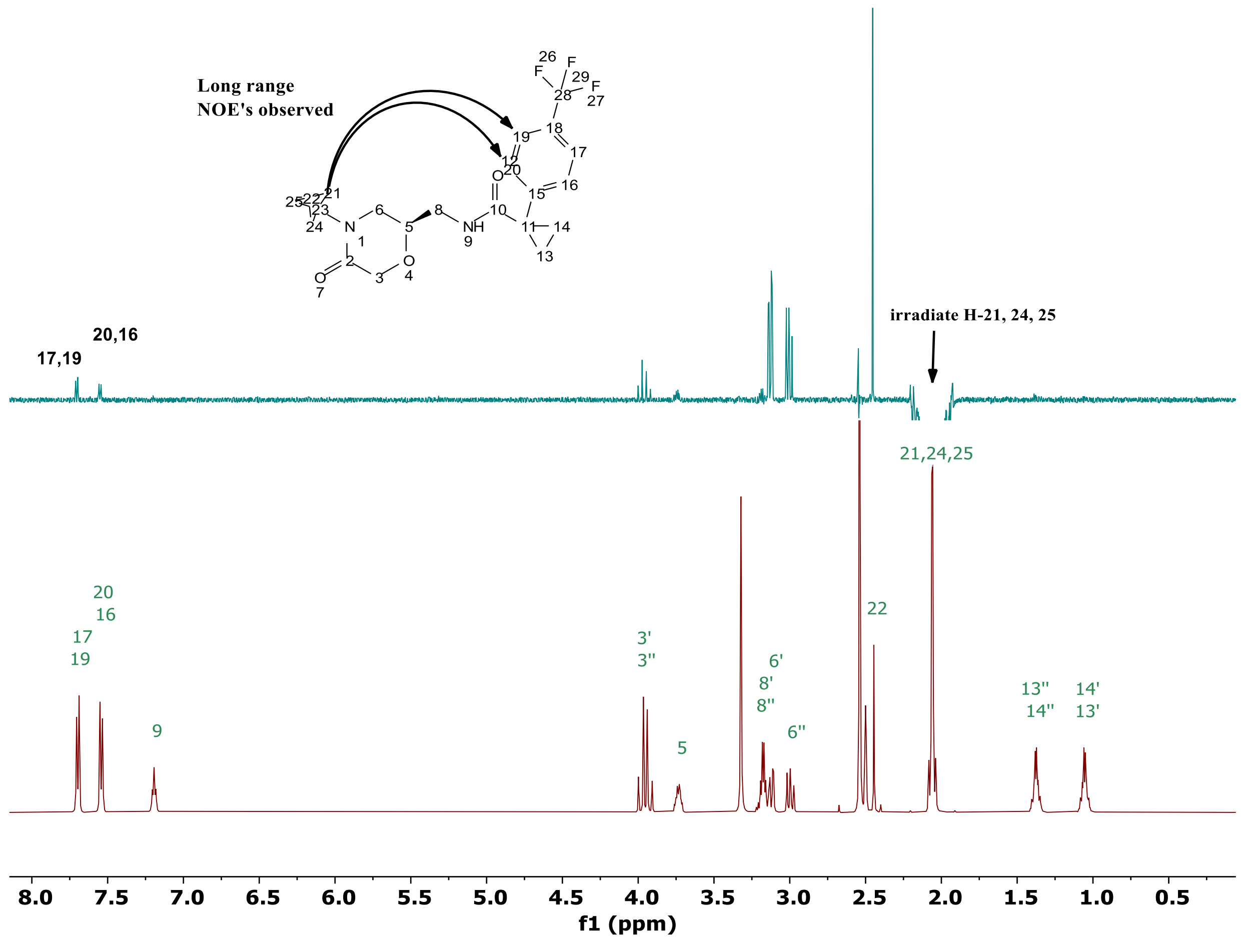

**Figure S35. Nuclear Overhauser Enhancement signals (NOEs) observed for OXM2 in DMSO-d$_6$.** The observation of long-range NOEs from H-21/24/25 to H-17/H-19 and to H20/H-16 suggests that OXM2 adopts a U-shaped conformation in solution.

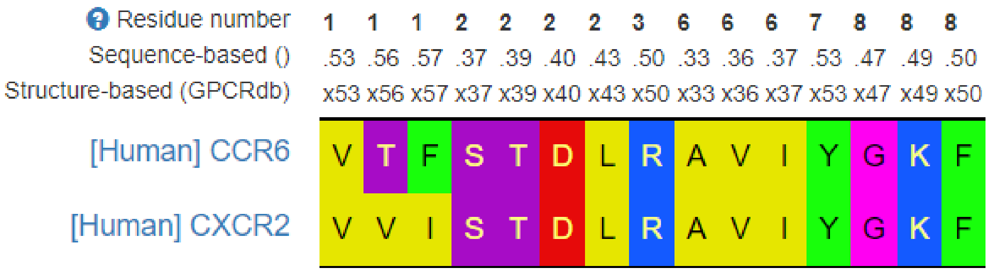

**Figure S36. Structure-based sequence alignment of human CXCR2 and human CCR6 at the SQA analogue SQA1 binding site.** The sequence alignment was performed using servers from www.GPCRdb.org[55].

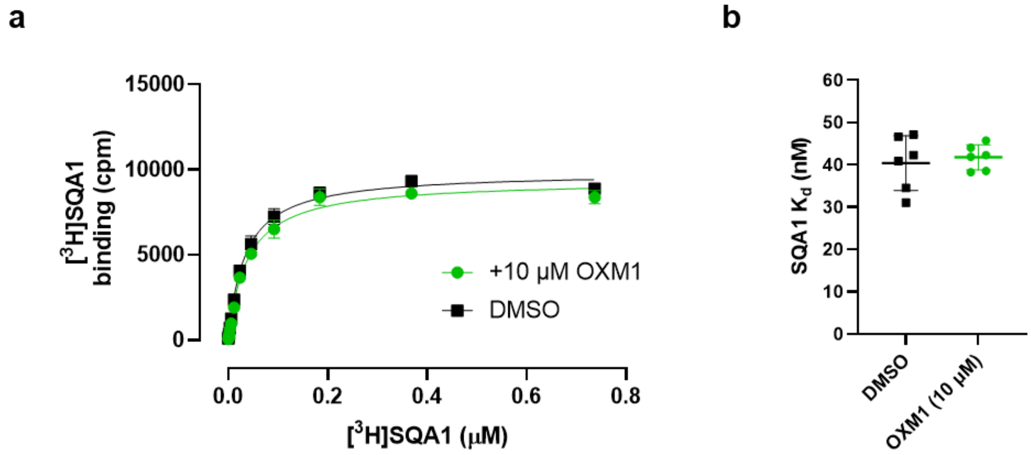

**Figure S37. Impact of OXM1 to SQA1 binding on CCR6. a**, Representative saturation binding curves of [$^3$H]-SQA1 to WT CCR6 with and without the presence of 10 μM of OXM1. **b**, Kd of SQA1 from the saturation binding experiment with and without the presence of OXM1. Average Kd and standard deviation presented in mean ± s.d. from *n* = 3 independent repeats are also shown.

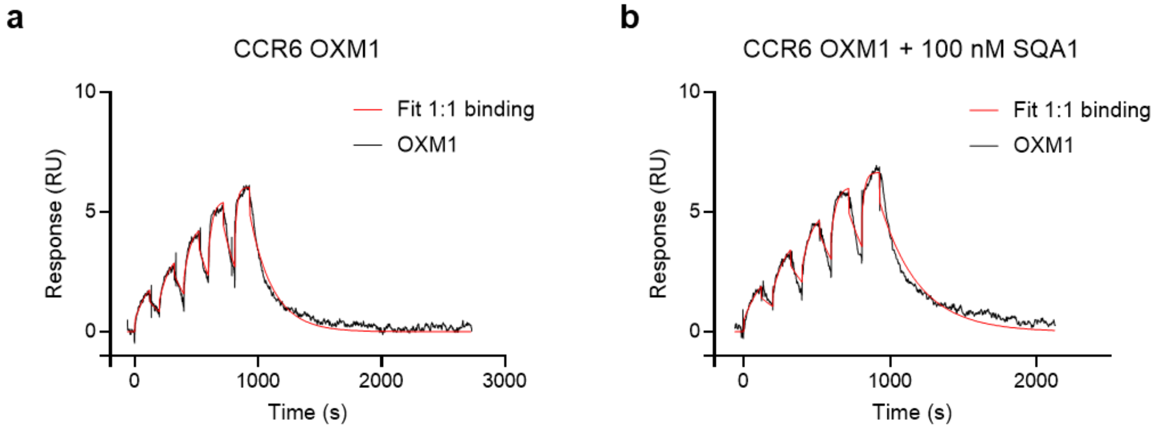

c

| Compound | Condition | Expt. | $k_a$ (1/Ms) | $k_d$ (1/s) | $K_D$ (nM) | Rmax (RU) | Comments |
|---|---|---|---|---|---|---|---|
| **OXM1** | **- SQA1** | 1 | 4.9E+04 | 5.4E-03 | 110 | 5.5 | Fig. S38a |
| | | 2 | 6.4E+04 | 6.1E-03 | 96.0 | 6.0 | |
| | | 3 | 4.3E+04 | 6.7E-03 | 156.7 | 6.3 | |
| | | **Average** | **5.2E+04 (±9.0E+03)** | **6.1E-03 (±5.4E-04)** | **121.107 (±25)** | | |
| **OXM1** | **+ SQA1** | 1 | 7.5E+04 | 4.2E-03 | 56.7 | 4.2 | |
| | | 2 | 6.9E+04 | 6.8E-03 | 97.9 | 5.7 | |
| | | 3 | 4.7E+04 | 3.8E-03 | 81.1 | 5.9 | Fig. S38b |
| | | **Average** | **6.4E+04 (±1.5E+04)** | **4.9E-03 (±1.6E-03)** | **78.6 (±20.7)** | | |

**Figure S38. Impact of SQA1 to OXM1 binding on CCR6. a**, Representative SPR sensorgrams of OXM1 binding to purified CCR6_Nβ10.2 protein. **b**, Representative SPR sensorgrams of OXM1 binding to purified CCR6_Nβ10.2 protein pre-incubated with 100 nM of SQA1. **c**, Table summary of experimantal conditions, maximal surface response (Rmax), binding kinetics ($k_a$, $k_b$), and Kd from each individual experiment and average reported in mean ± s.d. from $n$ = 3 independent repeats. In **a** and **b**, experimental data from the representative replicates as indicated in **c** are shown in black curves, fitting using a model of 1:1 binding are shown in red. Sensorgram images were prepared using GraphPad Prism.

### Supplementary Tables

**Table S1.** Cryo-EM data collection, refinement, and validation statistics.

| | CCR6/SQA1/OXM1 (EMDB-xxxx) (PDB xxxx) | CCR6/SQA1/OXM2 (EMDB-xxxx) (PDB xxxx) |
|---|---|---|
| **Data collection and processing** | | |
| Magnification | 215,000 | 130,000 |
| Voltage (kV) | 300 | 300 |
| Electron exposure (e–/Å$^2$) | 40 | 45 |
| Defocus range (μm) | -0.8 to -2.4 | -0.4 to -1.8 |
| Pixel size (Å) | 0.575 | 0.654 |
| Symmetry imposed | C1 | C1 |
| Initial particle images (no.) | 5,332,464 | 10,786,639 |
| Final particle images (no.) | 482,484 | 368,745 |
| Map resolution (Å) | 2.63 | 3.02 |
| FSC threshold | 0.143 | 0.143 |
| **Refinement** | | |
| Initial model used (PDB code) | Alphafold, 6WW2 | Alphafold, 6WW2 |
| Model resolution (Å) | 2.91 | 3.17 |
| FSC threshold | 0.5 | 0.5 |
| Map sharpening *B* factor (Å$^2$) | -92 | -112 |
| Model composition | | |
| Non-hydrogen atoms | 7403 | 7173 |
| Protein residues | 946 | 918 |
| *B* factors (Å$^2$) | | |
| Protein | 79.55 | 79.23 |
| Ligand | 110.12 | 109.95 |
| R.m.s. deviations | | |
| Bond lengths (Å) | 0.005 | 0.004 |
| Bond angles (°) | 0.945 | 0.854 |
| Validation | | |
| MolProbity score | 1.54 | 1.24 |
| Clashscore | 6.26 | 4.00 |
| Poor rotamers (%) | 0.0 | 0.0 |
| Ramachandran plot | | |
| Favored (%) | 96.78 | 97.78 |
| Allowed (%) | 3.22 | 2.22 |
| Disallowed (%) | 0.0 | 0.0 |

**Table S2.** OXM1 profiling against the chemokine receptor panel from DiscoveRx® in both agonist and antagonist modes. 1 μM of OXM1 was used in the profiling.

| GPCR ID | Assay Mode | Conc (μM) | Rep 1 RLU | Rep 2 RLU | Mean RLU | SD | %CV | % Inhibition |
|---|---|---|---|---|---|---|---|---|
| CCR1 | Antagonist | 1 | 1282680 | 1296960 | 1289820 | 10097 | 1% | 4% |
| CCR10 | Antagonist | 1 | 1303960 | 1318240 | 1311100 | 10097 | 1% | -4% |
| CCR2 | Antagonist | 1 | 2300480 | 2274440 | 2287460 | 18413 | 1% | -13% |
| CCR3 | Antagonist | 1 | 814520 | 780360 | 797440 | 24155 | 3% | -14% |
| CCR4 | Antagonist | 1 | 1175720 | 1201200 | 1188460 | 18017 | 2% | -2% |
| CCR5 | Antagonist | 1 | 916160 | 823760 | 869960 | 65337 | 8% | 2% |
| CCR6 | Antagonist | 1 | 543200 | 504000 | 523600 | 27719 | 5% | 51% |
| CCR7 | Antagonist | 1 | 3079160 | 3257240 | 3168200 | 125922 | 4% | 2% |
| CCR8 | Antagonist | 1 | 1331400 | 1292760 | 1312080 | 27323 | 2% | 1% |
| CCR9 | Antagonist | 1 | 874160 | 871640 | 872900 | 1782 | 0% | 2% |
| CMKLR1 | Antagonist | 1 | 1523760 | 1417360 | 1470560 | 75236 | 5% | -2% |
| CX3CR1 | Antagonist | 1 | 143360 | 147280 | 145320 | 2772 | 2% | 8% |
| CXCR1 | Antagonist | 1 | 2775080 | 3118360 | 2946720 | 242736 | 8% | 7% |
| CXCR2 | Antagonist | 1 | 964320 | 1045520 | 1004920 | 57417 | 6% | 0% |
| CXCR3 | Antagonist | 1 | 1285200 | 1311800 | 1298500 | 18809 | 1% | -4% |
| CXCR4 | Antagonist | 1 | 159880 | 150640 | 155260 | 6534 | 4% | 2% |
| CXCR5 | Antagonist | 1 | 1237600 | 1119440 | 1178520 | 83552 | 7% | -2% |
| CXCR6 | Antagonist | 1 | 107520 | 101080 | 104300 | 4554 | 4% | 13% |
| CXCR7 | Antagonist | 1 | 980840 | 1066520 | 1023680 | 60585 | 6% | 9% |
| XCR1 | Antagonist | 1 | 18480 | 19600 | 19040 | 792 | 4% | 13% |
| **GPCR ID** | **Assay Mode** | **Conc (μM)** | **Rep 1 RLU** | **Rep 2 RLU** | **Mean RLU** | **SD** | **%CV** | **% Activity** |
| CCR1 | Agonist | 1 | 679280 | 717080 | 698180 | 26729 | 4% | 4% |
| CCR10 | Agonist | 1 | 96600 | 96600 | 96600 | 0 | 0% | 0% |
| CCR2 | Agonist | 1 | 71120 | 95200 | 83160 | 17027 | 20% | 1% |

| | | | | | | | | |
|---|---|---|---|---|---|---|---|---|
| CCR3 | Agonist | 1 | 249760 | 244440 | 247100 | 3762 | 2% | 1% |
| CCR4 | Agonist | 1 | 174160 | 148680 | 161420 | 18017 | 11% | 1% |
| CCR5 | Agonist | 1 | 106400 | 111160 | 108780 | 3366 | 3% | 0% |
| CCR6 | Agonist | 1 | 67480 | 66920 | 67200 | 396 | 1% | -1% |
| CCR7 | Agonist | 1 | 788480 | 758240 | 773360 | 21383 | 3% | -1% |
| CCR8 | Agonist | 1 | 40880 | 43120 | 42000 | 1584 | 4% | 0% |
| CCR9 | Agonist | 1 | 153440 | 153160 | 153300 | 198 | 0% | 0% |
| CMKLR1 | Agonist | 1 | 101360 | 94640 | 98000 | 4752 | 5% | 0% |
| CX3CR1 | Agonist | 1 | 9240 | 10360 | 9800 | 792 | 8% | 0% |
| CXCR1 | Agonist | 1 | 212240 | 226240 | 219240 | 9899 | 5% | 0% |
| CXCR2 | Agonist | 1 | 190400 | 201600 | 196000 | 7920 | 4% | 1% |
| CXCR3 | Agonist | 1 | 509600 | 470960 | 490280 | 27323 | 6% | 2% |
| CXCR4 | Agonist | 1 | 90440 | 82600 | 86520 | 5544 | 6% | 9% |
| CXCR5 | Agonist | 1 | 230720 | 213920 | 222320 | 11879 | 5% | 1% |
| CXCR6 | Agonist | 1 | 24920 | 26320 | 25620 | 990 | 4% | 3% |
| CXCR7 | Agonist | 1 | 92120 | 107240 | 99680 | 10691 | 11% | 0% |
| XCR1 | Agonist | 1 | 6440 | 5880 | 6160 | 396 | 6% | -2% |

**Table S3.** Comparison of RDC data from OXM1, OXM3 and OXM4 in compressed poly-HEMA gels in DMSO-$d_6$ with calculated $^1D_{CH}$ and $^1D_{NH}$ couplings.

| Compound | Experimental NMR Data | | RDC-Refined Structure | |
|---|---|---|---|---|
| | $^1D_{CH}$ (Hz) in DMSO-$d_6$ | $^1D_{NH}$ (Hz) in DMSO-$d_6$ | Calculated $^1D_{CH}$ (Hz) | Calculated $^1$DNH (Hz) |
| **OXM1** | 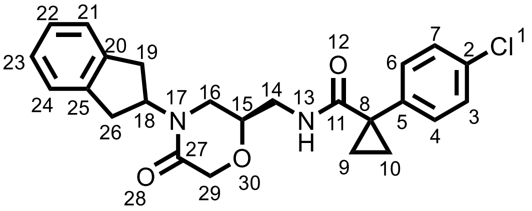 | | | |
| C22-H49 | -0.3 | - | -0.3 | - |
| C18-H45 | 0.6 | - | 0.6 | - |
| C15-H42 | -8.9 | - | -8.9 | - |
| C7-H34 | -6.1 | - | -6.1 | - |
| C29-H54/H55 | -3.0 | - | -3.0 | - |
| C26-H52/H53 | -5.5 | - | -5.5 | - |
| C10-H37/H38 | 6.3 | - | 6.3 | - |
| N13-H39 | - | 9.0 | - | 9.0 |
| **OXM2** | 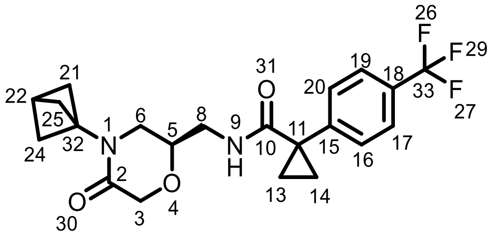 | | | |
| C14-H40/H41 | 5.8 | - | 6.1 | - |
| C22-H48 | -1.8 | - | -1.8 | - |
| C6-H34/H33 | 0.11 | - | 1.0 | - |
| C8-H35/H36 | -2.0 | - | -2.2 | - |
| C5-H32 | -3.6 | - | -3.6 | - |
| C3-H30H/31 | -0.02 | - | -0.46 | - |
| C20-H45 | -7.6 | - | -7.6 | - |
| C24-H49/H50 | -1.0 | - | -1.0 | - |
| N9-H37 | - | -4.4 | - | -4.4 |

| | | | | |
|---|---|---|---|---|
| **OXM3** | 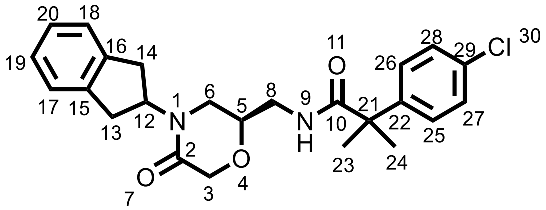 | | | |
| C5-H33 | -8.5 | - | -8.8 | - |
| C12-H39 | 1.0 | - | 1.6 | - |
| C27-H56 | 1.7 | - | 3.3 | - |
| C3-H31/H32 | -3.5 | - | -3.5 | - |
| C13-H40/H41 | 0.4 | - | 0.4 | - |
| C8-H36/H37 | -3.3 | - | -3.3 | - |
| C23-H48/H49/H50 | 0.7 | - | 0.6 | - |
| N9-H38 | - | 2.8 | - | 3.2 |
| **OXM4** | 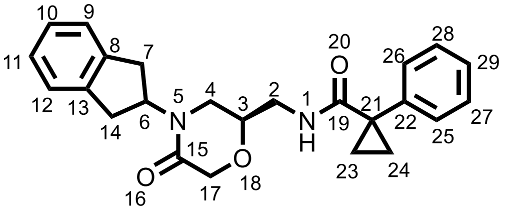 | | | |
| C6-H36 | 2.9 | - | 2.9 | - |
| C3-H33 | 13.3 | - | 13.3 | - |
| C27-H53 | 4.4 | - | 4.6 | - |
| C17-H45/H46 | -2.9 | - | -2.4 | - |
| C14-H43/H44 | 0 | - | 0.2 | - |
| C23-H47/H48 | 4.4 | - | 4.6 | - |
| N1-H30 | - | -0.7 | - | -0.7 |
